# Nanoneedle-Enabled Quantification of rAAV9 Capsid and Genome Integrity Reveals a Truncation Hotspot Locus in a 4.5 kb Transgene

**DOI:** 10.64898/2026.03.03.709319

**Authors:** Angad Garg, Elijah Litton, Tal Raz, Qimin Quan

## Abstract

**Background:** Adeno-associated virus (AAV) vectors are foundational to gene therapy but remain difficult to manufacture at high quality. Vector preparations frequently contain empty capsids and truncated genomes, diminishing potency and increasing immunogenic and production burdens. Conventional assays such as qPCR and ddPCR quantify only short regions, overestimating functional genomes and failing to resolve truncation patterns.

**Methods:** We applied the NanoMosaic Tessie nanoneedle platform to quantify AAV9 capsid and genome titers, directly distinguishing full-length (>4 kb) and truncated genomes. A 4.5 kb CAG– Luciferase–WPRE–bGH transgene packaged in AAV9 was analyzed using (i) nanoneedle “Probe Walk” assays to map truncations, (ii) PacBio SMRT long-read sequencing for orthogonal validation, and (iii) sedimentation velocity analytical ultracentrifugation (SV-AUC) to assess particle heterogeneity.

**Results:** Probe-walk mapping revealed asymmetric packaging with a ∼570 bp truncation hotspot 0.44–1.01 kb from the left inverted terminal repeat (ITR). PacBio sequencing confirmed positional concordance, identifying left partial reads clustering within the same region. SV-AUC resolved four major populations—empty (1.8%), partial (4.6%), full-length (70.4%), and high-molecular-weight (HMW) species (18.5%)—suggesting dimeric or multimeric capsids co-sedimenting with full-genome particles.

**Discussion and Conclusions:** The nanoneedle platform provided quantitative, region-specific insights into genome integrity that aligned with sequencing data while requiring minimal sample and processing time. The disproportion between molecular and AUC estimates indicates that apparent “full” species may contain long partial genomes or multimeric capsids bearing partial genomes. Together, these results establish the NanoMosaic Tessie system as a critical quality attribute (CQA) tool for assessing genome integrity and guiding process optimization. Integrating nanoneedle-based analytics early in development enables detection of truncation hotspots, improvement of vector fidelity, and acceleration of scalable, high-quality AAV manufacturing.

## Introduction

Adeno-associated virus (AAV) is a widely used gene therapy vector due to its favorable safety profile, ability to transduce both dividing and non-dividing cells, and potential for long-term gene expression (Wang J-H et al. 2024). However, manufacturing high-quality AAV vectors remains a significant challenge. Achieving both high yields and a high proportion of fully packaged functional genomes is hindered by the frequent generation of empty capsids and truncated genomes. These shortcomings reduce therapeutic potency, necessitate higher doses, and increase the risk of immune responses and production burden. Overcoming these barriers is critical to advancing safe, effective, and scalable AAV-based therapies.

A major source of heterogeneity in recombinant-AAV (rAAV) products is the formation of truncated or partial genomes. These typically result from DNA lesions during replication— induced by oxidative stress, cellular damage, or process-related fragmentation—and are repaired through non-homologous end joining (NHEJ), leading to the encapsidation of subgenomic or snapback particles (Junping Z. et al. 2022, 2024). The integrity of inverted terminal repeats (ITRs), which serve as both replication origins and packaging signals, also plays a central role; single-end mutations modestly elevate truncation rates, while dual-end mutations significantly increase genome heterogeneity (Savy A. et al. 2017, Chen Y. et al. 2024). Capsid size imposes an additional constraint, with genomes exceeding ∼5.2 kb consistently truncated at the 5^′^ end due to the 3^′^-to-5^′^ packaging directionality (Wu Z. et al. 2010, Li X. et al. 2023, Kapranov P. et al. 2012).

Beyond physical constraints, molecular contributors such as cryptic resolution sites within plasmid backbones—often located in antibiotic resistance elements—can mimic ITRs, triggering aberrant replication or encapsidation of non-vector DNA (Junping Z. et al. 2022). Additional sources of heterogeneity include recombination events, template switching, and replication stalling, all of which can generate truncations, internally deleted or chimeric genomes containing backbone–vector fusions (Kapranov P. et al. 2012, Tai PWL. et al. 2018). These mechanisms collectively explain the platform- and design-dependent variability in genome integrity observed across AAV products.

Importantly, downstream purification steps—while capable of enriching or depleting specific subpopulations—do not generate new truncations (Hashiba N. et al. 2025, Janc M. et al. 2024). Thus, it is critical to apply high-resolution integrity analysis during early manufacturing stages, particularly at the crude harvest and intermediate purification steps (e.g., iodixanol or CsCl gradients). Analyzing the distribution of genome species—including left- and right-end truncations, subgenomic fragments, and chimeric forms—enables manufacturers to (i) identify root causes of truncation (e.g., DNA damage, ITR instability, oversized constructs), (ii) assess platform and helper component effects, and (iii) optimize harvest timing and pooling of full-genome-enriched fractions.

Iterative testing at these stages allows process refinements—such as adjustments to transfection conditions, plasmid constructs, or cell stress mitigation—to be implemented upstream, before committing to full-scale production. As supported by recent studies, such targeted integrity assessment provides a mechanistically grounded framework to improve genome fidelity and overall vector quality (Junping Z. et al. 2022, 2023, 2024, Hashiba N. et al. 2025, Savy A. et al. 2017, Chen Y. et al. 2024).

Conventional methods like qPCR and ddPCR, while widely used, assess only short genomic regions (∼100–150 nt) and may substantially overestimate the proportion of functional genomes. To ensure therapeutic efficacy and regulatory compliance, there is a growing need for analytical tools that can distinguish between full-length, partial, and empty capsids—supporting both upstream development and final batch release.

To address this gap, the NanoMosaic Tessie platform offers a high-resolution, nanoneedle-based solution for assessing AAV genome integrity. Unlike qPCR-based approaches, NanoMosaic enables direct quantification of full-length transgenes (>4 kb), delivering an accurate measure of the therapeutic genome titer. With simple probe design, the platform can simultaneously quantify multiple genome regions, capturing both complete and truncated species.

Specifically, the full-length NanoMosaic assay detects intact therapeutic cassettes, while genome mapping assays generate quantitative, region-specific profiles of the AAV population. These assays reveal the molecular architecture of truncations and chimeric variants—allowing distinction between fully functional and aberrant particles.

In this study, we demonstrate the advantages of the NanoMosaic Tessie platform over ddPCR and AUC by analyzing a commercially sourced, purified AAV9 vector carrying a 4.5 kb transgene. We analyze the full-length and left- and right-end truncations and identify a truncation hotspot region likely associated with the vector sequence. The results highlight its value as a critical quality attribute (CQA) tool for guiding upstream decisions, monitoring genome integrity, and ensuring delivery of potent, high-fidelity gene therapy vectors.

## Materials and Methods

### Materials acquired

Sample was acquired from Vectorbuilder: Ultra-purified recombinant AAV9 virus, large-scale packaging, P240709-1025jfz pAAV[Exp]-CAG>Luciferase:WPRE (VB211007-1286euq). Catalog# P240709-1025jfz; Lot# 240726AAVN01. A certificate of analysis and the plasmid sequence with annotations is provided in Supplementary File S1.

### Nanoneedle assays

#### Viral genome Titer

Assays were performed according to the manufacturer’s instructions (NanoMosaic, AAV9 Viral-Genome Assay). Briefly: (i) MscI (NEB) digested plasmid DNA, that released a 4364-nt linearized transgene region serving as the DNA standard. This standard was quantified by Qubit BR and HS dsDNA kits as per manufacturers protocol. The rationale for MscI digestion and dilutions used for the standard curve are described in the text. (ii) Samples were processed as follows: The sample was treated with DNaseI (NEB) at 37°C for 30 min, then proteinase K (NEB) at 37°C for 30 min, and then incubated for 5 minutes at 95°C to release the viral DNA from capsids and inactivate proteinase K followed by slow cooling at 0.1°C/s to 10°C. Next, viral-DNA was MscI digested to remove the inverted terminal repeats (ITRs) and prevent any self-priming at 37°C for 30 min. (iii) Samples and standards were PCR amplified (NanoMosaic True Empty/Full/Partial Kit, AAV9 serotype) (see Supplementary File S2 - probed regions and primer sequences). (iv) nanoneedle plates were processed as follows: 1. plates were coated over-night with capture antibodies, 2. plates were blocked for 1 hour in blocking buffer and poly-T oligonucleotide, 3. the PCR amplified samples were hybridized to the plate for 1 hour at room temperature. 4. mass amplification reagents were added to the sample (SS2 for 30 min at room temperature, and SS3 for 15 min at room temperature per kit instructions). 5. Nanoneedle plates were then rinsed in DI water, quickly dried with pressurized air, and read on the Tessie instrument. The output data was analyzed using the NanoMosaic Montage software. Raw and analyzed data are provided in supplementary data (Supplementary File S3, Montage Raw and Processed Data).

#### Capsid Titer

Assays were performed according to the manufacturer’s capsid kit instructions (NanoMosaic True Empty/Full/Partial Kit, AAV9 serotype). Briefly: Nanoneedle plates were coated overnight with anti AAV9 intact particle mouse monoclonal capture antibody and then blocked. An 11-point standard curve was created (2-fold serial dilutions starting at 3.68E09) using empty AAV9 capsid particle standard. Samples were diluted to fall within the linear range of the standard curve, and six 2-fold serial dilutions were tested. Standards and samples were loaded in quadruplicates on the nanoneedle plate and incubated at room temperature for 1 hour. Biotin conjugated anti-AAV9 intact particle mouse monoclonal detection antibody was added and incubated at room temperature for 1 hour. Molecular mass amplification kit reagents were added (SS2 for 30 min at room temperature, and SS3 for 15 min at room temperature) and finally, plates were read on a Tessie instrument. The output data was analyzed using the NanoMosaic Montage software. Raw and analyzed data are provided in supplementary data (Supplementary File S3, Montage Raw and Processed Data).

### SMRT-sequencing and data analysis

PacBio long-read sequencing and data analysis was performed by SeqCenter (Pittsburgh, PA). Briefly, AAV DNA was extracted following the ThermoFisher Scientific PureLink Viral RNA/DNA mini kit (Thermo Fisher, 12280050) per PacBio recommendations. Samples were eluted into a final volume of 50 µl. Sequencing libraries were prepared following the ‘Preparing multiplexed AAV SMRTbell® libraries’ protocol, without thermal annealing, and using the SMRTbell® barcoded adapter plate 3.0 (Pacific Biosciences). Pooled libraries underwent the binding and cleanup steps using the Revio Polymerase kit (Pacific Biosciences).

Prepared libraries were loaded onto the PacBio Revio instrument. Demultiplexing, quality control, and adapter trimming were performed on instrument via Lima (PacBio barcode demultiplexer and primer remover). BAM files generated through HiFi sequencing were converted to FastQ files using the SamTools Fastq algorithm (Danecek P. et al. 2021). Generated reads from the PacBio instrument (Revio) were used as input for AAV Analysis using the Magdoll pipeline (https://github.com/Magdoll/AAV). Output PDF file is provided in supplementary data (Supplementary File S4, PacBio_AAV_report.

### Sedimentation Velocity Analytical Ultracentrifugation (SV-AUC)

The sample was thawed and equilibrated to room temperature. The absorbance (*A*_230_) of the sample was measured by Nanodrop. 420 µL of the sample at a concentration of 0.5 *A*_230_ units was loaded in a channel of an AUC cell with quartz windows and 12-mm double-sector Epon centerpiece. Sample was equilibrated at 20°C in the AUC chamber for 1 h (Beckman Coulter, Optima). Centrifugation was carried out at 20°C at 20,000 RPM. Scans were collected by measuring *A*_230_ and data sets were analyzed and fitted using UltraScan analysis software (Catalent, Baltimore, MD).

## Results

### The Nanoneedle technology

Traditional assays detect biomolecules by tagging analytes with fluorescent or absorbent reporters. While effective, these labels are poorly suited to wide-field, high-throughput imaging because they demand high-intensity illumination, long exposure times, and optics with high magnification and large numerical aperture. In complex sample matrices, reporter photophysics—such as blinking and photobleaching—also add variability to measurements.

The nanoneedle-platform replaces molecular labels with a solid-state optical transducer. Each nanoneedle is a cylindrical pillar patterned on the chip surface, typically 100–300 nm in diameter (Fig. 1C). Owing to their subwavelength dimensions and high refractive index material composition, these structures scatter light strongly. The optical response is highly sensitive to changes in the local refractive index at the nanoneedle surface. Once bound mass exceeds the detection threshold, the effective local refractive index—and, consequently, the scattering spectrum—shifts in proportion to the bound mass, enabling quantitative readout of molecular abundance (Fig. 1A, before assay; Fig. 1B, after assay).

**Figure 1:**
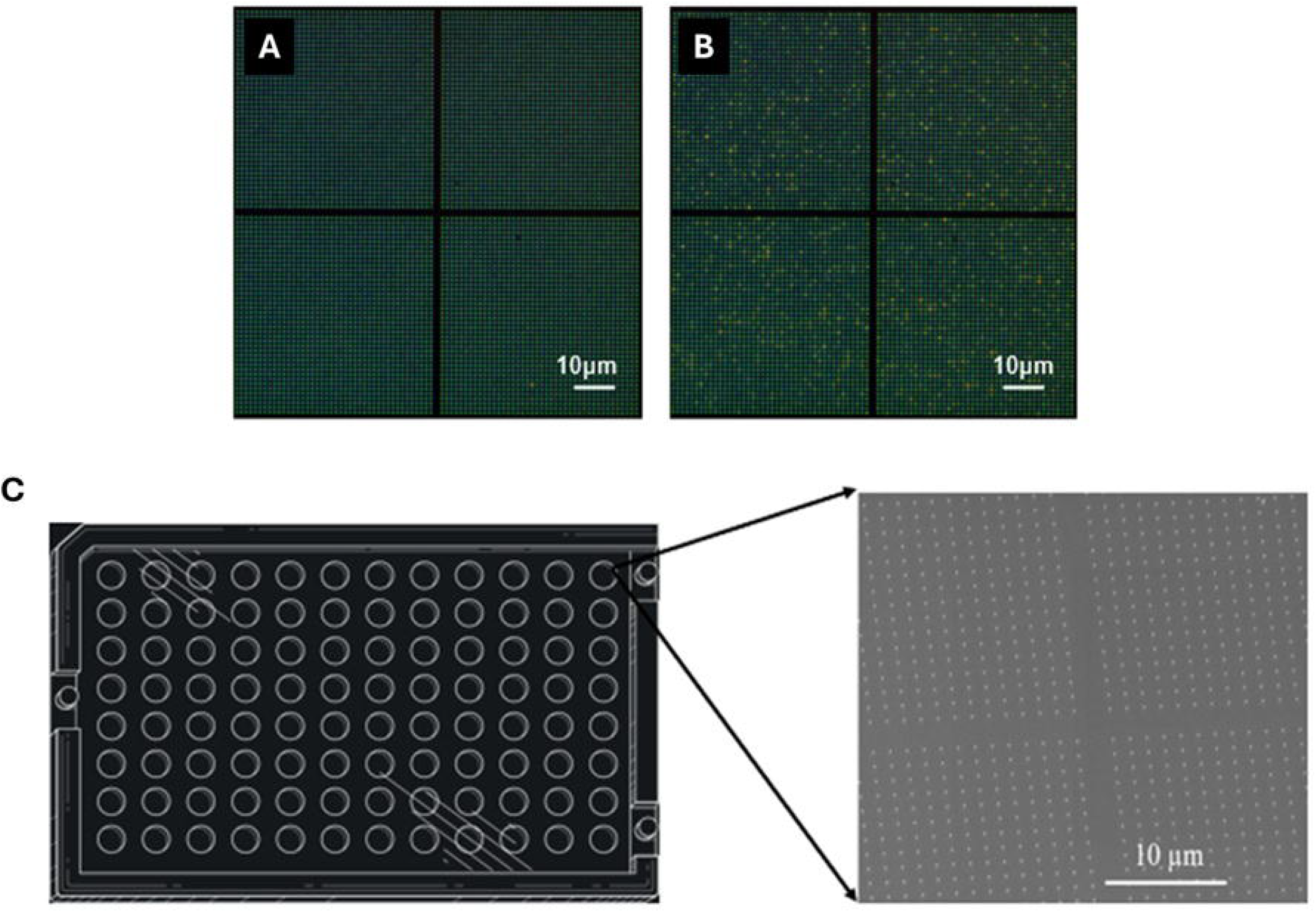
The Nanoneedle technology: **(A)** A portion of the nanoneedle sensor is shown with a light scattering pattern observed by the Tessie reader before the assay. **(B)** The same region shown in (A) after completion of the assay where a distribution of nanoneedles have a significant shift in mass to change the scattering spectrum. **(C)** A 96-well SBS nanoplate consumable is shown where each well has a 20,000 Nanoneedle sensor. The scanning electron microscopy image of the nanoneedles along with the scale bar is shown.

In the nano-plate consumable, densely packed nanoneedles are fabricated on a silicon chip. The nano-plate follows the Society for Biomolecular Screening (SBS) microplate standard to ensure compatibility with standard automated systems such as liquid handlers, where each well contains a sensor which is an array of ∼20,000 nanoneedles (Fig. 1C). The signal can be measured at each nanoneedle thus allowing high sensitivity due to the presence of multiple detection points. Each sensor can be imaged by dark-field microscopy in a single exposure with a color CMOS camera, allowing for a fast detection modality.

NanoMosaic’s nanoneedle platform was developed to quantify capsid and transgene titers. Workflow schematics for the capsid assay, transgene sample preparation, and transgene detection are shown in Figures 2, 3A, and 3B, respectively. In these workflows, biotin modification is used to append additional mass, thereby elevating the signal above the detection threshold.

**Figure 2:**
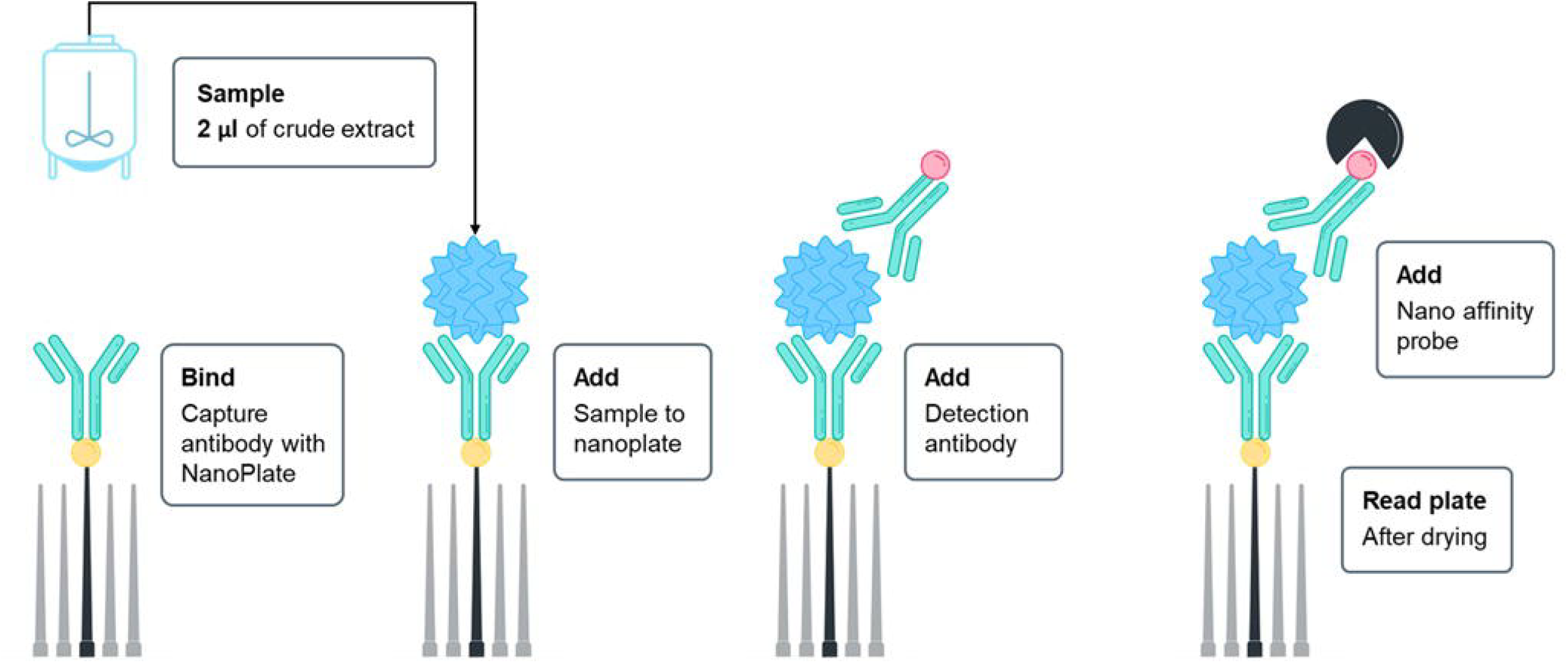
Nanoneedle capsid assay for AAV9 detection. Nanoneedles functionalized with an AAV9-specific antibody were used to capture AAV9 viral particles. A biotinylated AAV9 antibody was then applied, enabling mass addition through biotin anchoring. Capsid levels were quantified by measuring changes in light scattering before and after mass addition with respect to a standard curve.

**Figure 3:**
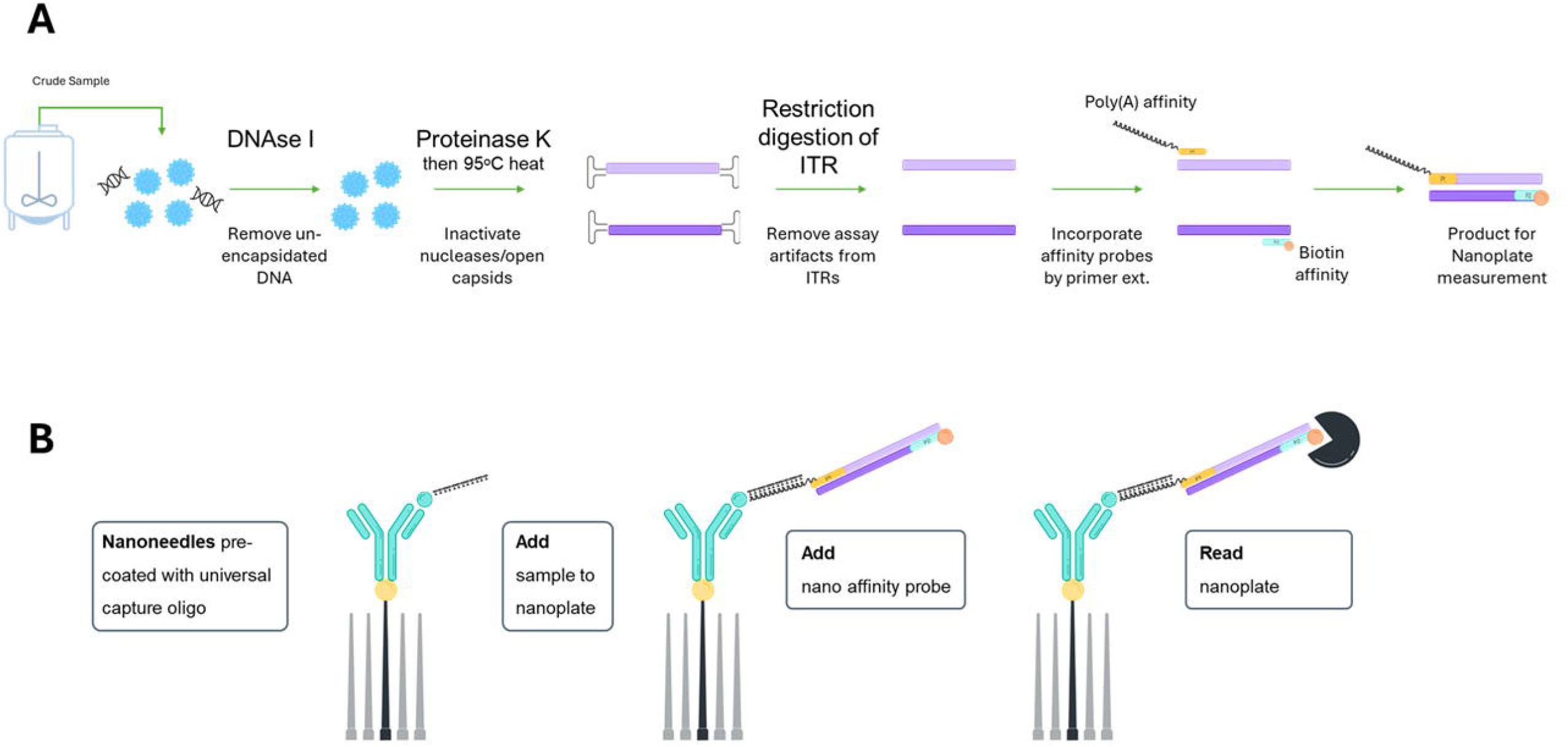
(A) Nanoneedle transgene assay sample preparation workflow. Samples were treated with DNase I to remove unencapsidated DNA, followed by Proteinase K digestion to degrade nucleases and initiate capsid disruption. Heat inactivation at 95°C deactivated Proteinase K and facilitated complete capsid opening and transgene release. Restriction digestion removed ITRs to prevent self-primed DNA synthesis artifacts (Brister JR et al., 2000). The transgene along with the DNA standard was then PCR-amplified using a forward primer with a 5^′^ poly(A) tail and a biotin-labeled reverse primer for nanoneedle-based detection. **(B) Nanoneedle transgene assay**. Nanoneedles were functionalized with a capture antibody to bind 3^′^-modified poly(T) oligonucleotides. Sample and standard amplicons hybridized to the poly(T) region, while the biotin label on the opposite end enabled mass addition for signal detection.

### “Probe Walk” AAV genome heterogeneity analysis

Recombinant AAV (rAAV) vector preparations often contain a heterogeneous mixture of genomes, including full-length, partially packaged, and truncated species (see Fig. 4). Accurate quantification of both full-length and subgenomic forms—including left- and right-end truncations—is essential for assessing product quality, ensuring potency, and meeting regulatory expectations for genome integrity. Primer-based assays targeting specific regions of the transgene provide a powerful means to resolve these variants, offering deeper insights into genome heterogeneity and informing AAV manufacturing optimization.

**Figure 4:**
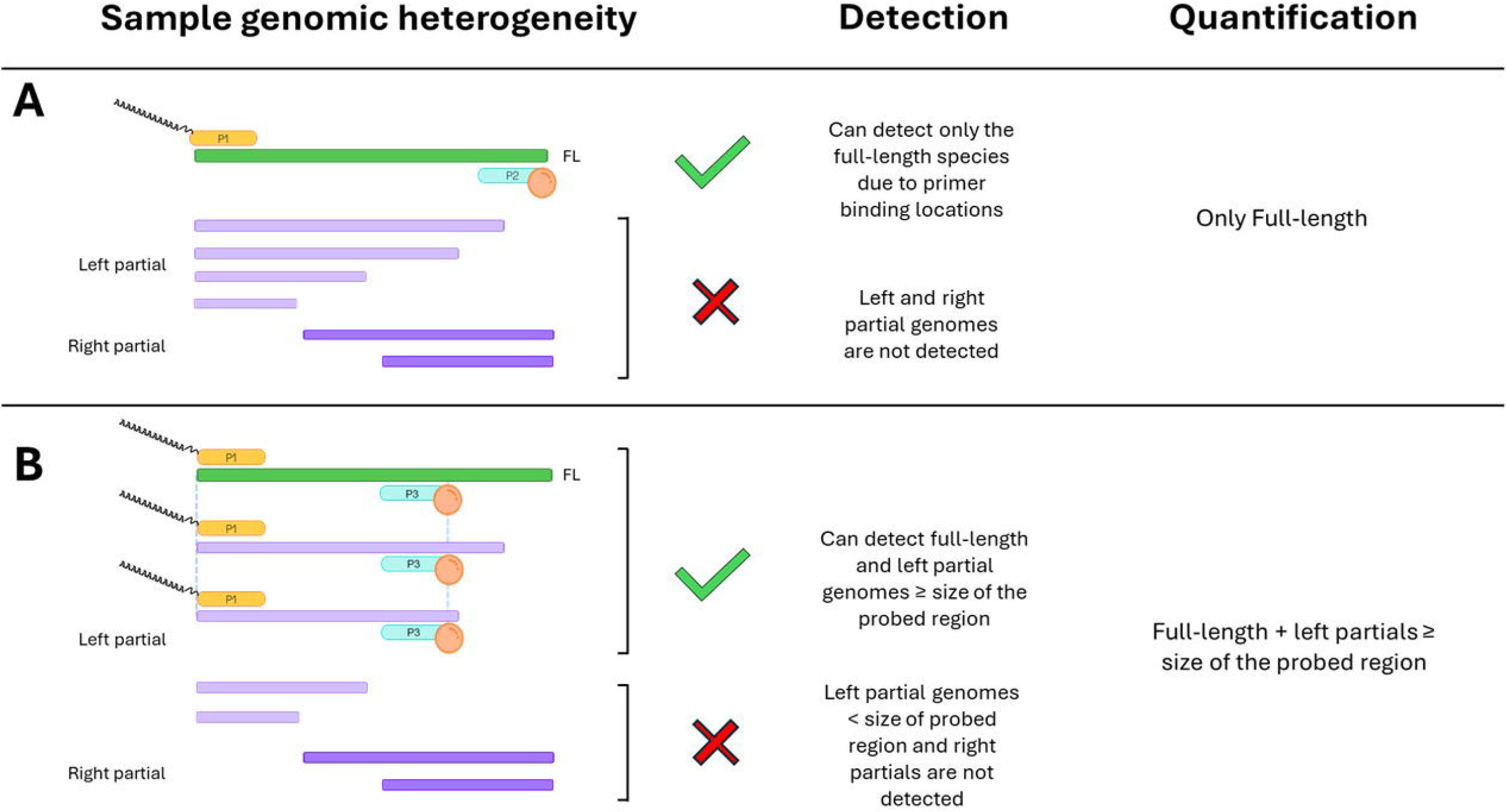
Nanoneedle “Probe Walk” primer placement strategy for detecting full-length and partial AAV genomes. The schematic depicts a hypothetical AAV population with full-length genomes (green bar), and left-(light purple bars) and right-(dark purple bars) partial genomes detected with: **(A)** Primers positioned at both ends of the transgene (P1 and P2) that selectively amplify full-length (FL) genomes; and **(B)** P1 primer paired with the P3 primer (positioned internally), which captures full-length and left-partial species ≥ the length of the probed region. Only genomes containing both primer-binding sites generate detectable amplicons.

To this end, we developed the “Probe Walk” methodology to map partial genomes and identify regions prone to truncation. The assay design and detection principle are illustrated in Figure 4. The full-length assay employs primers P1 and P2 at the 5’ and 3’ ends of the transgene, respectively, enabling amplification only from intact templates (Fig. 4A).

To detect left-partials, P1 remains at the 5’ end, while the reverse primer (P3) is placed internally. This configuration amplifies both full-length and partially packaged genomes that retain the region between P1 and P3, allowing detection of truncated species ≥ the probed length (Fig. 4B).

The schematic illustrates that only genomes containing both primer-binding sites yield amplicons (green check), while truncated templates missing either site fail to amplify (red cross). This flexible strategy can be extended by repositioning primers to probe different regions of the transgene and characterize truncation patterns more comprehensively.

We used this strategy to evaluate the truncation patterns in a CAG-Luciferase-WPRE-bGH p(A) transgene packaged rAAV9 vector purified by cesium chloride density gradient.

### Probe Walk analysis reveals a specific truncation profile within a packaged rAAV vector

To assess genome integrity within a 4.5 kb CAG–Luciferase–WPRE–bGH transgene packaged in AAV9, we applied the Probe Walk assay, using strategically positioned primers to quantify full-length [FL(I)], left-partials (L), right-partials (R), and both left and right-partial (LR) genome species (Fig. 5A, Supplementary File S2). Each probed region defines a distinct segment of the transgene, enabling quantitative resolution of truncation patterns. Quantitative outputs for each probed region are shown as a bar graph (Fig. 5B), with corresponding titers and precision of measurement (Fig. 5C). Linearity %CV indicates the variation in the back-calculated titer across independently measured dilutions, and recovery %CV of replicates represents the variation between technical repeats of the same dilution.

**Figure 5:**
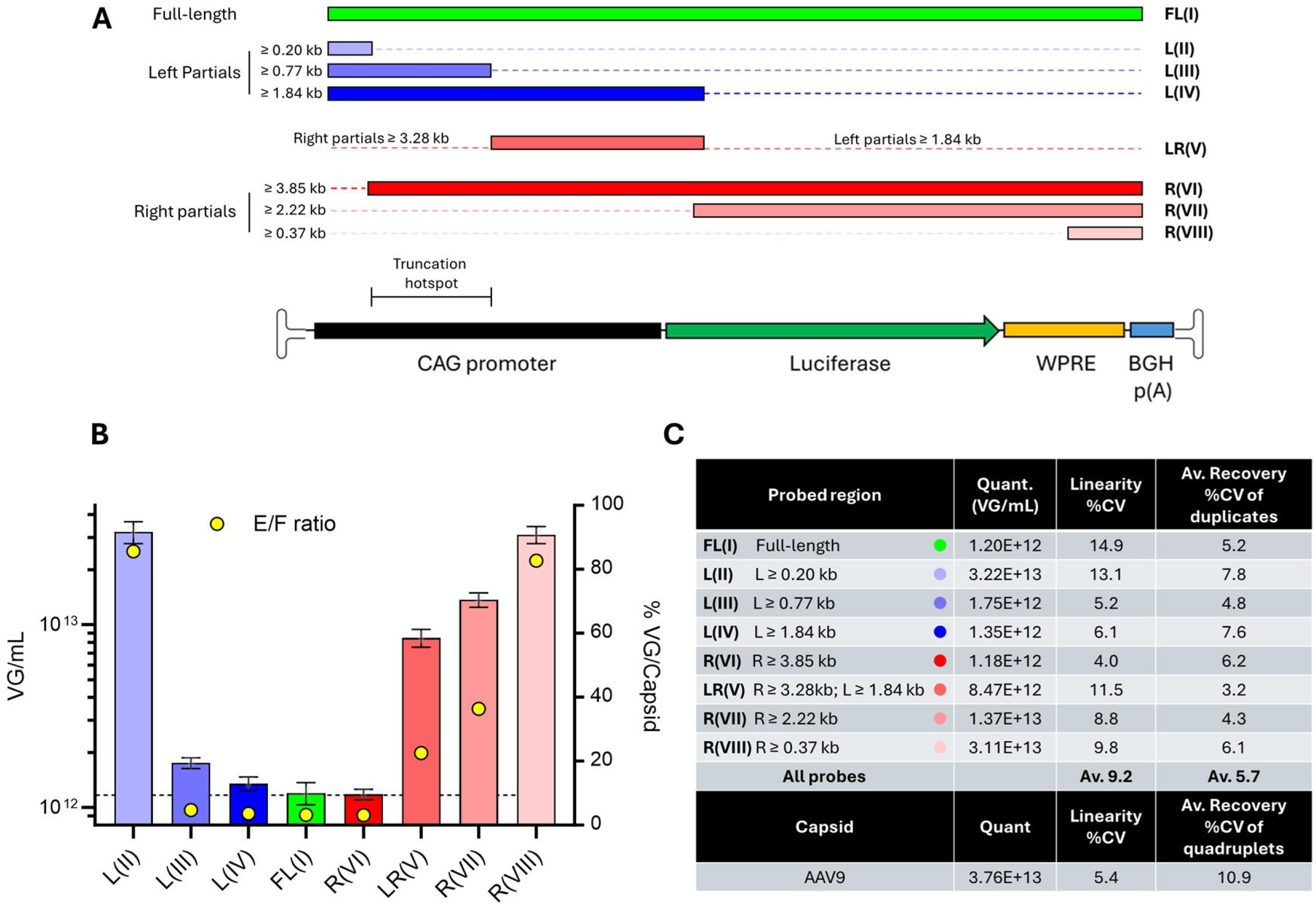
Probe Walk analysis across a packaged 4.5 kb CAG-Luciferase-WPRE-bGH transgene. **(A)** Schematic of the (+)-strand AAV ssDNA genome showing transgene elements and inverted terminal repeats (ITRs) depicted as folded structures. Color-coded bars indicate regions targeted in the Probe Walk assay, with each bar representing a specific primer pair designed to detect full-length (FL) genomes (green), left-partial (L, blue), right-partial (R, red), or combined left- and right-partial (LR) genomes. Color shading from light to dark reflects progressively larger probed regions, with the size range for each primer set indicated. A 0.57 kb region identified as a high-frequency truncation hotspot from the assay is marked above the transgene schematic. **(B)** The average quantified titers [VG/mL] for each probed region are shown as bars, with error bars representing ± S.D. across all dilutions and technical replicates. The Empty/Full ratio (% VG/capsid titer) is shown as yellow open circles. **(C)** Table showing titers, linearity coefficient of variation (%CV) based on 3–4 independent dilutions for viral genome titers and 5 independent dilutions for capsid titer, along with the average %CV across technical replicates for each viral genome region and capsid measurement.

To evaluate the distribution of left-partials, we held the forward primer near the left ITR constant and varied the reverse primer downstream from L(II) to L(III) to L(IV) (Fig. 5A). The titer decreased 18.4-fold between L(II) and L(III), indicating a high frequency of truncations between these two regions. Titer levels from L(III) to L(IV) and FL(I) decreased only modestly by 23% and 31%, respectively, suggesting that genomes extending beyond 0.77 kb were largely full-length species (Fig. 5B,C).

We then interrogated right-partials by fixing the reverse primer near the right ITR and varying the forward primer position, moving from R(VIII) to R(VII) to R(VI) (Fig. 5A). Titer declined 2.3-fold between R(VIII) and R(VII), suggesting a moderate frequency of right partials between these two regions. A further 11.6-fold drop was observed between R(VII) and R(VI), corresponding to the same region implicated in high-frequency left-partials [region between L(II) and L(III)]. Notably, the R(VI) and FL(I) titers were equivalent, indicating that there were no right-partials beyond 3.85 kb (Fig 5B,C).

To refine the location of the truncation “hotspot,” we examined **LR(V)**, capturing left-partial genomes ≥1.84 kb and right-partial genomes ≥3.28 kb. The majority of this signal likely originates from right-partial genomes, as left-partials beyond 1.84 kb [L(IV)] contributed minimally (Fig. 4B). A 7.1-fold decrease in titer from LR(V) to R(VI) further supports the presence of a high-frequency truncation region—approximately a 0.57 kb window—contributing disproportionately to genome loss (location is shown in Fig. 5A). This truncation hotspot region contained an internal 291-nucleotide 80% GC-rich region which also harbors homopolymer guanine nucleotide stretches. Inputting this region in mFold (Zuker M, 2003), we predicted strong secondary structures such as hairpins and G-quadruplexes. These secondary structures are known to impede viral DNA replication thereby leading to truncations (Xie J et al., 2017). Together, these data highlight asymmetric genome packaging and define a localized high-frequency truncation region within the AAV transgene.

To evaluate the empty/full ratio, we measured the capsid titer (Fig. 5C) and estimated the genome to capsid titer ratio for each region (Fig. 5B). The results echo the findings from the genomic titer that shows the asymmetric packaging favoring larger right-partials and shorter left-partials.

### Long-read sequencing

PacBio long-read sequencing was used as an orthogonal approach for partial genome analysis, with DNA libraries prepared after extraction without thermal annealing. A total of 158,590 reads were generated, of which 140,553 (87%) mapped to the transgene and, in part, to the plasmid backbone harboring the transgene. Among mapped reads, 60.6% were single-stranded AAV (ssAAV) and 39.4% were self-complementary AAV (scAAV). To align the long-read results with the probe-walk analysis, evaluation focused on the distribution of full-length and partial genomes mapping to the transgene. Overall, 55.4% of mapped reads were ssAAV full-length genomes, comprising 91% of all ssAAV genomes sequenced; left and right ssAAV partials accounted for only 3.2% of mapped reads, whereas left and right scAAV partials together represented 26.5%.

Triangulation of truncation sites showed that most left partials had read end points ∼250–1000 bp from the left start of the genome, peaking near ∼750 bp, concordant with the truncation hotspot identified by the probe walk analysis (0.44–1.01 kb from the left start). For right partials—where read orientation is reversed due to complementarity—reads initiated at the right ITR (Fig. 6; represented as read end position blue bar at ∼4500 coordinate) and exhibited end positions with higher frequency near the start of the genome, with additional termini distributed across the genome up to 3 kb from the right (Fig. 6; represented as blue bars across read start positions). This is consistent with decreasing titers when going from R(VIII) to R(VII) to LR(V) which shows a progressive loss in titers of longer regions for right-ITR initiated genomes (Fig.5).

**Figure 6:**
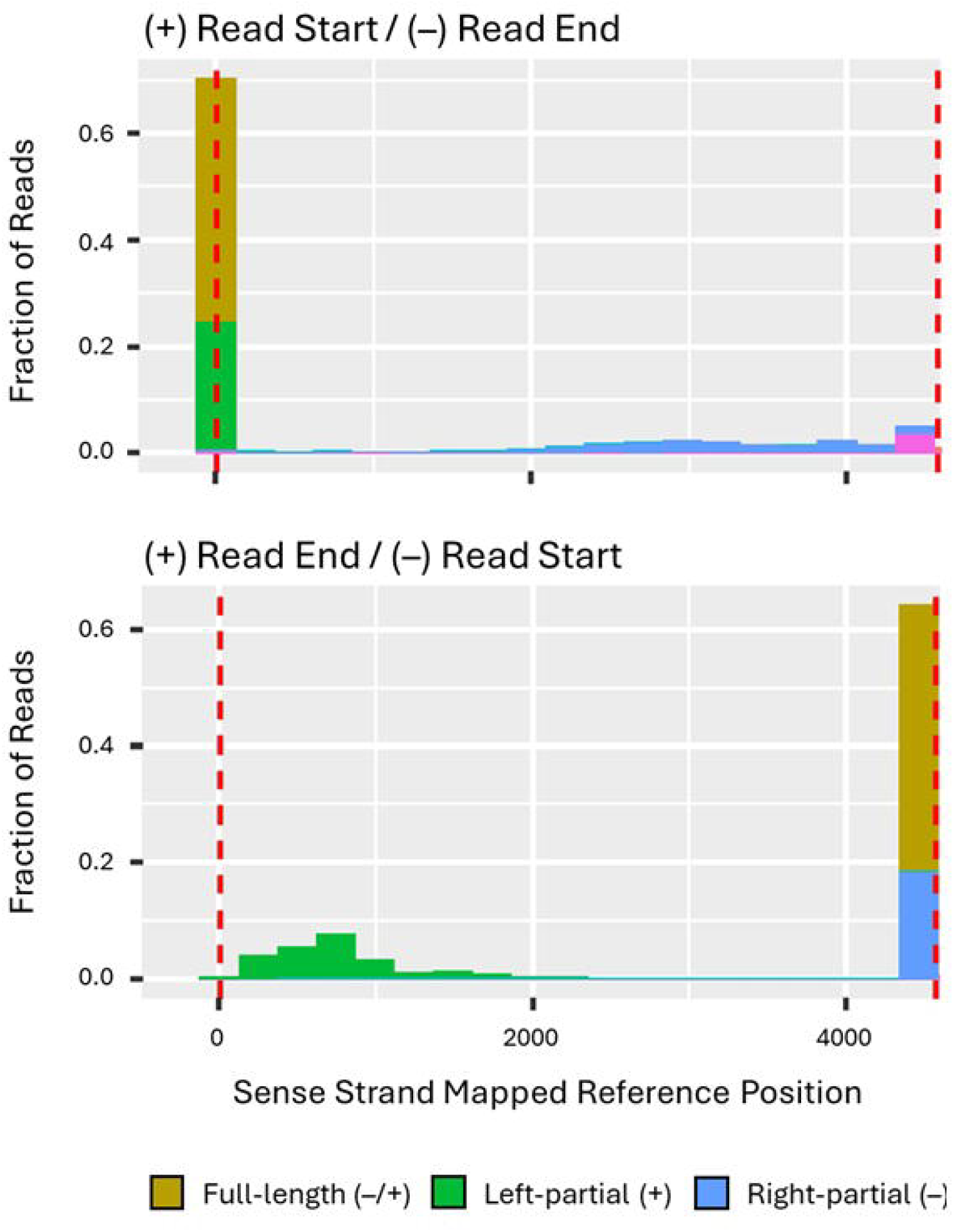
PacBio long-read sequencing reveals regions for truncations. The fraction of PacBio sequencing reads (y-axis) mapping to the transgene are plotted as a function of their start and end locations. The reads were mapped to the sense strand, and the top graph represents the read start location of the (+) strand and read end locations of the (–) strand genomes. The bottom graph represents the read end location of the (+) strand and read start locations of the (–) strand genomes. The fractions and the strand orientation of the full-length, left partials and right partials are indicated. (The pink bars indicate vector + backbone reads which were not considered in our analysis).

The sharp drop in titers from LR(V) to R(VI) observed in the probe-walk analysis suggested a high proportion of right-end reads terminating in the same region as left partials, it is plausible that long ssAAV partial genomes primed full-length ssAAV molecules and were converted to full-length copies during end-repair in long-read library preparation, potentially leading to under-quantification of partial genomes.

Based on the sharp titer drop from LR(V) to R(VI) in the probe-walk analysis (Fig. 5), we anticipated many right-end reads terminating in the same genomic region as the hotspot for left partials; however, long ssAAV partial genomes could prime full-length ssAAV molecules and be converted to full-length copies during library preparation end-repair step, resulting in under-quantification of partial genomes. Nonetheless, there was a strong concordance between the NanoMosaic probe walk analysis and long-read sequencing pertaining to the locations of the partials.

### Sedimentation Velocity Analytical Ultracentrifugation (SV-AUC)

SV-AUC was performed to estimate the sample’s density profile. Using AAV9 capsid density as the reference, SV-AUC estimated empty viral particles at 1.8%, partials at 4.6%, full-length genomes at 70.4%, and higher-molecular-weight (HMW) species at 18.5% (Fig. 7). Notably, partials constituted a smaller proportion than the designated full-length population, and HMW species with densities exceeding that of the full-length peak were also observed with a majority ranging from 105-150S (Fig. 7A).

**Figure 7:**
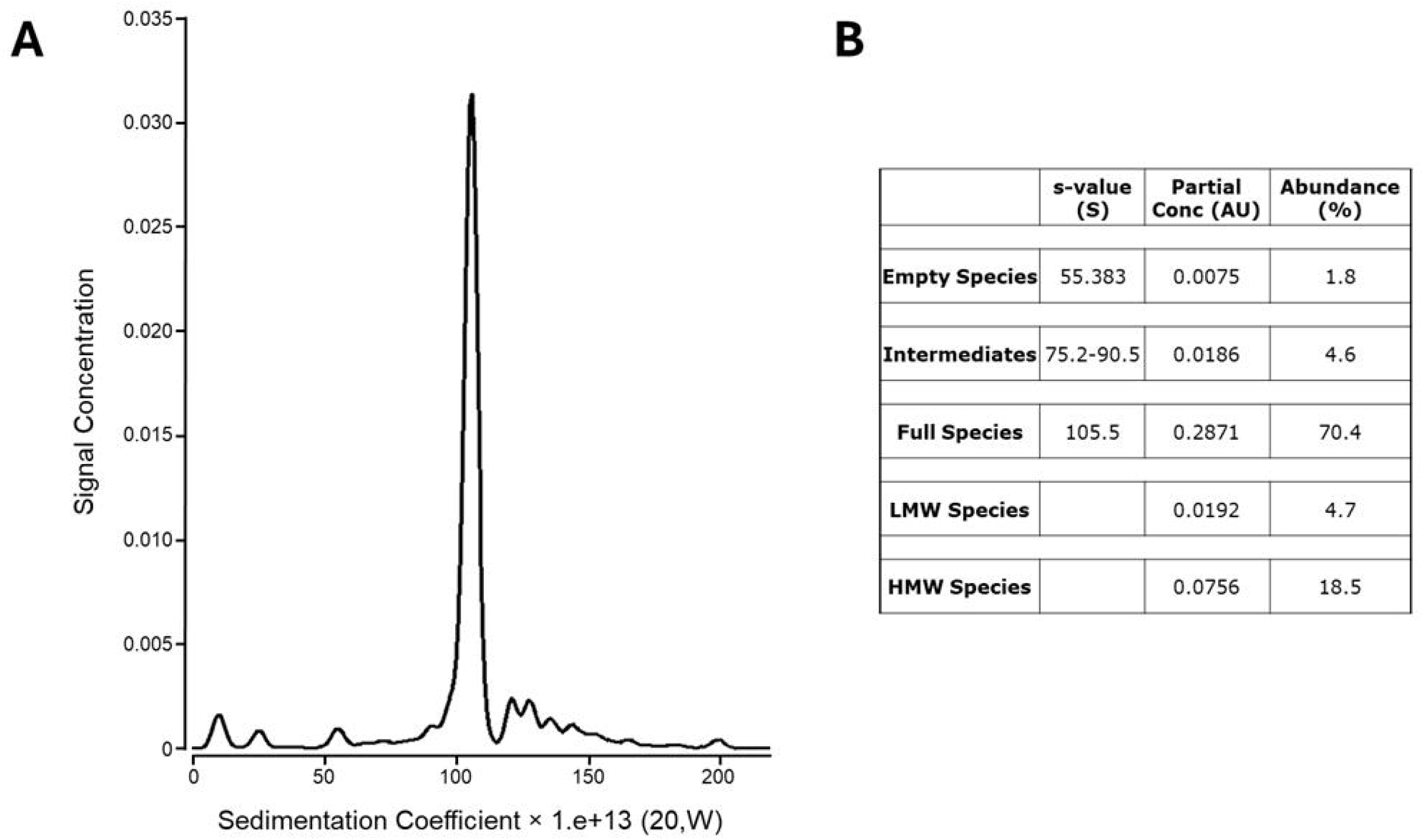
SV-AUC. **(A)** The distribution plot of the sedimentation coefficient by SV-AUC for the sample is shown. **(B)** The proportions of the analyzed fractions are listed (HMW: high molecular weight; LMW: low molecular weight species).

In the complementary probe-walk analysis, the full-length fraction represented only 3.2% of the population (Fig. 5B). When full-length genomes were grouped with right partials ≥2.22 kb [R(VII) region] or ≥3.28 kb [LR(V) region], the combined proportions were 36.4% and 22.5% (Fig. 5B), respectively, suggesting that larger partials may also contribute to the SV-AUC–designated full-length peak.

By long-read sequencing, full-length genomes were estimated at 55% of the population, whereas left and right partials together accounted for 26.5%. Notably, 15.9% of mapped reads were assigned to left-partial scAAV in the 500–2000 bp range with a peak around 1500 bp (reported on a double-stranded basis relative to the read-end position), which would be expected to produce a substantial partial peak between the empty (55S) and full peak (105S) by SV-AUC (Fig. 7); however, such a peak was not observed.

## Discussion

We describe a nanoneedle-based analytical platform (NanoMosaic “Tessie”) for AAV manufacturing analytics that quantifies capsid, full-length, and partial genomes. In contrast to workflows that rely on multiple instruments and consumables, this system uses a single instrument/consumable to run capsid and transgene titer assays in parallel, enabling one operator to execute both measurements and shortening process-development timelines. The assay design supports region-specific interrogation of the transgene, improving the accuracy of full-length and partial-genome quantification and revealing truncation hotspots that inform payload design and upstream optimization. The platform provides a standardized, matrix-agnostic quantification method that operates on crude extracts and purified material without filtration, uses SBS-format plates for high throughput and lower costs, requires only microliter sample volumes, and consolidates readouts into unified analytics and reporting pipeline— together strengthening QC relative to current fragmented approaches.

Applying the platform to a 4.5 kb transgene containing a GC-rich CAG promoter, we mapped a ∼570 bp truncation hotspot 0.44–1.01 kb from the left ITR. These “probe-walk” results highlight how targeted mapping can guide sequence engineering (e.g., mitigating secondary structures) and upstream process adjustments.

Whereas probe-walk interpretation is accurate when the sample population is uniformly ssAAV or scAAV genomes, it has limitations when both ssAAV and scAAV partial genomes are co-present in the sample, for e.g., when some genomes are partial self-complementary genomes (Xie J et al., 2017). The scAAV genomes are quantified as two ssAAV genomes thereby increasing ssAAV genome counts. Another consideration is that, during primer extension, partial genomes can also form heteroduplexes that prime synthesis of apparent full-length products. Using affinity primers at 10^4^–10^5^-fold excess over template reduces the probability of such heteroduplex-driven artifacts.

Comparison with PacBio long-read sequencing showed positional concordance for partial regions but not proportional agreement. Because PacBio library construction requires double-stranded DNA (Tran NT et al., 2024), complementary full-length ssAAV genomes can anneal to dsDNA and be preferentially ligated. Partial ssAAV species may hybridize and be repaired/end-polished toward full-length during library preparation—inflating full-length read counts—whereas unpaired ssDNA partials that fail adaptor ligation are lost (Hanscom T et al., 2023). Together, these effects undercount partial genomes. Consistent with this interpretation, 89% of left and right partials we detected were scAAV, indicating selection for double-stranded species. Additionally, ssAAV partials with hairpin-primed 3^′^ ends may be repaired during library prep to yield scAAV, analogous to short-hairpin–detoured viral replication (Xie J et al., 2017). Emerging AAV long-read methods that distinguish native from library-generated sequence (e.g., modified-base strategies) may mitigate these biases (Hanscom T et al., 2023).

SV-AUC identified a “full-length” peak that exceeded our molecular estimates by ∼23-fold and failed to mirror the partial-species proportions seen by probe-walk or PacBio long-read sequencing. Beyond this peak, 18.5% of signal resolved as higher–molecular-weight (HMW) species mainly spanning ∼105–150 S (Fig. 7A). AAV capsids can exist as dimeric or trimeric forms where more than one AAV particle can co-sediment in ultra-centrifugation experiments (Marunoa T et al., 2021; Hirohata K et al., 2024). Given an expected dimer:monomer S-ratio of ∼1.45 for AAV capsids, a full-length containing dimer capsid would be predicted near ∼152 S if a full-length containing monomeric capsid sediments at ∼105 S (Fig. 7B) (Marunoa T et al., 2021; Hirohata K et al., 2024). Since majority of the HMW fraction was between 105-152S, i.e., in between the size of a full-length containing genome and a dimer capsid containing full-length genome, it suggests that there is enrichment of multimeric species—potentially dimer capsids bearing partial genomes. Moreover, the apparent “full-length” peak may include multimers harboring shorter partial genomes whose density converges with that of true full-length capsids following CsCl purification, leading to co-sedimentation and inflation of the full-length estimate. Recent studies also indicate that environmental factors such as light and pH can degrade encapsidated DNA (Plegaria JS et al., 2025; Takino R et al., 2024), raising the possibility that packaged but degraded full-length transgenes co-sediment with intact packaged full-length transgene and further bias AUC-based quantification. Collectively, these effects provide a plausible mechanistic link between the SV-AUC and molecular readouts.

In sum, the NanoMosaic Tessie platform provides a rapid, high-throughput, and matrix-agnostic framework to assess key AAV quality attributes (capsid and genome titers, including partials), accelerating iterative upstream optimization and informing downstream purification strategies.

## Acknowledgements

We thank Kavita Sapru, Elena Prikhodko, Gokben Yildirir and Olga Vaganova from Catalent, Inc. (Baltimore, MD) for conducting SV-AUC and analyzing the data.

## Author Contributions

A.G., T.R., and Q.Q. designed research; A.G., E.L. and T.R. performed research; A.G., T.R, and Q.Q. analyzed data; and A.G. wrote the paper.

