## Supplementary File S1 for "Nanoneedle-Enabled Quantification of rAAV9 Capsid and Genome Integrity Reveals a Truncation Hotspot Locus in a 4.5 kb Transgene": COA1073227.pdf

VectorBuilder

### Certificate of Analysis

| | |
|---|---|
| **Product Name** | Ultra-purified recombinant AAV9 virus, large-scale packaging, P240709-1025jfz pAAV[Exp]-CAG>Luciferase:WPRE (VB211007-1286euq) |
| **Catalog #** | P240709-1025jfz |
| **Lot #** | 240726AAVN01 |
| **Production Date** | 2024-07-26 |
| **Storage Buffer** | PBS buffer (pH7.4) supplemented with 200 mM NaCl and 0.001% pluronic F-68 |
| **Storage Condition** | Store at –80°C |
| **Shelf Life** | One year under proper storage conditions |
| **Order #** | S240717-1019rzr-r1 |

The product is functionally tested for optimum performance using the following tests:

**Titer Determination**

Titer was determined by "quantitative PCR (qPCR)" method. Viral DNA was extracted from viral particles. The ITR sequence of AAV2 was used as a target to quantify the amount of viral genome.

Specification: >$10^{13}$ GC/ml — Result: $2.51 \times 10^{13}$ GC/ml

**Titer Determination**

Titer was determined by "droplet digital PCR (ddPCR)" method. Viral DNA was extracted from viral particles. The ITR sequence of AAV2 was used as a target to quantify the amount of viral genome.

Specification: None — Result: $1.16 \times 10^{13}$ GC/ml

**Sterility Test**

Virus sample was inoculated into culture medium to detect bacterial and fungal growth.

Specification: Negative — Result: Pass

**Mycoplasma Test**

Mycoplasma contamination was detected by inoculating viral sample into indicator cells. Cells were then stained for DNA with a fluorescent dye, and mycoplasmas that had been in your cell sample would be visible as fluorescent spots, or granules, surrounding the nucleus of the indicator cells.

Specification: Negative — Result: Pass

**Virus Purity**

Viral purity was determined by SDS–PAGE followed by silver staining.

Specification: >80% Pure — Result: Pass

---

All products are for research use only. Caution: Not intended for human or animal diagnostic or therapeutic uses. For inquiries, contact us at.

Bliss Li
