## Supplementary File S1 for "Nanoneedle-Enabled Quantification of rAAV9 Capsid and Genome Integrity Reveals a Truncation Hotspot Locus in a 4.5 kb Transgene": VB211007-1286euq(pAAV[Exp]-CAG-Luciferase&WPRE).pdf

### Vector Summary

| | |
|---|---|
| Vector ID | VB211007-1286euq |
| Vector Name | pAAV[Exp]-CAG>Luciferase:WPRE |
| Vector Size | 7195 bp |
| Viral Genome Size | 4576 bp |
| Vector Type | Mammalian Gene Expression AAV Vector |
| Inserted Promoter | CAG |
| Inserted ORF | Luciferase |
| Inserted Regulatory Element | WPRE |
| Plasmid Copy Number | High |
| Antibiotic Resistance | Ampicillin |
| Cloning Host | VB UltraStable (or alternative strain) |

### Vector Map

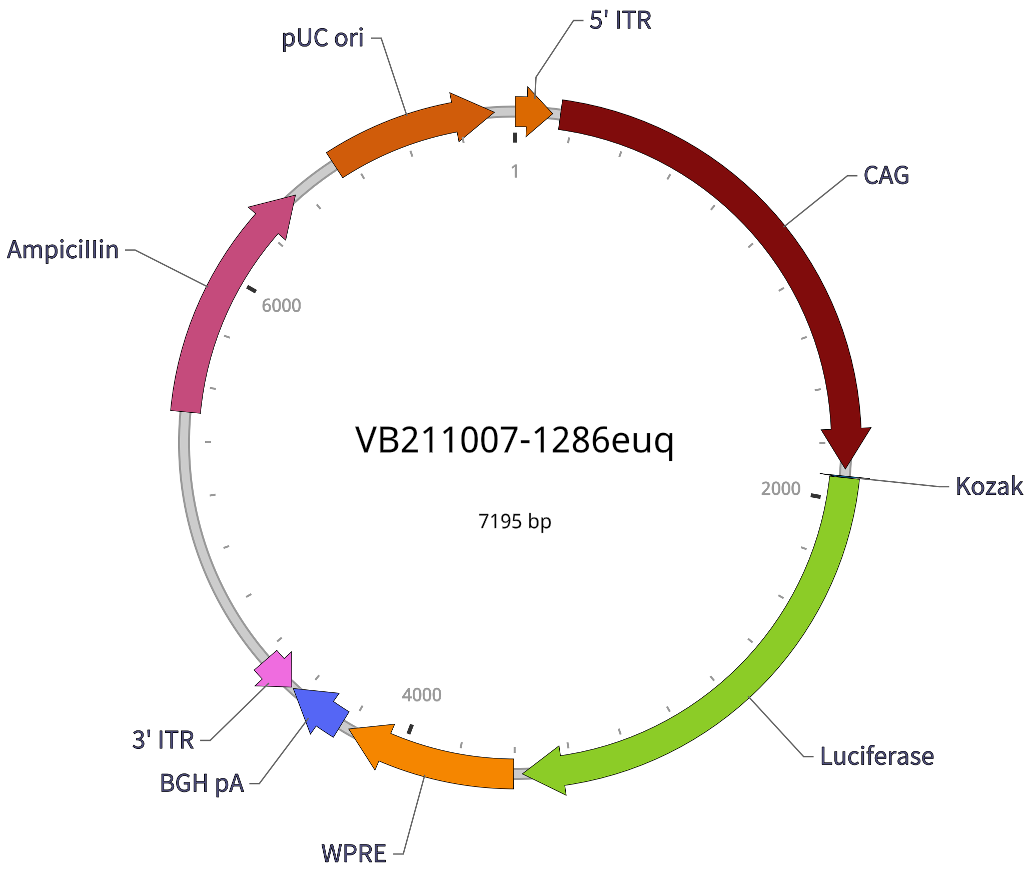


### Vector Components

| Name | Position | Size (bp) | Type | Description | Application notes |
|---|---|---|---|---|---|
| 5' ITR | 1-130 | 130 | ITR | AAV 5' inverted terminal repeat (functional equivalent of wild-type 5' ITR) | Allows rescue of virus from recombinant plasmid and replication of the viral genome; this ITR is identical to that of the wild-type AAV2 genome. |
| CAG | 158-1890 | 1733 | Promoter | CMV early enhancer fused to modified chicken β-actin promoter | Strong promoter. |
| Kozak | 1915-1920 | 6 | Miscellaneous | Kozak translation initiation sequence | Facilitates translation initiation of ATG start codon downstream of the Kozak sequence. |
| Luciferase | 1921-3573 | 1653 | CDS | Firefly luciferase | Most commonly used luciferase. |
| WPRE | 3604-4201 | 598 | Miscellaneous | Woodchuck hepatitis virus posttranscriptional regulatory element | Enhances virus stability in packaging cells, leading to higher titer of packaged virus; enhances higher expression of transgenes. |
| BGH pA | 4232-4439 | 208 | PolyA_signal | Bovine growth hormone polyadenylation signal | Allows transcription termination and polyadenylation of mRNA transcribed by Pol II RNA polymerase. |
| 3' ITR | complement (4447-4576) | 130 | ITR | AAV 3' inverted terminal repeat | Allows rescue of virus from recombinant plasmid and replication of the viral genome; this ITR is identical to that of the wild-type AAV2 genome. |
| AmpicilIin | 5504-6364 | 861 | CDS | AmpicilIin resistance gene | Allows E. coli to be resistant to ampicilIin. |
| pUC ori | 6535-7123 | 589 | Rep_origin | pUC origin of replication | Facilitates plasmid replication in E. coli; regulates high-copy plasmid number (500-700). |

**Note:** Components added by user are listed in **bold red** text.

### Vector Sequence

1 CTGCGCGCTC GCTCGCTCAC TGAGGCCGCC CGGGCAAAGC CCGGGCGTCG GGCGACCTTT GGTCGCCCGG CCTCAGTGAG
81 CGAGCGAGCG CGCAGAGAGG GAGTGGCCAA CTCCATCACT AGGGGTTCCT TCTAGACAAC TTTGTATAGA AAAGTTGCTC
161 GACATTGATT ATTGACTAGT TATTAATAGT AATCAATTAC GGGGTCATTA GTTCATAGCC CATATATGGA GTTCCGCGTT
241 ACATAACTTA CGGTAAATGG CCCGCCTGGC TGACCGCCCA ACGACCCCCG CCCATTGACG TCAATAATGA CGTATGTTCC
321 CATAGTAACG CCAATAGGGA CTTTCCATTG ACGTCAATGG GTGGAGTATT TACGGTAAAC TGCCCACTTG GCAGTACATC
401 AAGTGTATCA TATGCCAAGT ACGCCCCCTA TTGACGTCAA TGACGGTAAA TGGCCCGCCT GGCATTATGC CCAGTACATG
481 ACCTTATGGG ACTTTCCTAC TTGGCAGTAC ATCTACGTAT TAGTCATCGC TATTACCATG GTCGAGGTGA GCCCCACGTT
561 CTGCTTCACT CTCCCCATCT CCCCCCCCTC CCCACCCCCA ATTTTGTATT TATTTATTTT TTAATTATTT TGTGCAGCGA
641 TGGGGGCGGG GGGGGGGGGG GGGCGCGCGC CAGGCGGGGC GGGGCGGGGC GAGGGGCGGG GCGGGGCGAG GCGGAGAGGT
721 GCGGCGGCAG CCAATCAGAG CGGCGCGCTC CGAAAGTTTC CTTTTATGGC GAGGCGGCGG CGGCGGCGGC CCTATAAAAA
801 GCGAAGCGCG CGGCGGGCGG GAGTCGCTGC GCGCTGCCTT CGCCCCGTGC CCCGCTCCGC CGCCGCCTCG CGCCGCCCGC
881 CCCGGCTCTG ACTGACCGCG TTACTCCCAC AGGTGAGCGG GCGGGACGGC CCTTCTCCTC CGGGCTGTAA TTAGCGCTTG
961 GTTTAATGAC GGCTTGTTTC TTTTCTGTGG CTGCGTGAAA GCCTTGAGGG GCTCCGGGAG GGCCCTTTGT GCGGGGGGAG
1041 CGGCTCGGGG GGTGCGTGCG TGTGTGTGTG CGTGGGGAGC GCCGCGTGCG GCTCCGCGCT GCCCGGCGGC TGTGAGCGCT
1121 GCGGGCGCGG CGCGGGGCTT TGTGCGCTCC GCAGTGTGCG CGAGGGGAGC GCGGCCGGGG GCGGTGCCCC GCGGTGCGGG
1201 GGGGGCTGCG AGGGGAACAA AGGCTGCGTG CGGGGTGTGT GCGTGGGGGG GTGAGCAGGG GGTGTGGGCG CGTCGGTCGG
1281 GCTGCAACCC CCCCTGCACC CCCCTCCCCG AGTTGCTGAG CACGGCCCGG CTTCGGGTGC GGGGCTCCGT ACGGGGCGTG
1361 GCGCGGGGCT CGCCGTGCCG GGCGGGGGGT GGCGGCAGGT GGGGGTGCCG GGCGGGGCGG GGCCGCCTCG GGCCGGGGAG
1441 GGCTCGGGGG AGGGGCGCGG CGGCCCCCGG AGCGCCGGCG GCTGTCGAGG CGCGGCGAGC CGCAGCCATT GCCTTTTATG
1521 GTAATCGTGC GAGAGGGCGC AGGGACTTCC TTTGTCCCAA ATCTGTGCGG AGCCGAAATC TGGGAGGCGC CGCCGCACCC
1601 CCTCTAGCGG GCGCGGGGCG AAGCGGTGCG GCGCCGGCAG GAAGGAAATG GGCGGGGAGG GCCTTCGTGC GTCGCCGCGC
1681 CGCCGTCCCC TTCTCCCTCT CCAGCCTCGG GGCTGTCCGC GGGGGGACGG CTGCCTTCGG GGGGGACGGG GCAGGGCGGG
1761 GTTCGGCTTC TGGCGTGTGA CCGGCGGCTC TAGAGCCTCT GCTAACCATG TTCATGCCTT CTTCTTTTTC CTACAGCTCC
1841 TGGGCAACGT GCTGGTTATT GTGCTGTCTC ATCATTTTGG CAAAGAATTG CAAGTTTGTA CAAAAAAGCA GGCTGCCACC
1921 ATGGAAGACG CCAAAAACAT AAAGAAAGGC CCGGCGCCAT TCTATCCGCT AGAGGATGGA ACCGCTGGAG AGCAACTGCA
2001 TAAGGCTATG AAGAGATACG CCCTGGTTCC TGGAACAATT GCTTTTACAG ATGCACATAT CGAGGTGAAC ATCACGTACG
2081 CGGAATACTT CGAAATGTCC GTTCGGTTGG CAGAAGCTAT GAAACGATAT GGGCTGAATA CAAATCACAG AATCGTCGTA
2161 TGCAGTGAAA ACTCTCTTCA ATTCTTTATG CCGGTGTTGG GCGCGTTATT TATCGGAGTT GCAGTTGCGC CCGCGAACGA
2241 CATTTATAAT GAACGTGAAT TGCTCAACAG TATGAACATT TCGCAGCCTA CCGTAGTGTT TGTTTCCAAA AAGGGGTTGC
2321 AAAAAATTTT GAACGTGCAA AAAAAATTAC CAATAATCCA GAAAATTATT ATCATGGATT CTAAAACGGA TTACCAGGGA
2401 TTTCAGTCGA TGTACACGTT CGTCACATCT CATCTACCTC CCGGTTTTAA TGAATACGAT TTTGTACCAG AGTCCTTTGA
2481 TCGTGACAAA ACAATTGCAC TGATAATGAA CTCCTCTGGA TCTACTGGGT TACCTAAGGG TGTGGCCCTT CCGCATAGAA
2561 CTGCCTGCGT CAGATTCTCG CATGCCAGAG ATCCTATTTT TGGCAATCAA ATCATTCCGG ATACTGCGAT TTTAAGTGTT
2641 GTTCCATTCC ATCACGGTTT TGGAATGTTT ACTACACTCG GATATTTGAT ATGTGGATTT CGAGTCGTCT TAATGTATAG
2721 ATTTGAAGAA GAGCTGTTTT TACGATCCCT TCAGGATTAC AAAATTCAAA GTGCGTTGCT AGTACCAACC CTATTTTCAT
2801 TCTTCGCCAA AAGCACTCTG ATTGACAAAT ACGATTTATC TAATTTACAC GAAATTGCTT CTGGGGGCGC ACCTCTTTCG
2881 AAAGAAGTCG GGGAAGCGGT TGCAAAACGC TTCCATCTTC CAGGGATACG ACAAGGATAT GGGCTCACTG AGACTACATC
2961 AGCTATTCTG ATTACACCCG AGGGGGATGA TAAACCGGGC GCGGTCGGTA AAGTTGTTCC ATTTTTTGAA GCGAAGGTTG
3041 TGGATCTGGA TACCGGGAAA ACGCTGGGCG TTAATCAGAG AGGCGAATTA TGTGTCAGAG GACCTATGAT TATGTCCGGT
3121 TATGTAAACA ATCCGGAAGC GACCAACGCC TTGATTGACA AGGATGGATG GCTACATTCT GGAGACATAG CTTACTGGGA
3201 CGAAGACGAA CACTTCTTCA TAGTTGACCG CTTGAAGTCT TTAATTAAAT ACAAAGGATA CCAGGTGGCC CCCGCTGAAT
3281 TGGAGTCGAT ATTGTTACAA CACCCCAACA TCTTCGACGC GGGCGTGGCA GGTCTTCCCG ACGATGACGC CGGTGAACTT
3361 CCCGCCGCCG TTGTTGTTTT GGAGCACGGA AAGACGATGA CGGAAAAAGA GATCGTGGAT TACGTCGCCA GTCAAGTAAC

3441 AACCGCGAAA AAGTTGCGCG GAGGAGTTGT GTTTGTGGAC GAAGTACCGA AAGGTCTTAC CGGAAAACTC GACGCAAGAA
3521 AAATCAGAGA GATCCTCATA AAGGCCAAGA AGGGCGGAAA GTCCAAATTG TAAACCCAGC TTTCTTGTAC AAAGTGGGAA
3601 TTCCGATAAT CAACCTCTGG ATTACAAAAT TTGTGAAAGA TTGACTGGTA TTCTTAACTA TGTTGCTCCT TTTACGCTAT
3681 GTGGATACGC TGCTTTAATG CCTTTGTATC ATGCTATTGC TTCCCGTATG GCTTTCATTT TCTCCTCCTT GTATAAATCC
3761 TGGTTGCTGT CTCTTTATGA GGAGTTGTGG CCCGTTGTCA GGCAACGTGG CGTGGTGTGC ACTGTGTTTG CTGACGCAAC
3841 CCCCACTGGT TGGGGCATTG CCACCACCTG TCAGCTCCTT TCCGGGACTT TCGCTTTCCC CCTCCCTATT GCCACGGCGG
3921 AACTCATCGC CGCCTGCCTT GCCCGCTGCT GGACAGGGGC TCGGCTGTTG GGCACTGACA ATTCCGTGGT GTTGTCGGGG
4001 AAGCTGACGT CCTTTCCATG GCTGCTCGCC TGTGTTGCCA CCTGGATTCT GCGCGGGACG TCCTTCTGCT ACGTCCCTTC
4081 GGCCCTCAAT CCAGCGGACC TTCCTTCCCG CGGCCTGCTG CCGGCTCTGC GGCCTCTTCC GCGTCTTCGC CTTCGCCCTC
4161 AGACGAGTCG GATCTCCCTT TGGGCCGCCT CCCCGCATCG GGAATTCCTA GAGCTCGCTG ATCAGCCTCG ACTGTGCCTT
4241 CTAGTTGCCA GCCATCTGTT GTTTGCCCCT CCCCCGTGCC TTCCTTGACC CTGGAAGGTG CCACTCCCAC TGTCCTTTCC
4321 TAATAAAATG AGGAAATTGC ATCGCATTGT CTGAGTAGGT GTCATTCTAT TCTGGGGGGT GGGGTGGGGC AGGACAGCAA
4401 GGGGGAGGAT TGGGAAGAGA ATAGCAGGCA TGCTGGGGAG GGCCGCAGGA ACCCCTAGTG ATGGAGTTGG CCACTCCCTC
4481 TCTGCGCGCT CGCTCGCTCA CTGAGGCCGG GCGACCAAAG GTCGCCCGAC GCCCGGGCTT TGCCCGGGCG GCCTCAGTGA
4561 GCGAGCGAGC GCGCAGCTGC CTGCAGGGGC GCCTGATGCG GTATTTTCTC CTTACGCATC TGTGCGGTAT TTCACACCGC
4641 ATACGTCAAA GCAACCATAG TACGCGCCCT GTAGCGGCGC ATTAAGCGCG GCGGGTGTGG TGGTTACGCG CAGCGTGACC
4721 GCTACACTTG CCAGCGCCTT AGCGCCCGCT CCTTTCGCTT TCTTCCCTTC CTTTCTCGCC ACGTTCGCCG GCTTTCCCCG
4801 TCAAGCTCTA AATCGGGGGC TCCCTTTAGG GTTCCGATTT AGTGCTTTAC GGCACCTCGA CCCCAAAAAA CTTGATTTGG
4881 GTGATGGTTC ACGTAGTGGG CCATCGCCCT GATAGACGGT TTTTCGCCCT TTGACGTTGG AGTCCACGTT CTTTAATAGT
4961 GGACTCTTGT TCCAAACTGG AACAACACTC AACTCTATCT CGGGCTATTC TTTTGATTTA TAAGGGATTT TGCCGATTTC
5041 GGTCTATTGG TTAAAAAATG AGCTGATTTA ACAAAAATTT AACGCGAATT TTAACAAAAT ATTAACGTTT ACAATTTTAT
5121 GGTGCACTCT CAGTACAATC TGCTCTGATG CCGCATAGTT AAGCCAGCCC CGACACCCGC CAACACCCGC TGACGCGCCC
5201 TGACGGGCTT GTCTGCTCCC GGCATCCGCT TACAGACAAG CTGTGACCGT CTCCGGGAGC TGCATGTGTC AGAGGTTTTC
5281 ACCGTCATCA CCGAAACGCG CGAGACGAAA GGGCCTCGTG ATACGCCTAT TTTTATAGGT TAATGTCATG ATAATAATGG
5361 TTTCTTAGAC GTCAGGTGGC ACTTTTCGGG GAAATGTGCG CGGAACCCCT ATTTGTTTAT TTTTCTAAAT ACATTCAAAT
5441 ATGTATCCGC TCATGAGACA ATAACCCTGA TAAATGCTTC AATAATATTG AAAAAGGAAG AGTATGAGTA TTCAACATTT
5521 CCGTGTCGCC CTTATTCCCT TTTTTGCGGC ATTTTGCCTT CCTGTTTTTG CTCACCCAGA AACGCTGGTG AAAGTAAAAG
5601 ATGCTGAAGA TCAGTTGGGT GCACGAGTGG GTTACATCGA ACTGGATCTC AACAGCGGTA AGATCCTTGA GAGTTTTCGC
5681 CCCGAAGAAC GTTTTCCAAT GATGAGCACT TTTAAAGTTC TGCTATGTGG CGCGGTATTA TCCCGTATTG ACGCCGGGCA
5761 AGAGCAACTC GGTCGCCGCA TACACTATTC TCAGAATGAC TTGGTTGAGT ACTCACCAGT CACAGAAAAG CATCTTACGG
5841 ATGGCATGAC AGTAAGAGAA TTATGCAGTG CTGCCATAAC CATGAGTGAT AACACTGCGG CCAACTTACT TCTGACAACG
5921 ATCGGAGGAC CGAAGGAGCT AACCGCTTTT TTGCACAACA TGGGGGATCA TGTAACTCGC CTTGATCGTT GGGAACCGGA
6001 GCTGAATGAA GCCATACCAA ACGACGAGCG TGACACCACG ATGCCTGTAG CAATGGCAAC AACGTTGCGC AAACTATTAA
6081 CTGGCGAACT ACTTACTCTA GCTTCCCGGC AACAATTAAT AGACTGGATG GAGGCGGATA AAGTTGCAGG ACCACTTCTG
6161 CGCTCGGCCC TTCCGGCTGG CTGGTTTATT GCTGATAAAT CTGGAGCCGG TGAGCGTGGA AGCCGCGGTA TCATTGCAGC
6241 ACTGGGGCCA GATGGTAAGC CCTCCCGTAT CGTAGTTATC TACACGACGG GGAGTCAGGC AACTATGGAT GAACGAAATA
6321 GACAGATCGC TGAGATAGGT GCCTCACTGA TTAAGCATTG GTAACTGTCA GACCAAGTTT ACTCATATAT ACTTTAGATT
6401 GATTTAAAAC TTCATTTTTA ATTTAAAAGG ATCTAGGTGA AGATCCTTTT TGATAATCTC ATGACCAAAA TCCCTTAACG
6481 TGAGTTTTCG TTCCACTGAG CGTCAGACCC CGTAGAAAAG ATCAAAGGAT CTTCTTGAGA TCCTTTTTTT CTGCGCGTAA
6561 TCTGCTGCTT GCAAACAAAA AAACCACCGC TACCAGCGGT GGTTTGTTTG CCGGATCAAG AGCTACCAAC TCTTTTTCCG
6641 AAGGTAACTG GCTTCAGCAG AGCGCAGATA CCAAATACTG TTCTTCTAGT GTAGCCGTAG TTAGGCCACC ACTTCAAGAA
6721 CTCTGTAGCA CCGCCTACAT ACCTCGCTCT GCTAATCCTG TTACCAGTGG CTGCTGCCAG TGGCGATAAG TCGTGTCTTA
6801 CCGGGTTGGA CTCAAGACGA TAGTTACCGG ATAAGGCGCA GCGGTCGGGC TGAACGGGGG GTTCGTGCAC ACAGCCCAGC
6881 TTGGAGCGAA CGACCTACAC CGAACTGAGA TACCTACAGC GTGAGCTATG AGAAAGCGCC ACGCTTCCCG AAGGGAGAAA
6961 GGCGGACAGG TATCCGGTAA GCGGCAGGGT CGGAACAGGA GAGCGCACGA GGGAGCTTCC AGGGGGAAAC GCCTGGTATC
7041 TTTATAGTCC TGTCGGGTTT CGCCACCTCT GACTTGAGCG TCGATTTTTG TGATGCTCGT CAGGGGGGCG GAGCCTATGG
7121 AAAAACGCCA GCAACGCGGC CTTTTTACGG TTCCTGGCCT TTTGCTGGCC TTTTGCTCAC ATGTCCTGCA GGCAG

### Validation by Restriction Enzyme Digestion

| Restriction Enzymes | Cutting Sites | DNA Fragments (bp) |
|---|---|---|
| ApaLI | 3818, 5123, 5620, 6866 | 1305, 497, 1246, 4147 |
| EcoRI | 3599, 4203 | 604, 6591 |
| DraIII | 4896 | 7195 |
| ApaLI+EcoRI | 3599, 3818, 4203, 5123, 5620, 6866 | 219, 385, 920, 497, 1246, 3928 |
| ApaLI+DraIII | 3818, 4896, 5123, 5620, 6866 | 1078, 227, 497, 1246, 4147 |
