## Supplementary File S3 for "Nanoneedle-Enabled Quantification of rAAV9 Capsid and Genome Integrity Reveals a Truncation Hotspot Locus in a 4.5 kb Transgene": 20250408_VB_probe_walk_experiment_T23_2-46_2-32_and_2-25_2-32.pdf

### Tessie Analysis Report
#### 96 well, 1 plex

| | |
|---|---|
| Plate Type: | AA01 |
| Assay Design: | 20250408_VB_probe_walk_experiment_T23_2-46_2-32_and_2-25_2-32 |
| Scan Package: | T1023_00006921_1_1 |
| Instrument ID: | T1023 |
| Plate ID: | 00006921 |
| Pre Scan User: | N/A |
| Pre Scan Number: | 1 |
| Pre Scan Time: | 2025-04-08 13:08:39-04:00 |
| Post Scan User: | N/A |
| Post Scan Number: | 1 |
| Post Scan Time: | 2025-04-09 14:30:58-04:00 |
| Image Analysis Version: | 1.1.0.0 |
| Report Generation Time: | 2025-04-09 15:33:48-04:00 |
| Markers: | 2-46, 2-32, 2-25, 2-32 |

### NanoUnits Heatmap

| | 3 | 4 | 5 | 6 | 7 | 8 | 9 | 10 |
|---|---|---|---|---|---|---|---|---|
| A | 2-46, 2-32 0 | 2-46, 2-32 0 | 2-46, 2-32 VB 4.5kb 2048000x | 2-46, 2-32 VB 4.5kb 2048000x | 2-25, 2-32 0 | 2-25, 2-32 0 | 2-25, 2-32 VB 4.5kb 64000x | 2-25, 2-32 VB 4.5kb 64000x |
| B | 2-46, 2-32 8.83E+06 | 2-46, 2-32 8.83E+06 | 2-46, 2-32 VB 4.5kb 1024000x | 2-46, 2-32 VB 4.5kb 1024000x | 2-25, 2-32 7.06E+07 | 2-25, 2-32 7.06E+07 | 2-25, 2-32 VB 4.5kb 32000x | 2-25, 2-32 VB 4.5kb 32000x |
| C | 2-46, 2-32 1.77E+07 | 2-46, 2-32 1.77E+07 | 2-46, 2-32 VB 4.5kb 512000x | 2-46, 2-32 VB 4.5kb 512000x | 2-25, 2-32 1.41E+08 | 2-25, 2-32 1.41E+08 | 2-25, 2-32 VB 4.5kb 16000x | 2-25, 2-32 VB 4.5kb 16000x |
| D | 2-46, 2-32 3.53E+07 | 2-46, 2-32 3.53E+07 | 2-46, 2-32 VB 4.5kb 256000x | 2-46, 2-32 VB 4.5kb 256000x | 2-25, 2-32 2.82E+08 | 2-25, 2-32 2.82E+08 | 2-25, 2-32 VB 4.5kb 8000x | 2-25, 2-32 VB 4.5kb 8000x |
| E | 2-46, 2-32 7.06E+07 | 2-46, 2-32 7.06E+07 | 2-46, 2-32 VB 4.5kb 128000x | 2-46, 2-32 VB 4.5kb 128000x | 2-25, 2-32 5.65E+08 | 2-25, 2-32 5.65E+08 | 2-25, 2-32 VB 4.5kb 4000x | 2-25, 2-32 VB 4.5kb 4000x |
| F | 2-46, 2-32 1.41E+08 | 2-46, 2-32 1.41E+08 | 2-46, 2-32 VB 4.5kb 64000x | 2-46, 2-32 VB 4.5kb 64000x | 2-25, 2-32 1.13E+09 | 2-25, 2-32 1.13E+09 | 2-25, 2-32 VB 4.5kb 2000x | 2-25, 2-32 VB 4.5kb 2000x |
| G | 2-46, 2-32 2.82E+08 | 2-46, 2-32 2.82E+08 | 2-46, 2-32 VB 4.5kb 32000x | 2-46, 2-32 VB 4.5kb 32000x | 2-25, 2-32 2.26E+09 | 2-25, 2-32 2.26E+09 | 2-25, 2-32 VB 4.5kb 1000x | 2-25, 2-32 VB 4.5kb 1000x |
| H | 2-46, 2-32 5.65E+08 | 2-46, 2-32 5.65E+08 | 2-46, 2-32 VB 4.5kb 16000x | 2-46, 2-32 VB 4.5kb 16000x | 2-25, 2-32 4.52E+09 | 2-25, 2-32 4.52E+09 | 2-25, 2-32 VB 4.5kb 500x | 2-25, 2-32 VB 4.5kb 500x |

Color scale: 0, 1, 2, 3, 4, 5, 6, 7, 8, 9, 10

#### 2-46, 2-32 Standard Curve

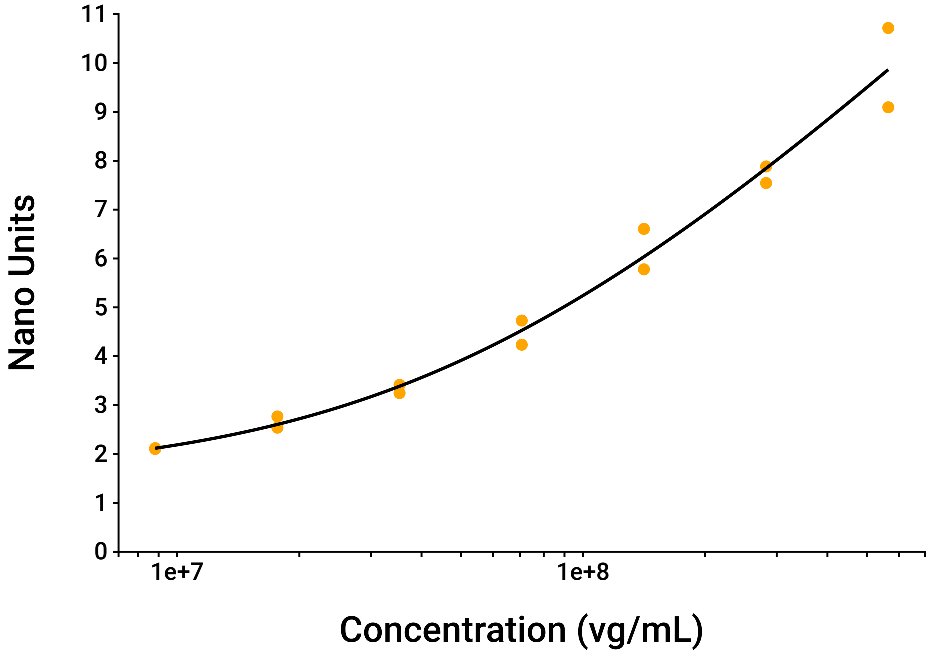


#### 2-25, 2-32 Standard Curve

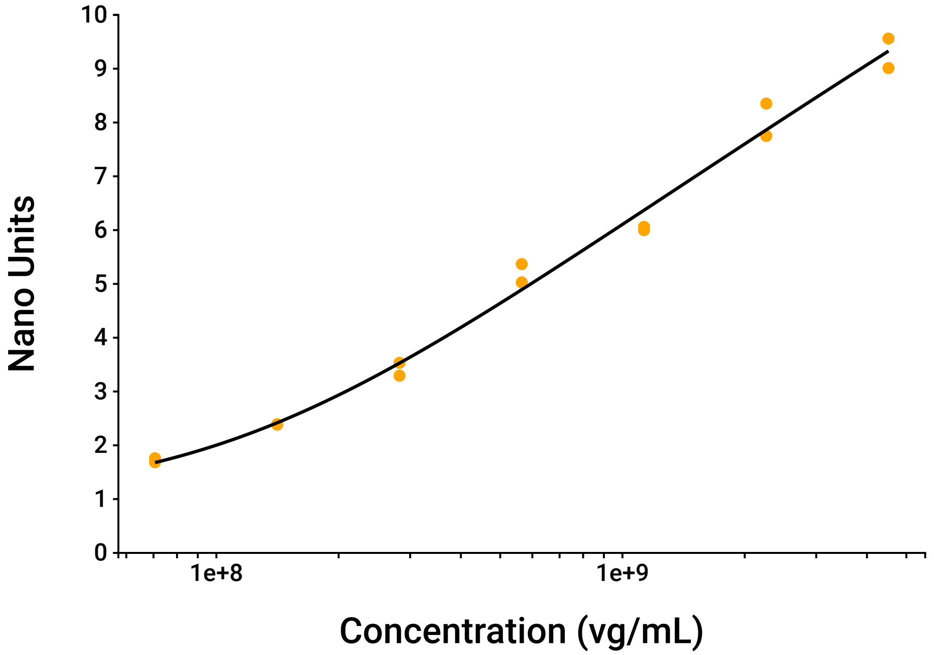

### VB 4.5kb Dilutions Data

| Target | Dilution Factor | Measured Concentration | Undiluted Concentration | %CV | Linearity %CV |
|---|---|---|---|---|---|
| 2-46, 2-32 (vg/mL) | 128000 | 2.65E+08 | 3.39E+13 | 0.38 | 9.76 |
| | 256000 | 1.23E+08 | 3.15E+13 | 12.9 | |
| | 512000 | 5.44E+07 | 2.78E+13 | 4.90 | |
| 2-25, 2-32 (vg/mL) | 1000 | 1.39E+09 | 1.39E+12 | 2.43 | 14.9 |
| | 2000 | 5.19E+08 | 1.04E+12 | 7.47 | |
| | 4000 | 2.93E+08 | 1.17E+12 | 5.74 | |

### 2-46, 2-32 Standard Data

| Concentration (vg/mL) | Nano Units | | |
|---|---|---|---|
| | Mean | Standard Deviation | %CV |
| 0 | 1.5 | 0.065 | 4.34 |
| 8.83E+06 | 2.11 | 0.0119 | 0.56 |
| 1.77E+07 | 2.65 | 0.162 | 6.12 |
| 3.53E+07 | 3.33 | 0.119 | 3.57 |
| 7.06E+07 | 4.48 | 0.35 | 7.81 |
| 1.41E+08 | 6.19 | 0.585 | 9.45 |
| 2.82E+08 | 7.71 | 0.24 | 3.11 |
| 5.65E+08 | 9.9 | 1.15 | 11.6 |

### 2-46, 2-32 Sample Data

| Sample Label | Concentration (vg/mL) | | |
|---|---|---|---|
| | Mean | Standard Deviation | %CV |
| VB 4.5kb | 3.11E+13 | 3.32E+12 | 10.7 |

### 2-25, 2-32 Standard Data

| Concentration (vg/mL) | Nano Units | | |
|---|---|---|---|
| | Mean | Standard Deviation | %CV |
| 0 | 0.953 | 0.0159 | 1.67 |
| 7.06E+07 | 1.72 | 0.0524 | 3.04 |
| 1.41E+08 | 2.39 | 0.00561 | 0.24 |
| 2.82E+08 | 3.41 | 0.169 | 4.94 |
| 5.65E+08 | 5.2 | 0.24 | 4.62 |
| 1.13E+09 | 6.03 | 0.0438 | 0.73 |
| 2.26E+09 | 8.05 | 0.426 | 5.29 |
| 4.52E+09 | 9.29 | 0.388 | 4.18 |

### 2-25, 2-32 Sample Data

| Sample Label | Concentration (vg/mL) | | |
|---|---|---|---|
| | Mean | Standard Deviation | %CV |
| VB 4.5kb | 1.2E+12 | 1.67E+11 | 13.9 |

### Curve Parameters

| Target | Curve | Equation | Parameters | $R^2$ |
|---|---|---|---|---|
| 2-46, 2-32 | Five parameter logisitic curve | $y = D - \frac{(A-D)}{\left[1+(x/C)^B\right]^E}$ | A = 1.5<br>B = 0.941<br>C = 5.16E+07<br>D = 2.21E+04<br>E = 0.000161 | 0.984 |
| 2-25, 2-32 | Five parameter logisitic curve | $y = D - \frac{(A-D)}{\left[1+(x/C)^B\right]^E}$ | A = 0.962<br>B = 1.35<br>C = 1.26E+08<br>D = 46.1<br>E = 0.0423 | 0.993 |

### Raw Data

| Sensor | Status | Design | | | | Result | | |
|---|---|---|---|---|---|---|---|---|
| | | Target | Concentration | Sample ID | Dilution Factor | Nano Units | Measured Conc. | Undiluted Conc. |
| A3 | Success | 2-46, 2-32 | 0 | -- | -- | 1.45 | -- | -- |
| A4 | Success | 2-46, 2-32 | 0 | -- | -- | 1.54 | -- | -- |
| A5 | Excluded | 2-46, 2-32 | -- | VB 4.5kb | 2.05E+06 | 2.2 | 1.03E+07 | 2.11E+13 |
| A6 | Excluded | 2-46, 2-32 | -- | VB 4.5kb | 2.05E+06 | 2.57 | 1.7E+07 | 3.48E+13 |
| A7 | Success | 2-25, 2-32 | 0 | -- | -- | 0.964 | -- | -- |
| A8 | Success | 2-25, 2-32 | 0 | -- | -- | 0.942 | -- | -- |
| A9 | Excluded | 2-25, 2-32 | -- | VB 4.5kb | 6.4E+04 | 1.09 | LOW | -- |
| A10 | Excluded | 2-25, 2-32 | -- | VB 4.5kb | 6.4E+04 | 1.25 | LOW | -- |
| B3 | Success | 2-46, 2-32 | 8.83E+06 | -- | -- | 2.1 | -- | -- |
| B4 | Success | 2-46, 2-32 | 8.83E+06 | -- | -- | 2.12 | -- | -- |
| B5 | Excluded | 2-46, 2-32 | -- | VB 4.5kb | 1.02E+06 | 3.07 | 2.75E+07 | 2.82E+13 |
| B6 | Excluded | 2-46, 2-32 | -- | VB 4.5kb | 1.02E+06 | 3.14 | 2.92E+07 | 2.99E+13 |
| B7 | Success | 2-25, 2-32 | 7.06E+07 | -- | -- | 1.69 | -- | -- |
| B8 | Success | 2-25, 2-32 | 7.06E+07 | -- | -- | 1.76 | -- | -- |
| B9 | Excluded | 2-25, 2-32 | -- | VB 4.5kb | 3.2E+04 | 1.33 | LOW | -- |
| B10 | Excluded | 2-25, 2-32 | -- | VB 4.5kb | 3.2E+04 | 1.27 | LOW | -- |
| C3 | Success | 2-46, 2-32 | 1.77E+07 | -- | -- | 2.77 | -- | -- |

| Sensor | Status | Design | | | | Result | | |
|---|---|---|---|---|---|---|---|---|
| | | Target | Concentration | Sample ID | Dilution Factor | Nano Units | Measured Conc. | Undiluted Conc. |
| C4 | Success | 2-46, 2-32 | 1.77E+07 | -- | -- | 2.54 | -- | -- |
| C5 | Success | 2-46, 2-32 | -- | VB 4.5kb | 5.12E+05 | 4.11 | 5.63E+07 | 2.88E+13 |
| C6 | Success | 2-46, 2-32 | -- | VB 4.5kb | 5.12E+05 | 3.99 | 5.25E+07 | 2.69E+13 |
| C7 | Success | 2-25, 2-32 | 1.41E+08 | -- | -- | 2.39 | -- | -- |
| C8 | Success | 2-25, 2-32 | 1.41E+08 | -- | -- | 2.38 | -- | -- |
| C9 | Excluded | 2-25, 2-32 | -- | VB 4.5kb | 1.6E+04 | 1.63 | LOW | -- |
| C10 | Excluded | 2-25, 2-32 | -- | VB 4.5kb | 1.6E+04 | 1.8 | 8.15E+07 | 1.3E+12 |
| D3 | Success | 2-46, 2-32 | 3.53E+07 | -- | -- | 3.41 | -- | -- |
| D4 | Success | 2-46, 2-32 | 3.53E+07 | -- | -- | 3.25 | -- | -- |
| D5 | Success | 2-46, 2-32 | -- | VB 4.5kb | 2.56E+05 | 5.49 | 1.12E+08 | 2.86E+13 |
| D6 | Success | 2-46, 2-32 | -- | VB 4.5kb | 2.56E+05 | 5.91 | 1.34E+08 | 3.43E+13 |
| D7 | Success | 2-25, 2-32 | 2.82E+08 | -- | -- | 3.29 | -- | -- |
| D8 | Success | 2-25, 2-32 | 2.82E+08 | -- | -- | 3.53 | -- | -- |
| D9 | Excluded | 2-25, 2-32 | -- | VB 4.5kb | 8E+03 | 2.21 | 1.2E+08 | 9.6E+11 |
| D10 | Excluded | 2-25, 2-32 | -- | VB 4.5kb | 8E+03 | 2.41 | 1.4E+08 | 1.12E+12 |
| E3 | Success | 2-46, 2-32 | 7.06E+07 | -- | -- | 4.73 | -- | -- |
| E4 | Success | 2-46, 2-32 | 7.06E+07 | -- | -- | 4.24 | -- | -- |
| E5 | Success | 2-46, 2-32 | -- | VB 4.5kb | 1.28E+05 | 7.66 | 2.64E+08 | 3.38E+13 |

| Sensor | Status | Design | | | | Result | | |
|---|---|---|---|---|---|---|---|---|
| | | Target | Concentration | Sample ID | Dilution Factor | Nano Units | Measured Conc. | Undiluted Conc. |
| E6 | Success | 2-46, 2-32 | -- | VB 4.5kb | 1.28E+05 | 7.67 | 2.65E+08 | 3.4E+13 |
| E7 | Success | 2-25, 2-32 | 5.65E+08 | -- | -- | 5.03 | -- | -- |
| E8 | Success | 2-25, 2-32 | 5.65E+08 | -- | -- | 5.37 | -- | -- |
| E9 | Success | 2-25, 2-32 | -- | VB 4.5kb | 4E+03 | 3.52 | 2.81E+08 | 1.12E+12 |
| E10 | Success | 2-25, 2-32 | -- | VB 4.5kb | 4E+03 | 3.67 | 3.05E+08 | 1.22E+12 |
| F3 | Success | 2-46, 2-32 | 1.41E+08 | -- | -- | 6.61 | -- | -- |
| F4 | Success | 2-46, 2-32 | 1.41E+08 | -- | -- | 5.78 | -- | -- |
| F5 | Excluded | 2-46, 2-32 | -- | VB 4.5kb | 6.4E+04 | 9.59 | 5.16E+08 | 3.3E+13 |
| F6 | Excluded | 2-46, 2-32 | -- | VB 4.5kb | 6.4E+04 | 9.4 | 4.84E+08 | 3.1E+13 |
| F7 | Success | 2-25, 2-32 | 1.13E+09 | -- | -- | 6.06 | -- | -- |
| F8 | Success | 2-25, 2-32 | 1.13E+09 | -- | -- | 6 | -- | -- |
| F9 | Success | 2-25, 2-32 | -- | VB 4.5kb | 2E+03 | 4.82 | 5.46E+08 | 1.09E+12 |
| F10 | Success | 2-25, 2-32 | -- | VB 4.5kb | 2E+03 | 4.61 | 4.92E+08 | 9.83E+11 |
| G3 | Success | 2-46, 2-32 | 2.82E+08 | -- | -- | 7.54 | -- | -- |
| G4 | Success | 2-46, 2-32 | 2.82E+08 | -- | -- | 7.88 | -- | -- |
| G5 | Excluded | 2-46, 2-32 | -- | VB 4.5kb | 3.2E+04 | 11 | HIGH | -- |
| G6 | Excluded | 2-46, 2-32 | -- | VB 4.5kb | 3.2E+04 | 10.3 | HIGH | -- |
| G7 | Success | 2-25, 2-32 | 2.26E+09 | -- | -- | 8.35 | -- | -- |

| Sensor | Status | Design | | | | Result | | |
|---|---|---|---|---|---|---|---|---|
| | | Target | Concentration | Sample ID | Dilution Factor | Nano Units | Measured Conc. | Undiluted Conc. |
| G8 | Success | 2-25, 2-32 | 2.26E+09 | -- | -- | 7.75 | -- | -- |
| G9 | Success | 2-25, 2-32 | -- | VB 4.5kb | 1E+03 | 6.78 | 1.37E+09 | 1.37E+12 |
| G10 | Success | 2-25, 2-32 | -- | VB 4.5kb | 1E+03 | 6.86 | 1.42E+09 | 1.42E+12 |
| H3 | Success | 2-46, 2-32 | 5.65E+08 | -- | -- | 10.7 | -- | -- |
| H4 | Success | 2-46, 2-32 | 5.65E+08 | -- | -- | 9.09 | -- | -- |
| H5 | Excluded | 2-46, 2-32 | -- | VB 4.5kb | 1.6E+04 | 10.2 | HIGH | -- |
| H6 | Excluded | 2-46, 2-32 | -- | VB 4.5kb | 1.6E+04 | 11.2 | HIGH | -- |
| H7 | Success | 2-25, 2-32 | 4.52E+09 | -- | -- | 9.56 | -- | -- |
| H8 | Success | 2-25, 2-32 | 4.52E+09 | -- | -- | 9.01 | -- | -- |
| H9 | Excluded | 2-25, 2-32 | -- | VB 4.5kb | 500 | 8.68 | 3.32E+09 | 1.66E+12 |
| H10 | Excluded | 2-25, 2-32 | -- | VB 4.5kb | 500 | 8.61 | 3.21E+09 | 1.61E+12 |

### Ignored Wells

| | 1 | 2 | 3 | 4 | 5 | 6 | 7 | 8 | 9 | 10 | 11 | 12 |
|---|---|---|---|---|---|---|---|---|---|---|---|---|
| A | A1 | A2 | | | | | | | | | A11 | A12 |
| B | B1 | B2 | | | | | | | | | B11 | B12 |
| C | C1 | C2 | | | | | | | | | C11 | C12 |
| D | D1 | D2 | | | | | | | | | D11 | D12 |
| E | E1 | E2 | | | | | | | | | E11 | E12 |
| F | F1 | F2 | | | | | | | | | F11 | F12 |
| G | G1 | G2 | | | | | | | | | G11 | G12 |
| H | H1 | H2 | | | | | | | | | H11 | H12 |
