## Supplementary File S3 for "Nanoneedle-Enabled Quantification of rAAV9 Capsid and Genome Integrity Reveals a Truncation Hotspot Locus in a 4.5 kb Transgene": 20250408_VB_probe_walk_experiment_T30_2-42_2-32_and_2-44_2-32.pdf

### Tessie Analysis Report

#### 96 well, 1 plex

| | |
|---|---|
| Plate Type: | AA01 |
| Assay Design: | 20250408_VB_probe_walk_experiment_T30_2-42_2-32_and_2-44_2-32 |
| Scan Package: | 00006920_T1030_1_1 |
| Instrument ID: | T1030 |
| Plate ID: | 00006920 |
| Pre Scan User: | N/A |
| Pre Scan Number: | 1 |
| Pre Scan Time: | 2025-04-08 13:08:51-04:00 |
| Post Scan User: | N/A |
| Post Scan Number: | 1 |
| Post Scan Time: | 2025-04-09 14:30:51-04:00 |
| Image Analysis Version: | 1.1.0.0 |
| Report Generation Time: | 2025-04-09 15:36:43-04:00 |
| Markers: | 2-42, 2-32, 2-44, 2-32 |

### NanoUnits Heatmap

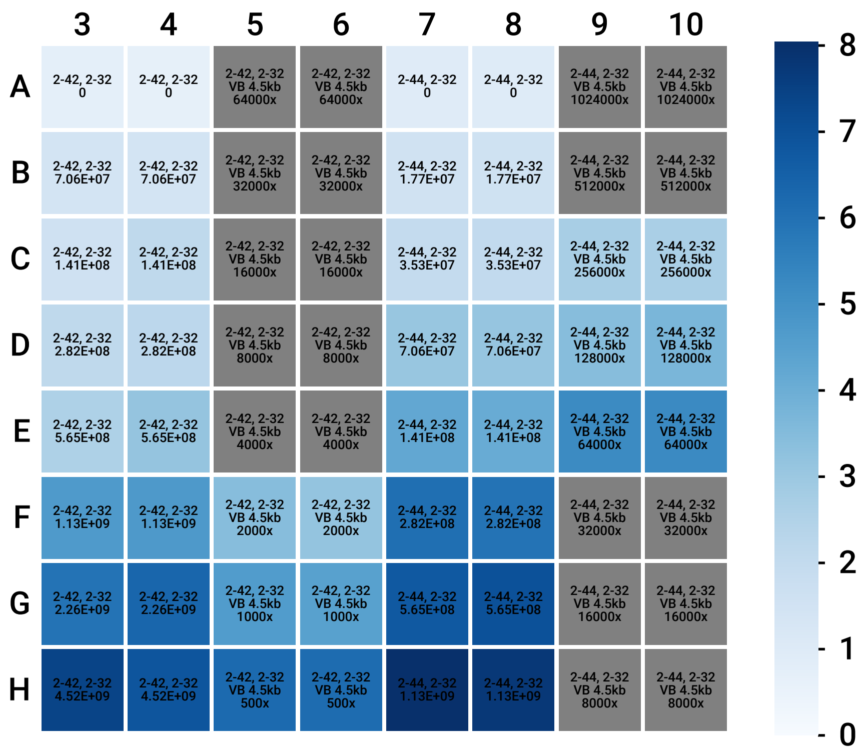

### 2-42, 2-32 Standard Curve

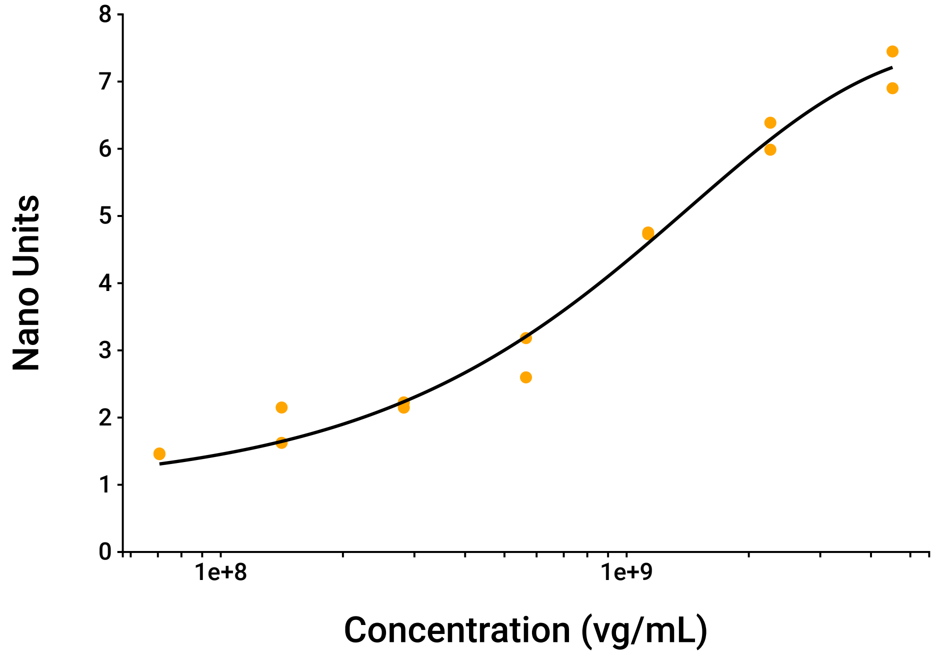


### 2-44, 2-32 Standard Curve

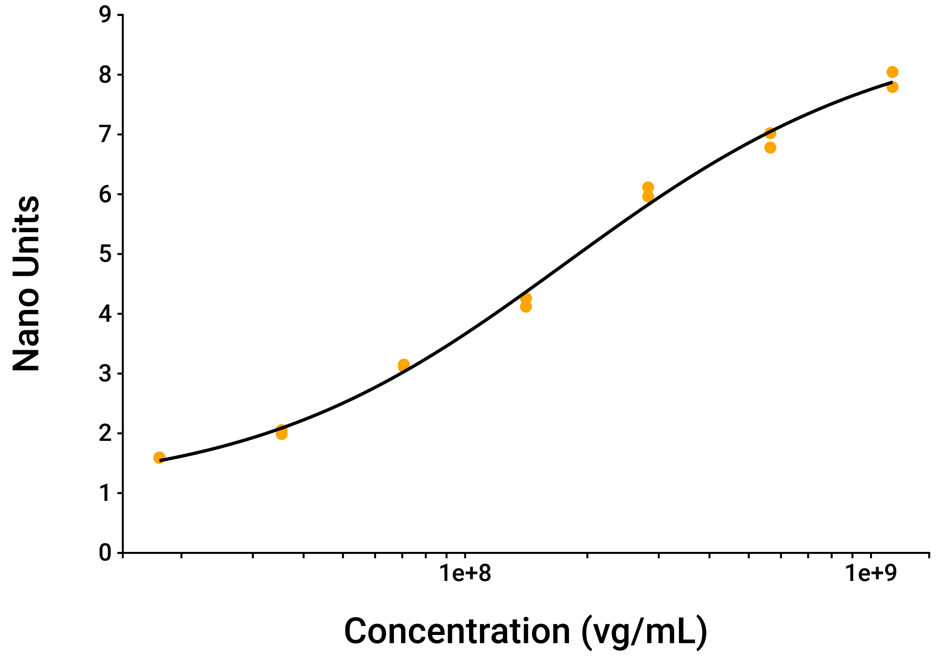

### VB 4.5kb Dilutions Data

| Target | Dilution Factor | Measured Concentration | Undiluted Concentration | %CV | Linearity %CV |
|---|---|---|---|---|---|
| 2-42, 2-32 (vg/mL) | 500 | 2.36E+09 | 1.18E+12 | 2.56 | 3.98 |
| | 1000 | 1.13E+09 | 1.13E+12 | 5.21 | |
| | 2000 | 6.09E+08 | 1.22E+12 | 10.9 | |
| 2-44, 2-32 (vg/mL) | 64000 | 2.14E+08 | 1.37E+13 | 0.60 | 8.82 |
| | 128000 | 9.73E+07 | 1.25E+13 | 10.4 | |
| | 256000 | 5.81E+07 | 1.49E+13 | 1.89 | |

### 2-42, 2-32 Standard Data

| Concentration (vg/mL) | Nano Units | | |
|---|---|---|---|
| | Mean | Standard Deviation | %CV |
| 0 | 0.735 | 0.0544 | 7.40 |
| 7.06E+07 | 1.46 | 0.00714 | 0.49 |
| 1.41E+08 | 1.89 | 0.372 | 19.7 |
| 2.82E+08 | 2.19 | 0.0552 | 2.53 |
| 5.65E+08 | 2.89 | 0.413 | 14.3 |
| 1.13E+09 | 4.74 | 0.0183 | 0.39 |
| 2.26E+09 | 6.19 | 0.284 | 4.59 |
| 4.52E+09 | 7.17 | 0.387 | 5.39 |

### 2-42, 2-32 Sample Data

| Sample Label | Concentration (vg/mL) | | |
|---|---|---|---|
| | Mean | Standard Deviation | %CV |
| VB 4.5kb | 1.18E+12 | 7.85E+10 | 6.68 |

### 2-44, 2-32 Standard Data

| Concentration (vg/mL) | Nano Units | | |
|---|---|---|---|
| | Mean | Standard Deviation | %CV |
| 0 | 1 | 0.0142 | 1.42 |
| 1.77E+07 | 1.59 | 0.00405 | 0.25 |
| 3.53E+07 | 2.02 | 0.0449 | 2.22 |
| 7.06E+07 | 3.13 | 0.0264 | 0.84 |
| 1.41E+08 | 4.19 | 0.098 | 2.34 |
| 2.82E+08 | 6.04 | 0.108 | 1.78 |
| 5.65E+08 | 6.9 | 0.17 | 2.47 |
| 1.13E+09 | 7.92 | 0.177 | 2.23 |

### 2-44, 2-32 Sample Data

| Sample Label | Concentration (vg/mL) | | |
|---|---|---|---|
| | Mean | Standard Deviation | %CV |
| VB 4.5kb | 1.37E+13 | 1.23E+12 | 9.01 |

### Curve Parameters

| Target | Curve | Equation | Parameters | $R^2$ |
|---|---|---|---|---|
| 2-42, 2-32 | Five parameter logisitic curve | $y = D - \frac{(A-D)}{\left[1+(x/C)^B\right]^E}$ | A = 0.92<br>B = 0.934<br>C = 6.2E+14<br>D = 7.56<br>E = 1.85E+05 | 0.987 |
| 2-44, 2-32 | Five parameter logisitic curve | $y = D - \frac{(A-D)}{\left[1+(x/C)^B\right]^E}$ | A = 1.02<br>B = 1.15<br>C = 1.51E+08<br>D = 8.93<br>E = 0.841 | 0.997 |

### Raw Data

| Sensor | Status | Design | | | | Result | | |
|---|---|---|---|---|---|---|---|---|
| | | Target | Concentration | Sample ID | Dilution Factor | Nano Units | Measured Conc. | Undiluted Conc. |
| A3 | Success | 2-42, 2-32 | 0 | -- | -- | 0.774 | -- | -- |
| A4 | Success | 2-42, 2-32 | 0 | -- | -- | 0.697 | -- | -- |
| A5 | Excluded | 2-42, 2-32 | -- | VB 4.5kb | 6.4E+04 | 0.993 | LOW | -- |
| A6 | Excluded | 2-42, 2-32 | -- | VB 4.5kb | 6.4E+04 | 1.09 | LOW | -- |
| A7 | Success | 2-44, 2-32 | 0 | -- | -- | 0.991 | -- | -- |
| A8 | Success | 2-44, 2-32 | 0 | -- | -- | 1.01 | -- | -- |
| A9 | Excluded | 2-44, 2-32 | -- | VB 4.5kb | 1.02E+06 | 1.61 | 1.98E+07 | 2.03E+13 |
| A10 | Excluded | 2-44, 2-32 | -- | VB 4.5kb | 1.02E+06 | 1.52 | LOW | -- |
| B3 | Success | 2-42, 2-32 | 7.06E+07 | -- | -- | 1.46 | -- | -- |
| B4 | Success | 2-42, 2-32 | 7.06E+07 | -- | -- | 1.47 | -- | -- |
| B5 | Excluded | 2-42, 2-32 | -- | VB 4.5kb | 3.2E+04 | 0.967 | LOW | -- |
| B6 | Excluded | 2-42, 2-32 | -- | VB 4.5kb | 3.2E+04 | 1.05 | LOW | -- |
| B7 | Success | 2-44, 2-32 | 1.77E+07 | -- | -- | 1.59 | -- | -- |
| B8 | Success | 2-44, 2-32 | 1.77E+07 | -- | -- | 1.6 | -- | -- |
| B9 | Excluded | 2-44, 2-32 | -- | VB 4.5kb | 5.12E+05 | 2.23 | 4.02E+07 | 2.06E+13 |
| B10 | Excluded | 2-44, 2-32 | -- | VB 4.5kb | 5.12E+05 | 2.21 | 3.96E+07 | 2.03E+13 |
| C3 | Success | 2-42, 2-32 | 1.41E+08 | -- | -- | 1.62 | -- | -- |

| Sensor | Status | Design | | | | Result | | |
|---|---|---|---|---|---|---|---|---|
| | | Target | Concentration | Sample ID | Dilution Factor | Nano Units | Measured Conc. | Undiluted Conc. |
| C4 | Success | 2-42, 2-32 | 1.41E+08 | -- | -- | 2.15 | -- | -- |
| C5 | Excluded | 2-42, 2-32 | -- | VB 4.5kb | 1.6E+04 | 1.35 | 7.84E+07 | 1.25E+12 |
| C6 | Excluded | 2-42, 2-32 | -- | VB 4.5kb | 1.6E+04 | 1.31 | 7.13E+07 | 1.14E+12 |
| C7 | Success | 2-44, 2-32 | 3.53E+07 | -- | -- | 1.99 | -- | -- |
| C8 | Success | 2-44, 2-32 | 3.53E+07 | -- | -- | 2.05 | -- | -- |
| C9 | Success | 2-44, 2-32 | -- | VB 4.5kb | 2.56E+05 | 2.74 | 5.88E+07 | 1.51E+13 |
| C10 | Success | 2-44, 2-32 | -- | VB 4.5kb | 2.56E+05 | 2.7 | 5.73E+07 | 1.47E+13 |
| D3 | Success | 2-42, 2-32 | 2.82E+08 | -- | -- | 2.15 | -- | -- |
| D4 | Success | 2-42, 2-32 | 2.82E+08 | -- | -- | 2.23 | -- | -- |
| D5 | Excluded | 2-42, 2-32 | -- | VB 4.5kb | 8E+03 | 1.7 | 1.53E+08 | 1.23E+12 |
| D6 | Excluded | 2-42, 2-32 | -- | VB 4.5kb | 8E+03 | 2.05 | 2.35E+08 | 1.88E+12 |
| D7 | Success | 2-44, 2-32 | 7.06E+07 | -- | -- | 3.11 | -- | -- |
| D8 | Success | 2-44, 2-32 | 7.06E+07 | -- | -- | 3.15 | -- | -- |
| D9 | Success | 2-44, 2-32 | -- | VB 4.5kb | 1.28E+05 | 3.46 | 9.01E+07 | 1.15E+13 |
| D10 | Success | 2-44, 2-32 | -- | VB 4.5kb | 1.28E+05 | 3.75 | 1.04E+08 | 1.34E+13 |
| E3 | Success | 2-42, 2-32 | 5.65E+08 | -- | -- | 2.6 | -- | -- |
| E4 | Success | 2-42, 2-32 | 5.65E+08 | -- | -- | 3.18 | -- | -- |
| E5 | Excluded | 2-42, 2-32 | -- | VB 4.5kb | 4E+03 | 2.11 | 2.5E+08 | 1E+12 |

| Sensor | Status | Design | | | | Result | | |
|---|---|---|---|---|---|---|---|---|
| | | Target | Concentration | Sample ID | Dilution Factor | Nano Units | Measured Conc. | Undiluted Conc. |
| E6 | Excluded | 2-42, 2-32 | -- | VB 4.5kb | 4E+03 | 2.13 | 2.56E+08 | 1.03E+12 |
| E7 | Success | 2-44, 2-32 | 1.41E+08 | -- | -- | 4.26 | -- | -- |
| E8 | Success | 2-44, 2-32 | 1.41E+08 | -- | -- | 4.12 | -- | -- |
| E9 | Success | 2-44, 2-32 | -- | VB 4.5kb | 6.4E+04 | 5.24 | 2.13E+08 | 1.36E+13 |
| E10 | Success | 2-44, 2-32 | -- | VB 4.5kb | 6.4E+04 | 5.26 | 2.15E+08 | 1.37E+13 |
| F3 | Success | 2-42, 2-32 | 1.13E+09 | -- | -- | 4.73 | -- | -- |
| F4 | Success | 2-42, 2-32 | 1.13E+09 | -- | -- | 4.75 | -- | -- |
| F5 | Success | 2-42, 2-32 | -- | VB 4.5kb | 2E+03 | 3.47 | 6.56E+08 | 1.31E+12 |
| F6 | Success | 2-42, 2-32 | -- | VB 4.5kb | 2E+03 | 3.2 | 5.62E+08 | 1.12E+12 |
| F7 | Success | 2-44, 2-32 | 2.82E+08 | -- | -- | 6.11 | -- | -- |
| F8 | Success | 2-44, 2-32 | 2.82E+08 | -- | -- | 5.96 | -- | -- |
| F9 | Excluded | 2-44, 2-32 | -- | VB 4.5kb | 3.2E+04 | 6.23 | 3.47E+08 | 1.11E+13 |
| F10 | Excluded | 2-44, 2-32 | -- | VB 4.5kb | 3.2E+04 | 6.95 | 5.3E+08 | 1.7E+13 |
| G3 | Success | 2-42, 2-32 | 2.26E+09 | -- | -- | 5.99 | -- | -- |
| G4 | Success | 2-42, 2-32 | 2.26E+09 | -- | -- | 6.39 | -- | -- |
| G5 | Success | 2-42, 2-32 | -- | VB 4.5kb | 1E+03 | 4.67 | 1.17E+09 | 1.17E+12 |
| G6 | Success | 2-42, 2-32 | -- | VB 4.5kb | 1E+03 | 4.5 | 1.08E+09 | 1.08E+12 |
| G7 | Success | 2-44, 2-32 | 5.65E+08 | -- | -- | 6.78 | -- | -- |

| Sensor | Status | Design | | | | Result | | |
|---|---|---|---|---|---|---|---|---|
| | | Target | Concentration | Sample ID | Dilution Factor | Nano Units | Measured Conc. | Undiluted Conc. |
| G8 | Success | 2-44, 2-32 | 5.65E+08 | -- | -- | 7.02 | -- | -- |
| G9 | Excluded | 2-44, 2-32 | -- | VB 4.5kb | 1.6E+04 | 8.31 | HIGH | -- |
| G10 | Excluded | 2-44, 2-32 | -- | VB 4.5kb | 1.6E+04 | 8.35 | HIGH | -- |
| H3 | Success | 2-42, 2-32 | 4.52E+09 | -- | -- | 7.45 | -- | -- |
| H4 | Success | 2-42, 2-32 | 4.52E+09 | -- | -- | 6.9 | -- | -- |
| H5 | Success | 2-42, 2-32 | -- | VB 4.5kb | 500 | 6.27 | 2.41E+09 | 1.2E+12 |
| H6 | Success | 2-42, 2-32 | -- | VB 4.5kb | 500 | 6.19 | 2.32E+09 | 1.16E+12 |
| H7 | Success | 2-44, 2-32 | 1.13E+09 | -- | -- | 8.04 | -- | -- |
| H8 | Success | 2-44, 2-32 | 1.13E+09 | -- | -- | 7.79 | -- | -- |
| H9 | Excluded | 2-44, 2-32 | -- | VB 4.5kb | 8E+03 | 9.06 | HIGH | -- |
| H10 | Excluded | 2-44, 2-32 | -- | VB 4.5kb | 8E+03 | 9.48 | HIGH | -- |

### Ignored Wells

| | 1 | 2 | 3 | 4 | 5 | 6 | 7 | 8 | 9 | 10 | 11 | 12 |
|---|---|---|---|---|---|---|---|---|---|---|---|---|
| A | A1 | A2 | | | | | | | | | A11 | A12 |
| B | B1 | B2 | | | | | | | | | B11 | B12 |
| C | C1 | C2 | | | | | | | | | C11 | C12 |
| D | D1 | D2 | | | | | | | | | D11 | D12 |
| E | E1 | E2 | | | | | | | | | E11 | E12 |
| F | F1 | F2 | | | | | | | | | F11 | F12 |
| G | G1 | G2 | | | | | | | | | G11 | G12 |
| H | H1 | H2 | | | | | | | | | H11 | H12 |
