## Supplementary File S3 for "Nanoneedle-Enabled Quantification of rAAV9 Capsid and Genome Integrity Reveals a Truncation Hotspot Locus in a 4.5 kb Transgene": 20250408_VB_probe_walk_experiment_T39_2-25_and_2-47_2-48_2-49.pdf

### Tessie Analysis Report

#### 96 well, 1 plex

| | |
|---|---|
| Plate Type: | AA01 |
| Assay Design: | 20250408_VB_probe_walk_experiment_T39_2-25_and_2-47_2-48_2-49 |
| Scan Package: | T1039_00006919_1_1 |
| Instrument ID: | T1039 |
| Plate ID: | 00006919 |
| Pre Scan User: | N/A |
| Pre Scan Number: | 1 |
| Pre Scan Time: | 2025-04-08 13:08:58-04:00 |
| Post Scan User: | N/A |
| Post Scan Number: | 1 |
| Post Scan Time: | 2025-04-09 14:30:24-04:00 |
| Image Analysis Version: | 1.1.0.0 |
| Report Generation Time: | 2025-04-09 16:03:26-04:00 |
| Markers: | 2-25, 2-47, 2-25, 2-48, 2-25, 2-49 |

### NanoUnits Heatmap

| | 1 | 2 | 3 | 4 | 5 | 6 | 7 | 8 | 9 | 10 | 11 | 12 |
|---|---|---|---|---|---|---|---|---|---|---|---|---|
| A | 2-25, 2-47<br>0 | 2-25, 2-47<br>0 | 2-25, 2-47<br>VB 4.5kb<br>1024000x | 2-25, 2-47<br>VB 4.5kb<br>1024000x | 2-25, 2-48<br>0 | 2-25, 2-48<br>0 | 2-25, 2-48<br>VB 4.5kb<br>256000x | 2-25, 2-48<br>VB 4.5kb<br>256000x | 2-25, 2-49<br>0 | 2-25, 2-49<br>0 | 2-25, 2-49<br>VB 4.5kb<br>256000x | 2-25, 2-49<br>VB 4.5kb<br>256000x |
| B | 2-25, 2-47<br>3.53E+07 | 2-25, 2-47<br>3.53E+07 | 2-25, 2-47<br>VB 4.5kb<br>512000x | 2-25, 2-47<br>VB 4.5kb<br>512000x | 2-25, 2-48<br>8.83E+06 | 2-25, 2-48<br>8.83E+06 | 2-25, 2-48<br>VB 4.5kb<br>128000x | 2-25, 2-48<br>VB 4.5kb<br>128000x | 2-25, 2-49<br>1.77E+07 | 2-25, 2-49<br>1.77E+07 | 2-25, 2-49<br>VB 4.5kb<br>128000x | 2-25, 2-49<br>VB 4.5kb<br>128000x |
| C | 2-25, 2-47<br>7.06E+07 | 2-25, 2-47<br>7.06E+07 | 2-25, 2-47<br>VB 4.5kb<br>256000x | 2-25, 2-47<br>VB 4.5kb<br>256000x | 2-25, 2-48<br>1.77E+07 | 2-25, 2-48<br>1.77E+07 | 2-25, 2-48<br>VB 4.5kb<br>64000x | 2-25, 2-48<br>VB 4.5kb<br>64000x | 2-25, 2-49<br>3.53E+07 | 2-25, 2-49<br>3.53E+07 | 2-25, 2-49<br>VB 4.5kb<br>64000x | 2-25, 2-49<br>VB 4.5kb<br>64000x |
| D | 2-25, 2-47<br>1.41E+08 | 2-25, 2-47<br>1.41E+08 | 2-25, 2-47<br>VB 4.5kb<br>128000x | 2-25, 2-47<br>VB 4.5kb<br>128000x | 2-25, 2-48<br>3.53E+07 | 2-25, 2-48<br>3.53E+07 | 2-25, 2-48<br>VB 4.5kb<br>32000x | 2-25, 2-48<br>VB 4.5kb<br>32000x | 2-25, 2-49<br>7.06E+07 | 2-25, 2-49<br>7.06E+07 | 2-25, 2-49<br>VB 4.5kb<br>32000x | 2-25, 2-49<br>VB 4.5kb<br>32000x |
| E | 2-25, 2-47<br>2.82E+08 | 2-25, 2-47<br>2.82E+08 | 2-25, 2-47<br>VB 4.5kb<br>64000x | 2-25, 2-47<br>VB 4.5kb<br>64000x | 2-25, 2-48<br>7.06E+07 | 2-25, 2-48<br>7.06E+07 | 2-25, 2-48<br>VB 4.5kb<br>16000x | 2-25, 2-48<br>VB 4.5kb<br>16000x | 2-25, 2-49<br>1.41E+08 | 2-25, 2-49<br>1.41E+08 | 2-25, 2-49<br>VB 4.5kb<br>16000x | 2-25, 2-49<br>VB 4.5kb<br>16000x |
| F | 2-25, 2-47<br>5.65E+08 | 2-25, 2-47<br>5.65E+08 | 2-25, 2-47<br>VB 4.5kb<br>32000x | 2-25, 2-47<br>VB 4.5kb<br>32000x | 2-25, 2-48<br>1.41E+08 | 2-25, 2-48<br>1.41E+08 | 2-25, 2-48<br>VB 4.5kb<br>8000x | 2-25, 2-48<br>VB 4.5kb<br>8000x | 2-25, 2-49<br>2.82E+08 | 2-25, 2-49<br>2.82E+08 | 2-25, 2-49<br>VB 4.5kb<br>8000x | 2-25, 2-49<br>VB 4.5kb<br>8000x |
| G | 2-25, 2-47<br>1.13E+09 | 2-25, 2-47<br>1.13E+09 | 2-25, 2-47<br>VB 4.5kb<br>16000x | 2-25, 2-47<br>VB 4.5kb<br>16000x | 2-25, 2-48<br>2.82E+08 | 2-25, 2-48<br>2.82E+08 | 2-25, 2-48<br>VB 4.5kb<br>4000x | 2-25, 2-48<br>VB 4.5kb<br>4000x | 2-25, 2-49<br>5.65E+08 | 2-25, 2-49<br>5.65E+08 | 2-25, 2-49<br>VB 4.5kb<br>4000x | 2-25, 2-49<br>VB 4.5kb<br>4000x |
| H | 2-25, 2-47<br>2.26E+09 | 2-25, 2-47<br>2.26E+09 | 2-25, 2-47<br>VB 4.5kb<br>8000x | 2-25, 2-47<br>VB 4.5kb<br>8000x | 2-25, 2-48<br>5.65E+08 | 2-25, 2-48<br>5.65E+08 | 2-25, 2-48<br>VB 4.5kb<br>2000x | 2-25, 2-48<br>VB 4.5kb<br>2000x | 2-25, 2-49<br>1.13E+09 | 2-25, 2-49<br>1.13E+09 | 2-25, 2-49<br>VB 4.5kb<br>2000x | 2-25, 2-49<br>VB 4.5kb<br>2000x |

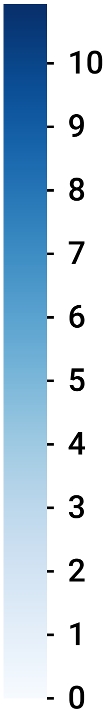

### 2-25, 2-47 Standard Curve

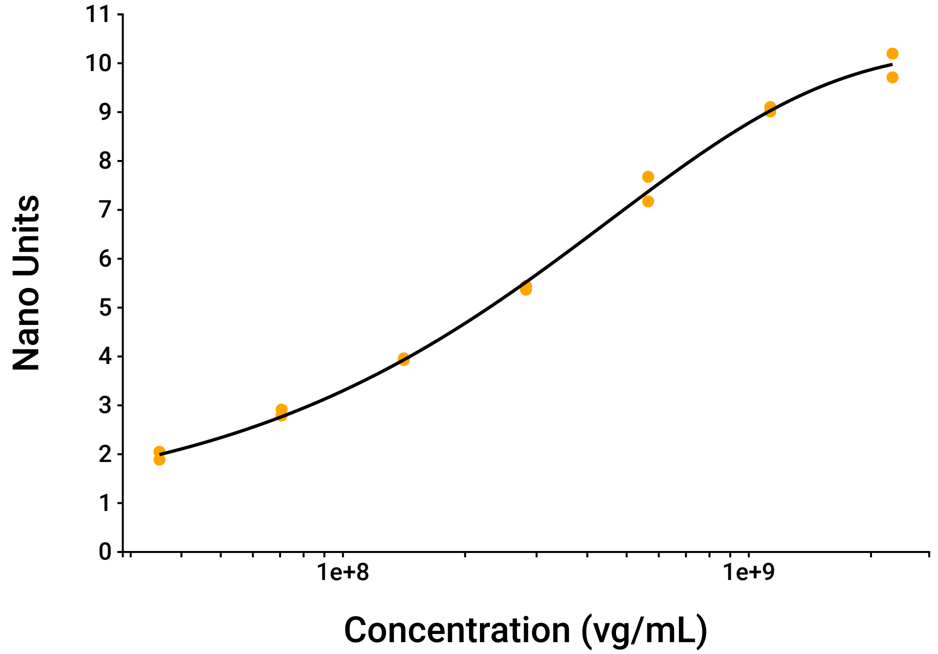


### 2-25, 2-48 Standard Curve

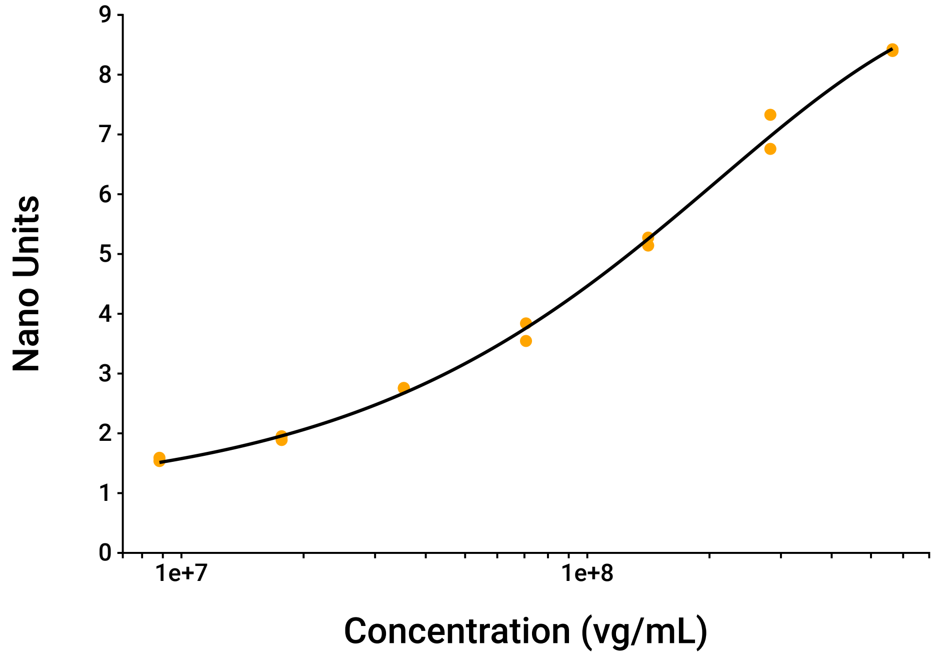

### 2-25, 2-49 Standard Curve

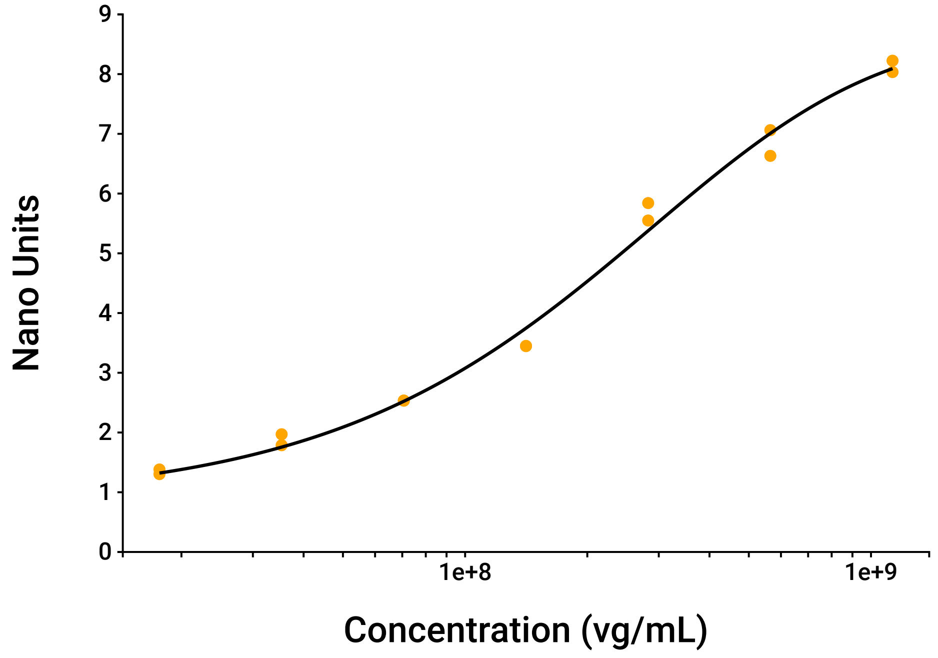


### VB 4.5kb Dilutions Data

| Target | Dilution Factor | Measured Concentration | Undiluted Concentration | %CV | Linearity %CV |
|---|---|---|---|---|---|
| 2-25, 2-47 (vg/mL) | 32000 | 1.06E+09 | 3.39E+13 | 12.3 | 13.1 |
| | 64000 | 4.53E+08 | 2.9E+13 | 4.21 | |
| | 128000 | 2.23E+08 | 2.86E+13 | 6.67 | |
| | 256000 | 1.46E+08 | 3.74E+13 | 7.92 | |
| 2-25, 2-48 (vg/mL) | 8000 | 2.32E+08 | 1.85E+12 | 5.55 | 5.24 |
| | 16000 | 1.08E+08 | 1.73E+12 | 8.52 | |
| | 32000 | 5.23E+07 | 1.67E+12 | 0.45 | |
| 2-25, 2-49 (vg/mL) | 2000 | 6.57E+08 | 1.31E+12 | 10.8 | 6.12 |
| | 4000 | 3.17E+08 | 1.27E+12 | 7.83 | |
| | 8000 | 1.68E+08 | 1.34E+12 | 1.73 | |
| | 16000 | 9.14E+07 | 1.46E+12 | 10.2 | |

### 2-25, 2-47 Standard Data

| Concentration (vg/mL) | Nano Units | | |
|---|---|---|---|
| | Mean | Standard Deviation | %CV |
| 0 | 0.755 | 0.0266 | 3.53 |
| 3.53E+07 | 1.97 | 0.114 | 5.80 |
| 7.06E+07 | 2.85 | 0.0847 | 2.97 |
| 1.41E+08 | 3.94 | 0.0217 | 0.55 |
| 2.82E+08 | 5.41 | 0.0543 | 1.00 |
| 5.65E+08 | 7.43 | 0.356 | 4.79 |
| 1.13E+09 | 9.06 | 0.0642 | 0.71 |
| 2.26E+09 | 9.95 | 0.343 | 3.44 |

### 2-25, 2-47 Sample Data

| Sample Label | Concentration (vg/mL) | | |
|---|---|---|---|
| | Mean | Standard Deviation | %CV |
| VB 4.5kb | 3.22E+13 | 4.45E+12 | 13.8 |

### 2-25, 2-48 Standard Data

| Concentration (vg/mL) | Nano Units | | |
|---|---|---|---|
| | Mean | Standard Deviation | %CV |
| 0 | 0.846 | 0.0319 | 3.77 |
| 8.83E+06 | 1.56 | 0.0383 | 2.45 |
| 1.77E+07 | 1.92 | 0.0444 | 2.31 |
| 3.53E+07 | 2.76 | 0.00498 | 0.18 |
| 7.06E+07 | 3.69 | 0.208 | 5.63 |
| 1.41E+08 | 5.21 | 0.0936 | 1.80 |
| 2.82E+08 | 7.04 | 0.403 | 5.72 |
| 5.65E+08 | 8.41 | 0.0201 | 0.24 |

### 2-25, 2-48 Sample Data

| Sample Label | Concentration (vg/mL) | | |
|---|---|---|---|
| | Mean | Standard Deviation | %CV |
| VB 4.5kb | 1.75E+12 | 1.15E+11 | 6.56 |

### 2-25, 2-49 Standard Data

| Concentration (vg/mL) | Nano Units | | |
|---|---|---|---|
| | Mean | Standard Deviation | %CV |
| 0 | 0.789 | 0.0278 | 3.53 |
| 1.77E+07 | 1.34 | 0.0552 | 4.11 |
| 3.53E+07 | 1.88 | 0.129 | 6.89 |
| 7.06E+07 | 2.53 | 0.000149 | 0.01 |
| 1.41E+08 | 3.45 | 0.00312 | 0.09 |
| 2.82E+08 | 5.7 | 0.206 | 3.62 |
| 5.65E+08 | 6.85 | 0.304 | 4.45 |
| 1.13E+09 | 8.13 | 0.135 | 1.66 |

### 2-25, 2-49 Sample Data

| Sample Label | Concentration (vg/mL) | | |
|---|---|---|---|
| | Mean | Standard Deviation | %CV |
| VB 4.5kb | 1.35E+12 | 1.15E+11 | 8.57 |

### Curve Parameters

| Target | Curve | Equation | Parameters | $R^2$ |
|---|---|---|---|---|
| 2-25, 2-47 | Five parameter logisitic curve | $y = D - \frac{(A-D)}{\left[1+(x/C)^B\right]^E}$ | A = 0.771<br>B = 0.777<br>C = 2.45E+15<br>D = 10.3<br>E = 1.72E+05 | 0.998 |
| 2-25, 2-48 | Five parameter logisitic curve | $y = D - \frac{(A-D)}{\left[1+(x/C)^B\right]^E}$ | A = 0.876<br>B = 0.804<br>C = 4.33E+14<br>D = 9.37<br>E = 1.19E+05 | 0.997 |
| 2-25, 2-49 | Five parameter logisitic curve | $y = D - \frac{(A-D)}{\left[1+(x/C)^B\right]^E}$ | A = 0.841<br>B = 0.985<br>C = 1.05E+09<br>D = 8.67<br>E = 3.57 | 0.994 |

### Raw Data

| Sensor | Status | Design | | | | Result | | |
|---|---|---|---|---|---|---|---|---|
| | | Target | Concentration | Sample ID | Dilution Factor | Nano Units | Measured Conc. | Undiluted Conc. |
| A1 | Success | 2-25, 2-47 | 0 | -- | -- | 0.736 | -- | -- |
| A2 | Success | 2-25, 2-47 | 0 | -- | -- | 0.774 | -- | -- |
| A3 | Excluded | 2-25, 2-47 | -- | VB 4.5kb | 1.02E+06 | 2.12 | 4.05E+07 | 4.15E+13 |
| A4 | Excluded | 2-25, 2-47 | -- | VB 4.5kb | 1.02E+06 | 2.1 | 3.96E+07 | 4.05E+13 |
| A5 | Success | 2-25, 2-48 | 0 | -- | -- | 0.868 | -- | -- |
| A6 | Success | 2-25, 2-48 | 0 | -- | -- | 0.823 | -- | -- |
| A7 | Excluded | 2-25, 2-48 | -- | VB 4.5kb | 2.56E+05 | 1.29 | LOW | -- |
| A8 | Excluded | 2-25, 2-48 | -- | VB 4.5kb | 2.56E+05 | 1.37 | LOW | -- |
| A9 | Success | 2-25, 2-49 | 0 | -- | -- | 0.809 | -- | -- |
| A10 | Success | 2-25, 2-49 | 0 | -- | -- | 0.769 | -- | -- |
| A11 | Excluded | 2-25, 2-49 | -- | VB 4.5kb | 2.56E+05 | 1.02 | LOW | -- |
| A12 | Excluded | 2-25, 2-49 | -- | VB 4.5kb | 2.56E+05 | 1.05 | LOW | -- |
| B1 | Success | 2-25, 2-47 | 3.53E+07 | -- | -- | 2.05 | -- | -- |
| B2 | Success | 2-25, 2-47 | 3.53E+07 | -- | -- | 1.89 | -- | -- |
| B3 | Excluded | 2-25, 2-47 | -- | VB 4.5kb | 5.12E+05 | 3.06 | 8.64E+07 | 4.43E+13 |
| B4 | Excluded | 2-25, 2-47 | -- | VB 4.5kb | 5.12E+05 | 2.89 | 7.68E+07 | 3.93E+13 |
| B5 | Success | 2-25, 2-48 | 8.83E+06 | -- | -- | 1.54 | -- | -- |

| Sensor | Status | Design | | | | Result | | |
|---|---|---|---|---|---|---|---|---|
| | | Target | Concentration | Sample ID | Dilution Factor | Nano Units | Measured Conc. | Undiluted Conc. |
| B6 | Success | 2-25, 2-48 | 8.83E+06 | -- | -- | 1.59 | -- | -- |
| B7 | Excluded | 2-25, 2-48 | -- | VB 4.5kb | 1.28E+05 | 1.73 | 1.29E+07 | 1.66E+12 |
| B8 | Excluded | 2-25, 2-48 | -- | VB 4.5kb | 1.28E+05 | 1.78 | 1.4E+07 | 1.79E+12 |
| B9 | Success | 2-25, 2-49 | 1.77E+07 | -- | -- | 1.38 | -- | -- |
| B10 | Success | 2-25, 2-49 | 1.77E+07 | -- | -- | 1.3 | -- | -- |
| B11 | Excluded | 2-25, 2-49 | -- | VB 4.5kb | 1.28E+05 | 1.2 | LOW | -- |
| B12 | Excluded | 2-25, 2-49 | -- | VB 4.5kb | 1.28E+05 | 1.05 | LOW | -- |
| C1 | Success | 2-25, 2-47 | 7.06E+07 | -- | -- | 2.79 | -- | -- |
| C2 | Success | 2-25, 2-47 | 7.06E+07 | -- | -- | 2.91 | -- | -- |
| C3 | Success | 2-25, 2-47 | -- | VB 4.5kb | 2.56E+05 | 4.11 | 1.54E+08 | 3.95E+13 |
| C4 | Success | 2-25, 2-47 | -- | VB 4.5kb | 2.56E+05 | 3.89 | 1.38E+08 | 3.53E+13 |
| C5 | Success | 2-25, 2-48 | 1.77E+07 | -- | -- | 1.89 | -- | -- |
| C6 | Success | 2-25, 2-48 | 1.77E+07 | -- | -- | 1.95 | -- | -- |
| C7 | Excluded | 2-25, 2-48 | -- | VB 4.5kb | 6.4E+04 | 2.25 | 2.45E+07 | 1.57E+12 |
| C8 | Excluded | 2-25, 2-48 | -- | VB 4.5kb | 6.4E+04 | 2.37 | 2.73E+07 | 1.74E+12 |
| C9 | Success | 2-25, 2-49 | 3.53E+07 | -- | -- | 1.97 | -- | -- |
| C10 | Success | 2-25, 2-49 | 3.53E+07 | -- | -- | 1.79 | -- | -- |
| C11 | Excluded | 2-25, 2-49 | -- | VB 4.5kb | 6.4E+04 | 1.48 | 2.4E+07 | 1.54E+12 |

| Sensor | Status | Design | | | | Result | | |
|---|---|---|---|---|---|---|---|---|
| | | Target | Concentration | Sample ID | Dilution Factor | Nano Units | Measured Conc. | Undiluted Conc. |
| C12 | Excluded | 2-25, 2-49 | -- | VB 4.5kb | 6.4E+04 | 1.53 | 2.6E+07 | 1.66E+12 |
| D1 | Success | 2-25, 2-47 | 1.41E+08 | -- | -- | 3.93 | -- | -- |
| D2 | Success | 2-25, 2-47 | 1.41E+08 | -- | -- | 3.96 | -- | -- |
| D3 | Success | 2-25, 2-47 | -- | VB 4.5kb | 1.28E+05 | 4.82 | 2.13E+08 | 2.72E+13 |
| D4 | Success | 2-25, 2-47 | -- | VB 4.5kb | 1.28E+05 | 5.05 | 2.34E+08 | 2.99E+13 |
| D5 | Success | 2-25, 2-48 | 3.53E+07 | -- | -- | 2.75 | -- | -- |
| D6 | Success | 2-25, 2-48 | 3.53E+07 | -- | -- | 2.76 | -- | -- |
| D7 | Success | 2-25, 2-48 | -- | VB 4.5kb | 3.2E+04 | 3.23 | 5.21E+07 | 1.67E+12 |
| D8 | Success | 2-25, 2-48 | -- | VB 4.5kb | 3.2E+04 | 3.24 | 5.24E+07 | 1.68E+12 |
| D9 | Success | 2-25, 2-49 | 7.06E+07 | -- | -- | 2.53 | -- | -- |
| D10 | Success | 2-25, 2-49 | 7.06E+07 | -- | -- | 2.53 | -- | -- |
| D11 | Excluded | 2-25, 2-49 | -- | VB 4.5kb | 3.2E+04 | 1.99 | 4.55E+07 | 1.46E+12 |
| D12 | Excluded | 2-25, 2-49 | -- | VB 4.5kb | 3.2E+04 | 1.9 | 4.17E+07 | 1.33E+12 |
| E1 | Success | 2-25, 2-47 | 2.82E+08 | -- | -- | 5.44 | -- | -- |
| E2 | Success | 2-25, 2-47 | 2.82E+08 | -- | -- | 5.37 | -- | -- |
| E3 | Success | 2-25, 2-47 | -- | VB 4.5kb | 6.4E+04 | 6.86 | 4.66E+08 | 2.98E+13 |
| E4 | Success | 2-25, 2-47 | -- | VB 4.5kb | 6.4E+04 | 6.7 | 4.39E+08 | 2.81E+13 |
| E5 | Success | 2-25, 2-48 | 7.06E+07 | -- | -- | 3.84 | -- | -- |

| Sensor | Status | Design | | | | Result | | |
|---|---|---|---|---|---|---|---|---|
| | | Target | Concentration | Sample ID | Dilution Factor | Nano Units | Measured Conc. | Undiluted Conc. |
| E6 | Success | 2-25, 2-48 | 7.06E+07 | -- | -- | 3.55 | -- | -- |
| E7 | Success | 2-25, 2-48 | -- | VB 4.5kb | 1.6E+04 | 4.77 | 1.15E+08 | 1.84E+12 |
| E8 | Success | 2-25, 2-48 | -- | VB 4.5kb | 1.6E+04 | 4.5 | 1.02E+08 | 1.63E+12 |
| E9 | Success | 2-25, 2-49 | 1.41E+08 | -- | -- | 3.45 | -- | -- |
| E10 | Success | 2-25, 2-49 | 1.41E+08 | -- | -- | 3.45 | -- | -- |
| E11 | Success | 2-25, 2-49 | -- | VB 4.5kb | 1.6E+04 | 2.8 | 8.48E+07 | 1.36E+12 |
| E12 | Success | 2-25, 2-49 | -- | VB 4.5kb | 1.6E+04 | 3.04 | 9.8E+07 | 1.57E+12 |
| F1 | Success | 2-25, 2-47 | 5.65E+08 | -- | -- | 7.68 | -- | -- |
| F2 | Success | 2-25, 2-47 | 5.65E+08 | -- | -- | 7.17 | -- | -- |
| F3 | Success | 2-25, 2-47 | -- | VB 4.5kb | 3.2E+04 | 8.7 | 9.66E+08 | 3.09E+13 |
| F4 | Success | 2-25, 2-47 | -- | VB 4.5kb | 3.2E+04 | 9.06 | 1.15E+09 | 3.68E+13 |
| F5 | Success | 2-25, 2-48 | 1.41E+08 | -- | -- | 5.27 | -- | -- |
| F6 | Success | 2-25, 2-48 | 1.41E+08 | -- | -- | 5.14 | -- | -- |
| F7 | Success | 2-25, 2-48 | -- | VB 4.5kb | 8E+03 | 6.38 | 2.22E+08 | 1.78E+12 |
| F8 | Success | 2-25, 2-48 | -- | VB 4.5kb | 8E+03 | 6.57 | 2.41E+08 | 1.92E+12 |
| F9 | Success | 2-25, 2-49 | 2.82E+08 | -- | -- | 5.55 | -- | -- |
| F10 | Success | 2-25, 2-49 | 2.82E+08 | -- | -- | 5.84 | -- | -- |
| F11 | Success | 2-25, 2-49 | -- | VB 4.5kb | 8E+03 | 4.09 | 1.66E+08 | 1.32E+12 |

| Sensor | Status | Design | | | | Result | | |
|---|---|---|---|---|---|---|---|---|
| | | Target | Concentration | Sample ID | Dilution Factor | Nano Units | Measured Conc. | Undiluted Conc. |
| F12 | Success | 2-25, 2-49 | -- | VB 4.5kb | 8E+03 | 4.15 | 1.7E+08 | 1.36E+12 |
| G1 | Success | 2-25, 2-47 | 1.13E+09 | -- | -- | 9.01 | -- | -- |
| G2 | Success | 2-25, 2-47 | 1.13E+09 | -- | -- | 9.1 | -- | -- |
| G3 | Success | 2-25, 2-47 | -- | VB 4.5kb | 1.6E+04 | 9.27 | 1.29E+09 | 2.06E+13 |
| G4 | Success | 2-25, 2-47 | -- | VB 4.5kb | 1.6E+04 | 10.2 | HIGH | -- |
| G5 | Success | 2-25, 2-48 | 2.82E+08 | -- | -- | 7.33 | -- | -- |
| G6 | Success | 2-25, 2-48 | 2.82E+08 | -- | -- | 6.76 | -- | -- |
| G7 | Success | 2-25, 2-48 | -- | VB 4.5kb | 4E+03 | 7.94 | 4.34E+08 | 1.73E+12 |
| G8 | Success | 2-25, 2-48 | -- | VB 4.5kb | 4E+03 | 8.8 | HIGH | -- |
| G9 | Success | 2-25, 2-49 | 5.65E+08 | -- | -- | 7.06 | -- | -- |
| G10 | Success | 2-25, 2-49 | 5.65E+08 | -- | -- | 6.63 | -- | -- |
| G11 | Success | 2-25, 2-49 | -- | VB 4.5kb | 4E+03 | 5.53 | 3E+08 | 1.2E+12 |
| G12 | Success | 2-25, 2-49 | -- | VB 4.5kb | 4E+03 | 5.8 | 3.35E+08 | 1.34E+12 |
| H1 | Success | 2-25, 2-47 | 2.26E+09 | -- | -- | 9.71 | -- | -- |
| H2 | Success | 2-25, 2-47 | 2.26E+09 | -- | -- | 10.2 | -- | -- |
| H3 | Success | 2-25, 2-47 | -- | VB 4.5kb | 8E+03 | 9.76 | 1.81E+09 | 1.45E+13 |
| H4 | Success | 2-25, 2-47 | -- | VB 4.5kb | 8E+03 | 10.9 | HIGH | -- |
| H5 | Success | 2-25, 2-48 | 5.65E+08 | -- | -- | 8.42 | -- | -- |

| Sensor | Status | Design | | | | Result | | |
|---|---|---|---|---|---|---|---|---|
| | | Target | Concentration | Sample ID | Dilution Factor | Nano Units | Measured Conc. | Undiluted Conc. |
| H6 | Success | 2-25, 2-48 | 5.65E+08 | -- | -- | 8.4 | -- | -- |
| H7 | Success | 2-25, 2-48 | -- | VB 4.5kb | 2E+03 | 9.16 | HIGH | -- |
| H8 | Success | 2-25, 2-48 | -- | VB 4.5kb | 2E+03 | 9.6 | HIGH | -- |
| H9 | Success | 2-25, 2-49 | 1.13E+09 | -- | -- | 8.03 | -- | -- |
| H10 | Success | 2-25, 2-49 | 1.13E+09 | -- | -- | 8.22 | -- | -- |
| H11 | Success | 2-25, 2-49 | -- | VB 4.5kb | 2E+03 | 7.44 | 7.08E+08 | 1.42E+12 |
| H12 | Success | 2-25, 2-49 | -- | VB 4.5kb | 2E+03 | 7.15 | 6.07E+08 | 1.21E+12 |
