## Supplementary File S3 for "Nanoneedle-Enabled Quantification of rAAV9 Capsid and Genome Integrity Reveals a Truncation Hotspot Locus in a 4.5 kb Transgene": 20250418_AAV9_4.5kb_capsid_measurement.pdf

### Tessie Analysis Report

#### 96 well, 1 plex

| | |
|---|---|
| Plate Type: | AA01 |
| Assay Design: | 20250418_AAV9_4.5kb_capsid_measurement |
| Assay Design Date: | 2025-10-27 12:12:44-04:00 |
| Plate ID: | 00007169 |
| Scan Package: | 00007169_T1030_1_1 |
| Instrument ID: | T1030 |
| Pre Scan User: | N/A |
| Pre Scan Number: | 1 |
| Pre Scan Date: | 2025-04-17 16:37:02-04:00 |
| Post Scan User: | N/A |
| Post Scan Number: | 1 |
| Post Scan Date: | 2025-04-18 13:44:15-04:00 |
| Image Analysis Version: | 1.1.1.0 |
| Report Generation Date: | 2025-10-27 12:15:50-04:00 |
| Targets: | AAV9 |

### NanoUnits Heatmap

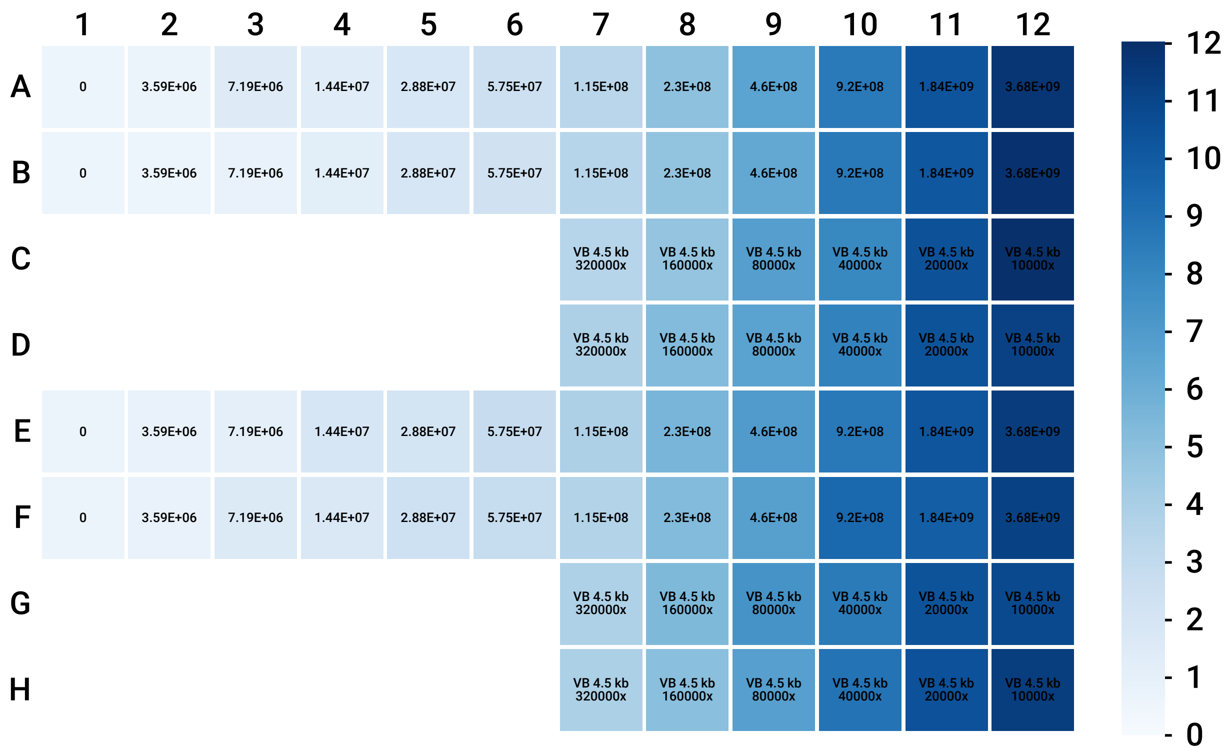

### AAV9 Standard Curve

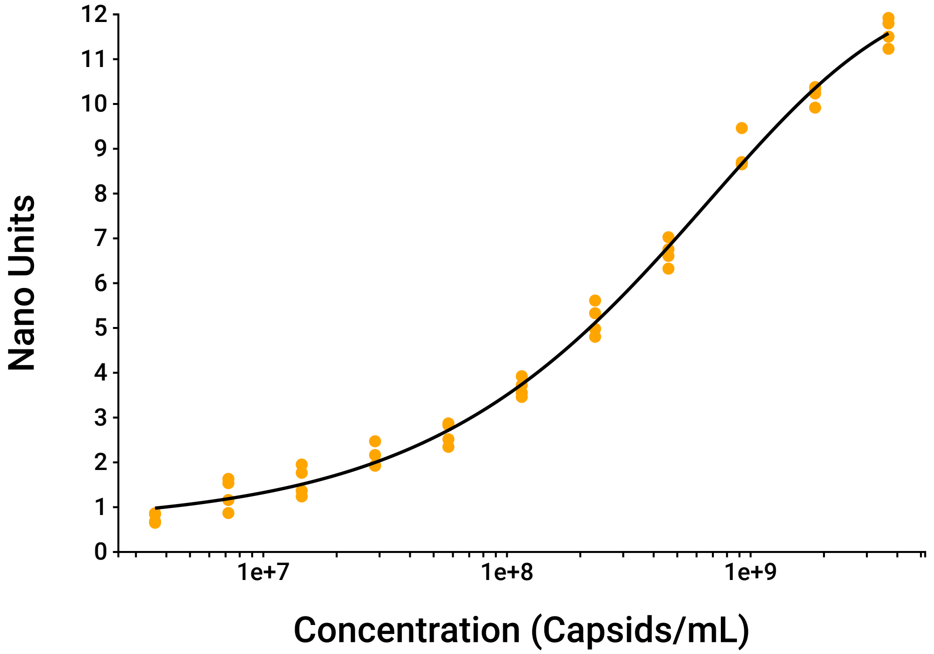


### AAV9 Standard Data

| Concentration (Capsids/mL) | Nano Units | | |
|---|---|---|---|
| | Mean | Standard Deviation | %CV |
| 0 | 0.635 | 0.0612 | 9.63 |
| 3.59E+06 | 0.76 | 0.108 | 14.3 |
| 7.19E+06 | 1.3 | 0.352 | 27.1 |
| 1.44E+07 | 1.58 | 0.334 | 21.1 |
| 2.88E+07 | 2.12 | 0.258 | 12.2 |
| 5.75E+07 | 2.64 | 0.252 | 9.54 |
| 1.15E+08 | 3.67 | 0.204 | 5.55 |
| 2.3E+08 | 5.18 | 0.361 | 6.97 |
| 4.6E+08 | 6.68 | 0.293 | 4.38 |
| 9.2E+08 | 8.87 | 0.393 | 4.42 |
| 1.84E+09 | 10.2 | 0.205 | 2.01 |
| 3.68E+09 | 11.6 | 0.309 | 2.66 |

### AAV9 Sample Data

| Sample Label | Concentration (Capsids/mL) | | |
|---|---|---|---|
| | Mean | Standard Deviation | %CV |
| VB 4.5 kb§ | 3.76E+13§ | 4.34E+12§ | 11.5§ |

§One or more dilutions excluded due to concentration being above upper limit

### VB 4.5 kb Dilutions Data

| Dilution Factor | Measured Concentration (Capsids/mL) | Undiluted Concentration (Capsids/mL) | %CV | Linearity %CV |
|---|---|---|---|---|
| 20000 | 1.94E+09 | 3.87E+13 | 1.63 | 5.34 |
| 40000 | 8.52E+08 | 3.41E+13 | 16.3 | |
| 80000 | 4.83E+08 | 3.87E+13 | 13.5 | |
| 160000 | 2.35E+08 | 3.76E+13 | 12.4 | |
| 320000 | 1.21E+08 | 3.88E+13 | 10.7 | |

### Curve Parameters

| Target | Curve | Equation | Parameters | $R^2$ |
|---|---|---|---|---|
| AAV9 | Five parameter logisitic curve | $y = D - \frac{(A-D)}{\left[1+(x/C)^B\right]^E}$ | A = 0.605<br>B = 0.658<br>C = 1.13E+10<br>D = 12.6<br>E = 6.37 | 0.995 |

### Raw Data

| Sensor | Status | Design | | | | Result | | |
|---|---|---|---|---|---|---|---|---|
| | | Target | Concentration | Sample ID | Dilution Factor | Nano Units | Measured Conc. | Undiluted Conc. |
| A1 | Success | AAV9 | 0 | -- | -- | 0.579 | -- | -- |
| A2 | Success | AAV9 | 3.59E+06 | -- | -- | 0.683 | -- | -- |
| A3 | Success | AAV9 | 7.19E+06 | -- | -- | 1.54 | -- | -- |
| A4 | Success | AAV9 | 1.44E+07 | -- | -- | 1.37 | -- | -- |
| A5 | Success | AAV9 | 2.88E+07 | -- | -- | 1.92 | -- | -- |
| A6 | Success | AAV9 | 5.75E+07 | -- | -- | 2.52 | -- | -- |
| A7 | Success | AAV9 | 1.15E+08 | -- | -- | 3.46 | -- | -- |
| A8 | Success | AAV9 | 2.3E+08 | -- | -- | 4.98 | -- | -- |
| A9 | Success | AAV9 | 4.6E+08 | -- | -- | 6.61 | -- | -- |
| A10 | Success | AAV9 | 9.2E+08 | -- | -- | 8.65 | -- | -- |
| A11 | Success | AAV9 | 1.84E+09 | -- | -- | 10.4 | -- | -- |
| A12 | Success | AAV9 | 3.68E+09 | -- | -- | 11.8 | -- | -- |
| B1 | Success | AAV9 | 0 | -- | -- | 0.586 | -- | -- |
| B2 | Success | AAV9 | 3.59E+06 | -- | -- | 0.652 | -- | -- |
| B3 | Success | AAV9 | 7.19E+06 | -- | -- | 0.869 | -- | -- |
| B4 | Success | AAV9 | 1.44E+07 | -- | -- | 1.24 | -- | -- |
| B5 | Success | AAV9 | 2.88E+07 | -- | -- | 1.93 | -- | -- |
| B6 | Success | AAV9 | 5.75E+07 | -- | -- | 2.35 | -- | -- |
| B7 | Success | AAV9 | 1.15E+08 | -- | -- | 3.55 | -- | -- |
| B8 | Success | AAV9 | 2.3E+08 | -- | -- | 4.8 | -- | -- |
| B9 | Success | AAV9 | 4.6E+08 | -- | -- | 6.32 | -- | -- |
| B10 | Success | AAV9 | 9.2E+08 | -- | -- | 8.68 | -- | -- |
| B11 | Success | AAV9 | 1.84E+09 | -- | -- | 10.2 | -- | -- |
| B12 | Success | AAV9 | 3.68E+09 | -- | -- | 11.9 | -- | -- |
| C7 | Success | AAV9 | -- | VB 4.5 kb | 3.2E+05 | 3.54 | 1.02E+08 | 3.26E+13 |
| C8 | Success | AAV9 | -- | VB 4.5 kb | 1.6E+05 | 4.78 | 1.97E+08 | 3.16E+13 |
| C9 | Success | AAV9 | -- | VB 4.5 kb | 8E+04 | 6.85 | 4.67E+08 | 3.74E+13 |
| C10 | Success | AAV9 | -- | VB 4.5 kb | 4E+04 | 7.94 | 7.01E+08 | 2.8E+13 |

| Sensor | Status | Design | | | | Result | | |
|---|---|---|---|---|---|---|---|---|
| | | Target | Concentration | Sample ID | Dilution Factor | Nano Units | Measured Conc. | Undiluted Conc. |
| C11 | Success | AAV9 | -- | VB 4.5 kb | 2E+04 | 10.5 | 1.95E+09 | 3.91E+13 |
| C12 | Success | AAV9 | -- | VB 4.5 kb | 1E+04 | 12 | HIGH | -- |
| D7 | Success | AAV9 | -- | VB 4.5 kb | 3.2E+05 | 3.94 | 1.29E+08 | 4.13E+13 |
| D8 | Success | AAV9 | -- | VB 4.5 kb | 1.6E+05 | 5.33 | 2.52E+08 | 4.04E+13 |
| D9 | Success | AAV9 | -- | VB 4.5 kb | 8E+04 | 6.64 | 4.3E+08 | 3.44E+13 |
| D10 | Success | AAV9 | -- | VB 4.5 kb | 4E+04 | 8.26 | 7.89E+08 | 3.16E+13 |
| D11 | Success | AAV9 | -- | VB 4.5 kb | 2E+04 | 10.4 | 1.91E+09 | 3.81E+13 |
| D12 | Success | AAV9 | -- | VB 4.5 kb | 1E+04 | 11.2 | 2.94E+09 | 2.94E+13 |
| E1 | Success | AAV9 | 0 | -- | -- | 0.693 | -- | -- |
| E2 | Success | AAV9 | 3.59E+06 | -- | -- | 0.842 | -- | -- |
| E3 | Success | AAV9 | 7.19E+06 | -- | -- | 1.16 | -- | -- |
| E4 | Success | AAV9 | 1.44E+07 | -- | -- | 1.95 | -- | -- |
| E5 | Success | AAV9 | 2.88E+07 | -- | -- | 2.16 | -- | -- |
| E6 | Success | AAV9 | 5.75E+07 | -- | -- | 2.83 | -- | -- |
| E7 | Success | AAV9 | 1.15E+08 | -- | -- | 3.92 | -- | -- |
| E8 | Success | AAV9 | 2.3E+08 | -- | -- | 5.61 | -- | -- |
| E9 | Success | AAV9 | 4.6E+08 | -- | -- | 7.03 | -- | -- |
| E10 | Success | AAV9 | 9.2E+08 | -- | -- | 8.7 | -- | -- |
| E11 | Success | AAV9 | 1.84E+09 | -- | -- | 10.3 | -- | -- |
| E12 | Success | AAV9 | 3.68E+09 | -- | -- | 11.5 | -- | -- |
| F1 | Success | AAV9 | 0 | -- | -- | 0.683 | -- | -- |
| F2 | Success | AAV9 | 3.59E+06 | -- | -- | 0.864 | -- | -- |
| F3 | Success | AAV9 | 7.19E+06 | -- | -- | 1.63 | -- | -- |
| F4 | Success | AAV9 | 1.44E+07 | -- | -- | 1.77 | -- | -- |
| F5 | Success | AAV9 | 2.88E+07 | -- | -- | 2.47 | -- | -- |
| F6 | Success | AAV9 | 5.75E+07 | -- | -- | 2.87 | -- | -- |

| Sensor | Status | Design | | | | Result | | |
|---|---|---|---|---|---|---|---|---|
| | | Target | Concentration | Sample ID | Dilution Factor | Nano Units | Measured Conc. | Undiluted Conc. |
| F7 | Success | AAV9 | 1.15E+08 | -- | -- | 3.72 | -- | -- |
| F8 | Success | AAV9 | 2.3E+08 | -- | -- | 5.33 | -- | -- |
| F9 | Success | AAV9 | 4.6E+08 | -- | -- | 6.75 | -- | -- |
| F10 | Success | AAV9 | 9.2E+08 | -- | -- | 9.46 | -- | -- |
| F11 | Success | AAV9 | 1.84E+09 | -- | -- | 9.92 | -- | -- |
| F12 | Success | AAV9 | 3.68E+09 | -- | -- | 11.2 | -- | -- |
| G7 | Success | AAV9 | -- | VB 4.5 kb | 3.2E+05 | 3.88 | 1.24E+08 | 3.98E+13 |
| G8 | Success | AAV9 | -- | VB 4.5 kb | 1.6E+05 | 5.42 | 2.63E+08 | 4.21E+13 |
| G9 | Success | AAV9 | -- | VB 4.5 kb | 8E+04 | 7.42 | 5.79E+08 | 4.63E+13 |
| G10 | Success | AAV9 | -- | VB 4.5 kb | 4E+04 | 8.59 | 8.95E+08 | 3.58E+13 |
| G11 | Success | AAV9 | -- | VB 4.5 kb | 2E+04 | 10.4 | 1.91E+09 | 3.82E+13 |
| G12 | Success | AAV9 | -- | VB 4.5 kb | 1E+04 | 10.9 | 2.41E+09 | 2.41E+13 |
| H7 | Success | AAV9 | -- | VB 4.5 kb | 3.2E+05 | 3.95 | 1.29E+08 | 4.14E+13 |
| H8 | Success | AAV9 | -- | VB 4.5 kb | 1.6E+05 | 5.1 | 2.28E+08 | 3.66E+13 |
| H9 | Success | AAV9 | -- | VB 4.5 kb | 8E+04 | 6.8 | 4.58E+08 | 3.66E+13 |
| H10 | Success | AAV9 | -- | VB 4.5 kb | 4E+04 | 8.94 | 1.02E+09 | 4.09E+13 |
| H11 | Success | AAV9 | -- | VB 4.5 kb | 2E+04 | 10.5 | 1.97E+09 | 3.94E+13 |
| H12 | Success | AAV9 | -- | VB 4.5 kb | 1E+04 | 11.4 | 3.32E+09 | 3.32E+13 |

### User Defined Values

| Metric | Target | Value | Units |
|---|---|---|---|
| Limits | AAV9 | 3 - 10.5 | Nano Units |
| CV Filter Threshold | -- | 20.0 | % |

### Ignored Wells

| | 1 | 2 | 3 | 4 | 5 | 6 | 7 | 8 | 9 | 10 | 11 | 12 |
|---|---|---|---|---|---|---|---|---|---|---|---|---|
| A | | | | | | | | | | | | |
| B | | | | | | | | | | | | |
| C | C1 | C2 | C3 | C4 | C5 | C6 | | | | | | |
| D | D1 | D2 | D3 | D4 | D5 | D6 | | | | | | |
| E | | | | | | | | | | | | |
| F | | | | | | | | | | | | |
| G | G1 | G2 | G3 | G4 | G5 | G6 | | | | | | |
| H | H1 | H2 | H3 | H4 | H5 | H6 | | | | | | |
