## Supplementary File S3 for "Nanoneedle-Enabled Quantification of rAAV9 Capsid and Genome Integrity Reveals a Truncation Hotspot Locus in a 4.5 kb Transgene": 20250603_2-43_2-49_titer.pdf

### Tessie Analysis Report

#### 96 well, 1 plex

| | |
|---|---|
| Plate Type: | AA01 |
| Assay Design: | 20250606_7168.2-43_2-49 |
| Assay Design Date: | 2025-10-27 12:30:25-04:00 |
| Plate ID: | 00007168 |
| Pre Scan Package: | AA0100007168_PRE_1 |
| Post Scan Package: | AA0100007168_POST_1 |
| Instrument ID: | T1039 |
| Pre Scan User: | N/A |
| Pre Scan Number: | 1 |
| Pre Scan Date: | 2025-06-02 17:27:46-04:00 |
| Post Scan User: | N/A |
| Post Scan Number: | 1 |
| Post Scan Date: | 2025-06-03 15:27:13-04:00 |
| Image Analysis Version: | 1.1.1.0 |
| Report Generation Date: | 2025-10-27 12:47:33-04:00 |
| Targets: | VB 2-43 2-49 |

#### NanoUnits Heatmap

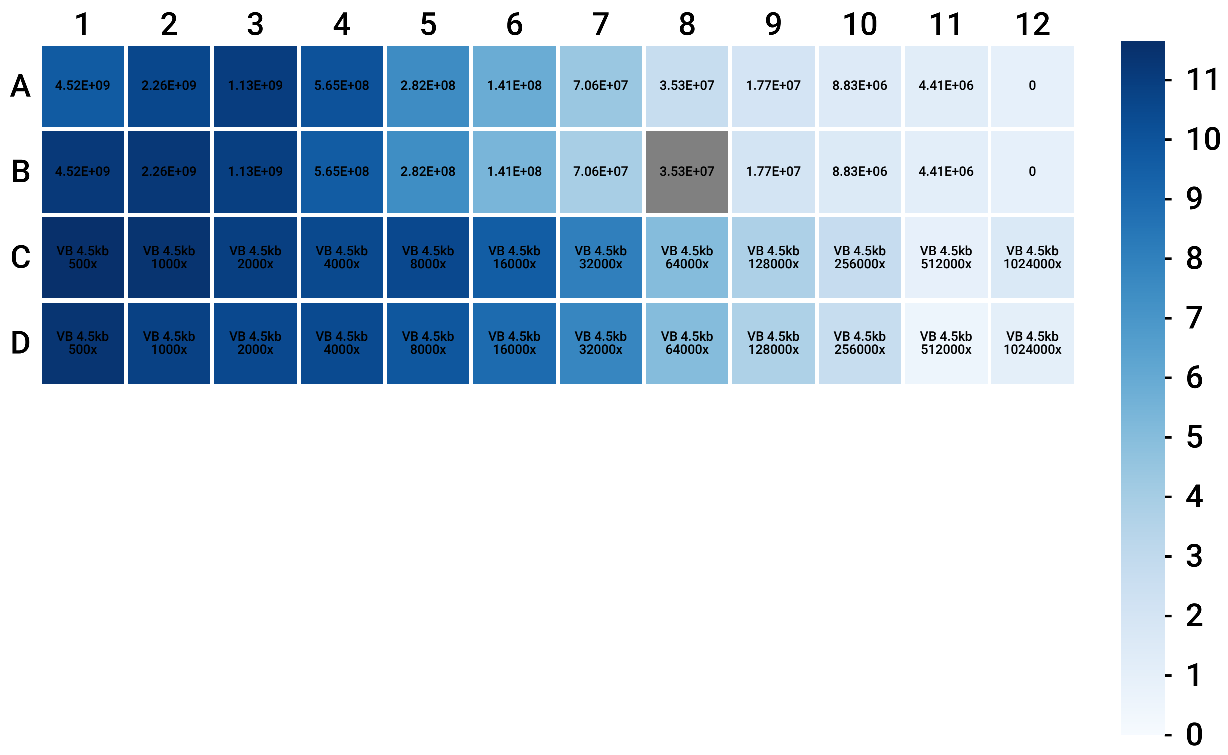

### VB 2-43 2-49 Standard Curve

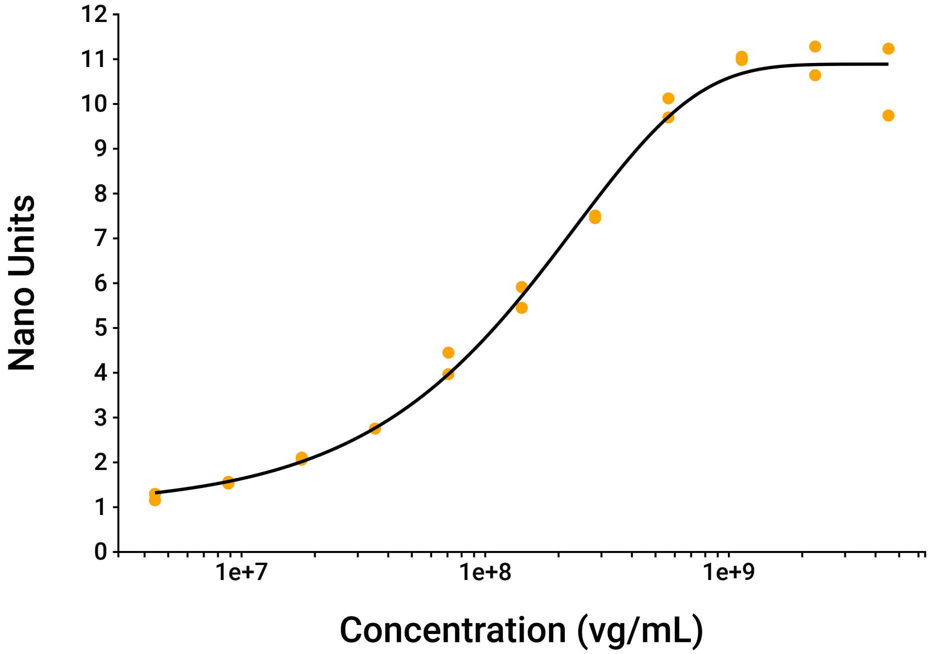


### VB 2-43 2-49 Standard Data

| Concentration (vg/mL) | Nano Units | | |
|---|---|---|---|
| | Mean | Standard Deviation | %CV |
| 0 | 0.989 | 0.0113 | 1.15 |
| 4.41E+06 | 1.22 | 0.102 | 8.33 |
| 8.83E+06 | 1.55 | 0.0266 | 1.72 |
| 1.77E+07 | 2.08 | 0.0325 | 1.56 |
| 3.53E+07 | 2.75 | -- | -- |
| 7.06E+07 | 4.21 | 0.34 | 8.09 |
| 1.41E+08 | 5.68 | 0.329 | 5.78 |
| 2.82E+08 | 7.48 | 0.0412 | 0.55 |
| 5.65E+08 | 9.91 | 0.3 | 3.03 |
| 1.13E+09 | 11 | 0.052 | 0.47 |
| 2.26E+09 | 11 | 0.453 | 4.13 |
| 4.52E+09 | 10.5 | 1.05 | 10.1 |

### VB 2-43 2-49 Sample Data

| Sample Label | Concentration (vg/mL) | | |
|---|---|---|---|
| | Mean | Standard Deviation | %CV |
| VB 4.5kb‡§ | 8.47E+12‡§ | 9.44E+11‡§ | 11.1‡§ |

‡One or more dilutions excluded due to concentration being below lower limit

§One or more dilutions excluded due to concentration being above upper limit

### VB 4.5kb Dilutions Data

| Dilution Factor | Measured Concentration (vg/mL) | Undiluted Concentration (vg/mL) | %CV | Linearity %CV |
|---|---|---|---|---|
| 32000 | 2.95E+08 | 9.42E+12 | 7.11 | 11.5 |
| 64000 | 1.13E+08 | 7.22E+12 | 0.04 | |
| 128000 | 6.43E+07 | 8.23E+12 | 1.49 | |
| 256000 | 3.51E+07 | 9E+12 | 4.07 | |

### Curve Parameters

| Target | Curve | Equation | Parameters | $R^2$ |
|---|---|---|---|---|
| VB 2-43 2-49 | Five parameter logisitic curve | $y = D - \frac{(A-D)}{\left[1+(x/C)^B\right]^E}$ | A = 0.999<br>B = 0.859<br>C = 9.7E+14<br>D = 10.9<br>E = 4.83E+05 | 0.993 |

### Raw Data

| Sensor | Status | Design | | | | Result | | |
|---|---|---|---|---|---|---|---|---|
| | | Target | Concentration | Sample ID | Dilution Factor | Nano Units | Measured Conc. | Undiluted Conc. |
| A1 | Success | VB 2-43 2-49 | 4.52E+09 | -- | -- | 9.74 | -- | -- |
| A2 | Success | VB 2-43 2-49 | 2.26E+09 | -- | -- | 10.6 | -- | -- |
| A3 | Success | VB 2-43 2-49 | 1.13E+09 | -- | -- | 11.1 | -- | -- |
| A4 | Success | VB 2-43 2-49 | 5.65E+08 | -- | -- | 10.1 | -- | -- |
| A5 | Success | VB 2-43 2-49 | 2.82E+08 | -- | -- | 7.51 | -- | -- |
| A6 | Success | VB 2-43 2-49 | 1.41E+08 | -- | -- | 5.91 | -- | -- |
| A7 | Success | VB 2-43 2-49 | 7.06E+07 | -- | -- | 4.45 | -- | -- |
| A8 | Success | VB 2-43 2-49 | 3.53E+07 | -- | -- | 2.75 | -- | -- |
| A9 | Success | VB 2-43 2-49 | 1.77E+07 | -- | -- | 2.06 | -- | -- |
| A10 | Success | VB 2-43 2-49 | 8.83E+06 | -- | -- | 1.53 | -- | -- |
| A11 | Success | VB 2-43 2-49 | 4.41E+06 | -- | -- | 1.3 | -- | -- |
| A12 | Success | VB 2-43 2-49 | 0 | -- | -- | 0.981 | -- | -- |
| B1 | Success | VB 2-43 2-49 | 4.52E+09 | -- | -- | 11.2 | -- | -- |
| B2 | Success | VB 2-43 2-49 | 2.26E+09 | -- | -- | 11.3 | -- | -- |
| B3 | Success | VB 2-43 2-49 | 1.13E+09 | -- | -- | 11 | -- | -- |
| B4 | Success | VB 2-43 2-49 | 5.65E+08 | -- | -- | 9.7 | -- | -- |
| B5 | Success | VB 2-43 2-49 | 2.82E+08 | -- | -- | 7.45 | -- | -- |

| Sensor | Status | Design | | | | Result | | |
|---|---|---|---|---|---|---|---|---|
| | | Target | Concentration | Sample ID | Dilution Factor | Nano Units | Measured Conc. | Undiluted Conc. |
| B6 | Success | VB 2-43 2-49 | 1.41E+08 | -- | -- | 5.45 | -- | -- |
| B7 | Success | VB 2-43 2-49 | 7.06E+07 | -- | -- | 3.97 | -- | -- |
| B8 | Excluded | VB 2-43 2-49 | 3.53E+07 | -- | -- | 1.48 | -- | -- |
| B9 | Success | VB 2-43 2-49 | 1.77E+07 | -- | -- | 2.11 | -- | -- |
| B10 | Success | VB 2-43 2-49 | 8.83E+06 | -- | -- | 1.57 | -- | -- |
| B11 | Success | VB 2-43 2-49 | 4.41E+06 | -- | -- | 1.15 | -- | -- |
| B12 | Success | VB 2-43 2-49 | 0 | -- | -- | 0.997 | -- | -- |
| C1 | Success | VB 2-43 2-49 | -- | VB 4.5kb | 500 | 11.6 | HIGH | -- |
| C2 | Success | VB 2-43 2-49 | -- | VB 4.5kb | 1E+03 | 11.5 | HIGH | -- |
| C3 | Success | VB 2-43 2-49 | -- | VB 4.5kb | 2E+03 | 11 | HIGH | -- |
| C4 | Success | VB 2-43 2-49 | -- | VB 4.5kb | 4E+03 | 10.5 | 9.63E+08 | 3.85E+12 |
| C5 | Success | VB 2-43 2-49 | -- | VB 4.5kb | 8E+03 | 10.6 | 9.81E+08 | 7.85E+12 |
| C6 | Success | VB 2-43 2-49 | -- | VB 4.5kb | 1.6E+04 | 9.65 | 5.49E+08 | 8.79E+12 |
| C7 | Success | VB 2-43 2-49 | -- | VB 4.5kb | 3.2E+04 | 8.11 | 3.09E+08 | 9.9E+12 |
| C8 | Success | VB 2-43 2-49 | -- | VB 4.5kb | 6.4E+04 | 5.09 | 1.13E+08 | 7.21E+12 |
| C9 | Success | VB 2-43 2-49 | -- | VB 4.5kb | 1.28E+05 | 3.8 | 6.5E+07 | 8.32E+12 |
| C10 | Success | VB 2-43 2-49 | -- | VB 4.5kb | 2.56E+05 | 2.8 | 3.62E+07 | 9.26E+12 |
| C11 | Success | VB 2-43 2-49 | -- | VB 4.5kb | 5.12E+05 | 0.959 | LOW | -- |

| Sensor | Status | Design | | | | Result | | |
|---|---|---|---|---|---|---|---|---|
| | | Target | Concentration | Sample ID | Dilution Factor | Nano Units | Measured Conc. | Undiluted Conc. |
| C12 | Success | VB 2-43 2-49 | -- | VB 4.5kb | 1.02E+06 | 1.68 | 1.09E+07 | 1.11E+13 |
| D1 | Success | VB 2-43 2-49 | -- | VB 4.5kb | 500 | 11.4 | HIGH | -- |
| D2 | Success | VB 2-43 2-49 | -- | VB 4.5kb | 1E+03 | 10.9 | 2.18E+09 | 2.18E+12 |
| D3 | Success | VB 2-43 2-49 | -- | VB 4.5kb | 2E+03 | 10.6 | 9.86E+08 | 1.97E+12 |
| D4 | Success | VB 2-43 2-49 | -- | VB 4.5kb | 4E+03 | 10.5 | 9.13E+08 | 3.65E+12 |
| D5 | Success | VB 2-43 2-49 | -- | VB 4.5kb | 8E+03 | 9.92 | 6.26E+08 | 5.01E+12 |
| D6 | Success | VB 2-43 2-49 | -- | VB 4.5kb | 1.6E+04 | 9.03 | 4.27E+08 | 6.82E+12 |
| D7 | Success | VB 2-43 2-49 | -- | VB 4.5kb | 3.2E+04 | 7.8 | 2.8E+08 | 8.95E+12 |
| D8 | Success | VB 2-43 2-49 | -- | VB 4.5kb | 6.4E+04 | 5.09 | 1.13E+08 | 7.22E+12 |
| D9 | Success | VB 2-43 2-49 | -- | VB 4.5kb | 1.28E+05 | 3.75 | 6.36E+07 | 8.14E+12 |
| D10 | Success | VB 2-43 2-49 | -- | VB 4.5kb | 2.56E+05 | 2.72 | 3.41E+07 | 8.74E+12 |
| D11 | Success | VB 2-43 2-49 | -- | VB 4.5kb | 5.12E+05 | 0.572 | LOW | -- |
| D12 | Success | VB 2-43 2-49 | -- | VB 4.5kb | 1.02E+06 | 1.09 | LOW | -- |

### User Defined Values

| Metric | Target | Value | Units |
|---|---|---|---|
| Limits | VB 2-43 2-49 | 2.5 - 9 | Nano Units |
| CV Filter Threshold | -- | 20.0 | % |

### Ignored Wells

| | 1 | 2 | 3 | 4 | 5 | 6 | 7 | 8 | 9 | 10 | 11 | 12 |
|---|---|---|---|---|---|---|---|---|---|---|---|---|
| A | | | | | | | | | | | | |
| B | | | | | | | | | | | | |
| C | | | | | | | | | | | | |
| D | | | | | | | | | | | | |
| E | E1 | E2 | E3 | E4 | E5 | E6 | E7 | E8 | E9 | E10 | E11 | E12 |
| F | F1 | F2 | F3 | F4 | F5 | F6 | F7 | F8 | F9 | F10 | F11 | F12 |
| G | G1 | G2 | G3 | G4 | G5 | G6 | G7 | G8 | G9 | G10 | G11 | G12 |
| H | H1 | H2 | H3 | H4 | H5 | H6 | H7 | H8 | H9 | H10 | H11 | H12 |
