## Supplementary File S4 for "Nanoneedle-Enabled Quantification of rAAV9 Capsid and Genome Integrity Reveals a Truncation Hotspot Locus in a 4.5 kb Transgene"

### *AAV Report*

*Assigned Types By Read Alignment Characteristics, overview*

| Assigned Type | Count | Frequency (%) |
|---|---|---|
| ssAAV | 85178 | 53.71 |
| scAAV | 55375 | 34.92 |
| other | 8793 | 5.54 |
| unmapped | 8432 | 5.32 |
| host | 799 | 0.50 |
| chimeric | 13 | 0.01 |

| Assigned Type | Count | Frequency (%) |
|---|---|---|
| ssAAV | 85178 | 60.6 |
| scAAV | 55375 | 39.4 |

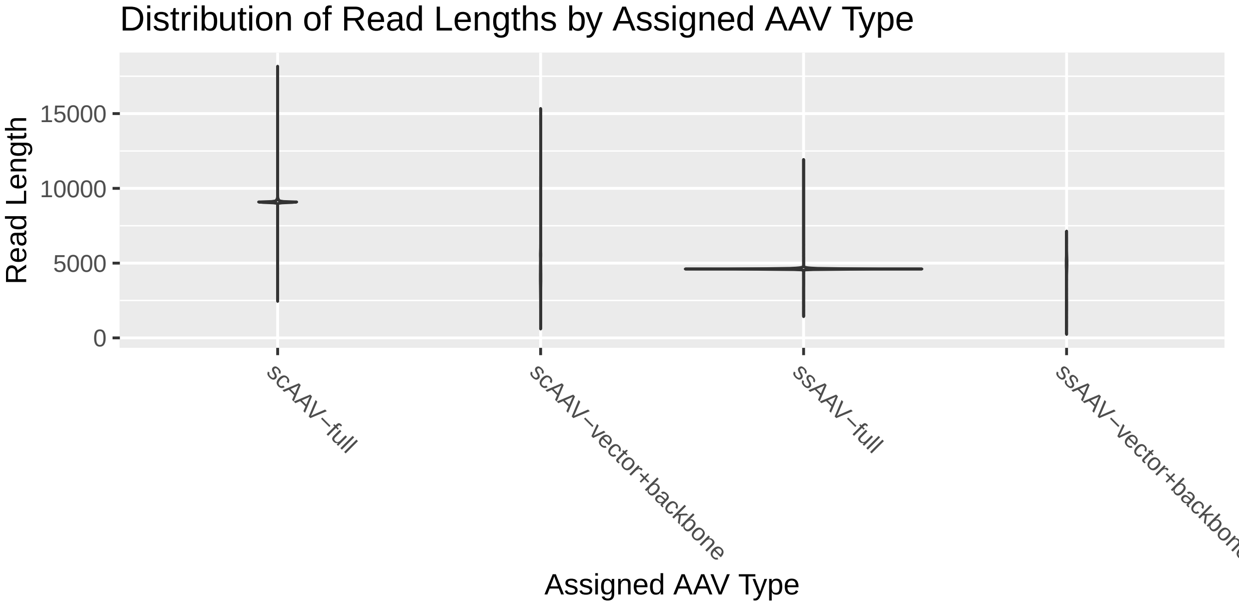

#### *Assigned AAV Types, detailed (top 20 only)*

| Assigned Type, detailed | Assigned Subtype | Count | Frequency (%) |
|---|---|---|---|
| ssAAV | full | 77927 | 55.44 |
| scAAV | left−partial | 22358 | 15.91 |
| scAAV | right−partial | 14917 | 10.61 |
| scAAV | full | 4937 | 3.51 |
| scAAV | backbone | 3321 | 2.36 |
| ssAAV | right−partial | 3076 | 2.19 |
| scAAV | vector+backbone | 2754 | 1.96 |
| scAAV | partial | 1998 | 1.42 |
| scAAV | vector+backbone\|right−partial | 1972 | 1.40 |
| ssAAV | partial | 1512 | 1.08 |
| ssAAV | left−partial | 1473 | 1.05 |
| scAAV | full\|backbone | 928 | 0.66 |
| ssAAV | backbone | 896 | 0.64 |
| scAAV | full\|right−partial | 453 | 0.32 |
| scAAV | backbone\|left−partial | 387 | 0.28 |
| scAAV | full\|left−partial | 331 | 0.24 |
| ssAAV | vector+backbone | 294 | 0.21 |
| scAAV | right−partial\|left−partial | 216 | 0.15 |
| scAAV | backbone\|right−partial | 156 | 0.11 |
| scAAV | full\|partial | 144 | 0.10 |

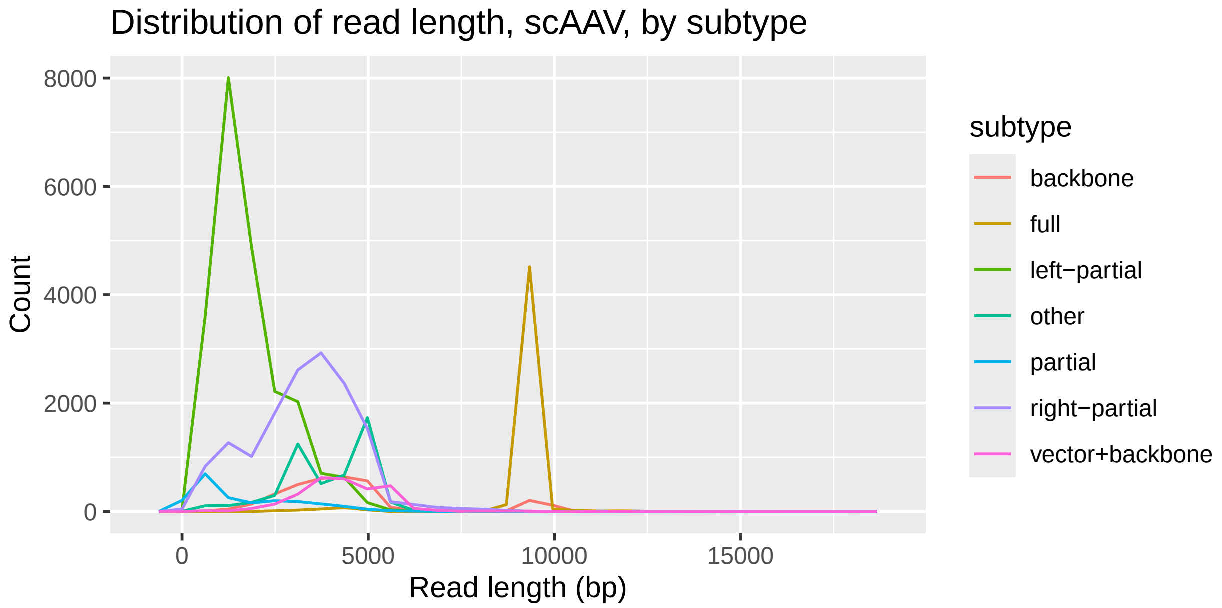

Distribution of read length, scAAV, by subtype
Count
8000
6000
4000
2000
0
0
5000
10000
15000
Read length (bp)
subtype
backbone
full
left−partial
other
partial
right−partial
vector+backbone


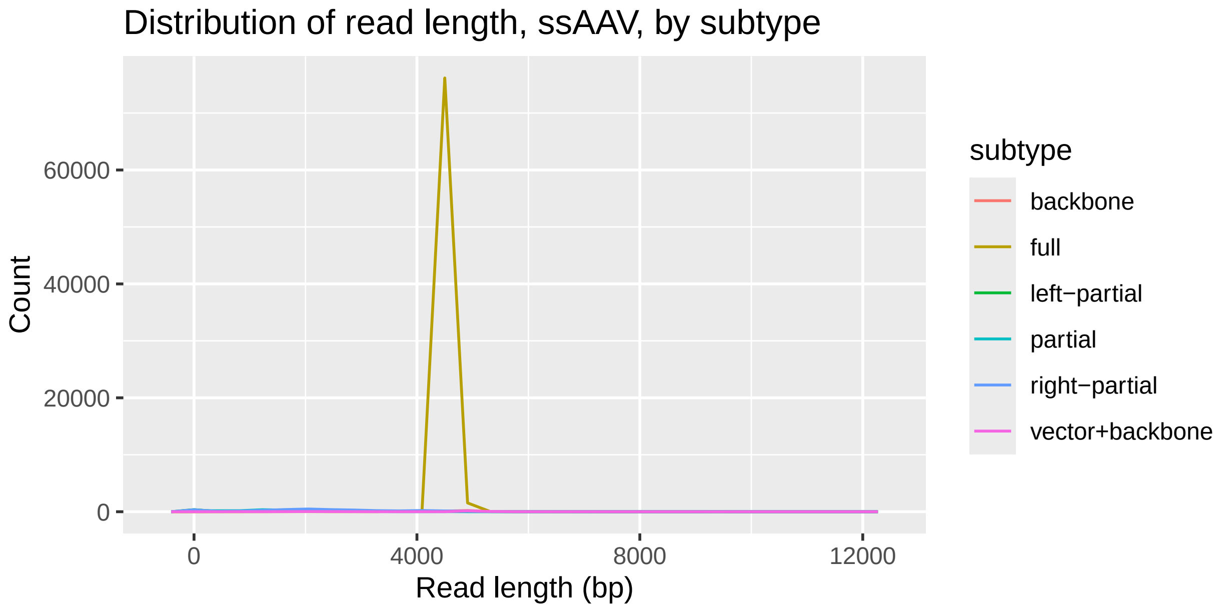

Distribution of read length, ssAAV, by subtype
Count
60000
40000
20000
0
0
4000
8000
12000
Read length (bp)
subtype
backbone
full
left−partial
partial
right−partial
vector+backbone

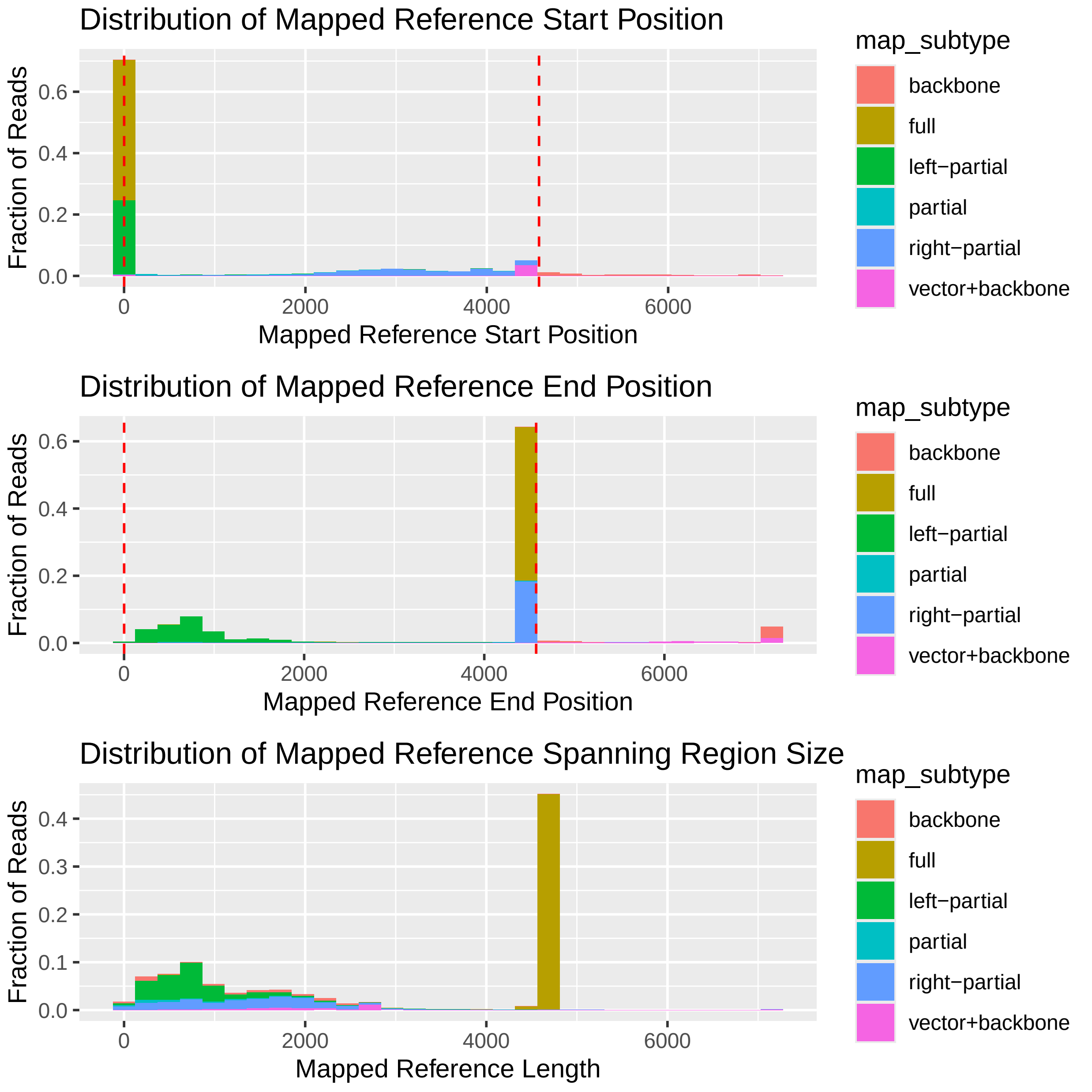
Distribution of Mapped Reference Start Position
Fraction of Reads
0.0
0.2
0.4
0.6
0
2000
4000
6000
Mapped Reference Start Position
map_subtype
backbone
full
left−partial
partial
right−partial
vector+backbone
Distribution of Mapped Reference End Position
Fraction of Reads
0.0
0.2
0.4
0.6
0
2000
4000
6000
Mapped Reference End Position
map_subtype
backbone
full
left−partial
partial
right−partial
vector+backbone
Distribution of Mapped Reference Spanning Region Size
Fraction of Reads
0.0
0.1
0.2
0.3
0.4
0
2000
4000
6000
Mapped Reference Length
map_subtype
backbone
full
left−partial
partial
right−partial
vector+backbone

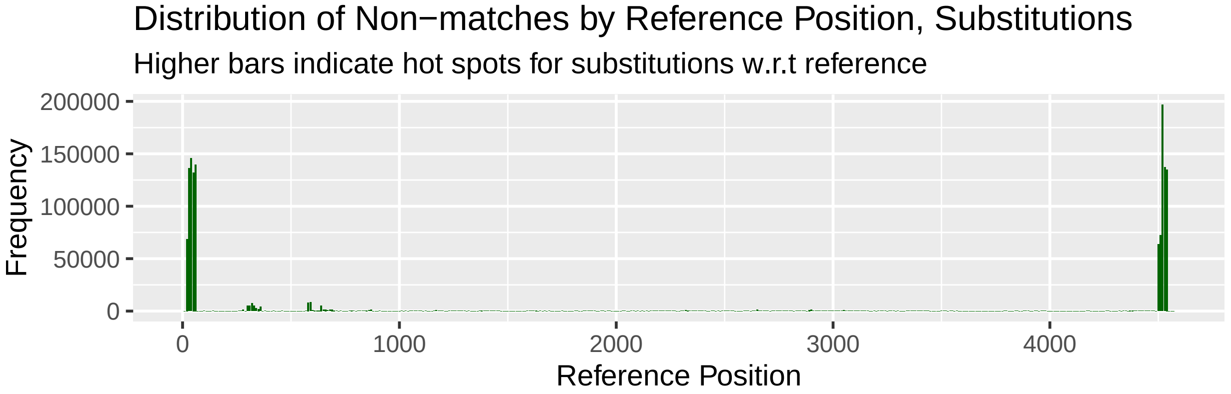
Distribution of Non−matches by Reference Position, Substitutions
Higher bars indicate hot spots for substitutions w.r.t reference
Frequency
200000
150000
100000
50000
0
0
1000
2000
3000
4000
Reference Position

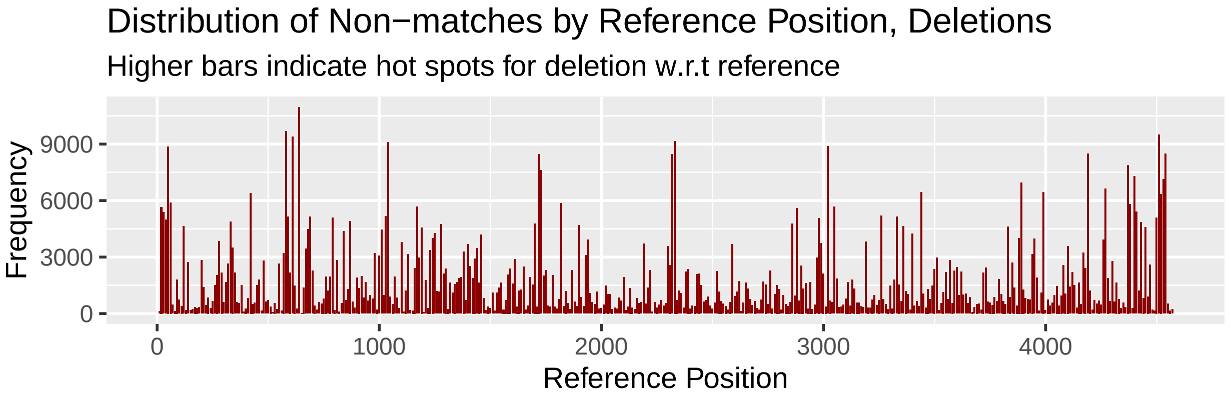
Distribution of Non−matches by Reference Position, Deletions
Higher bars indicate hot spots for deletion w.r.t reference
Frequency
9000
6000
3000
0
0
1000
2000
3000
4000
Reference Position

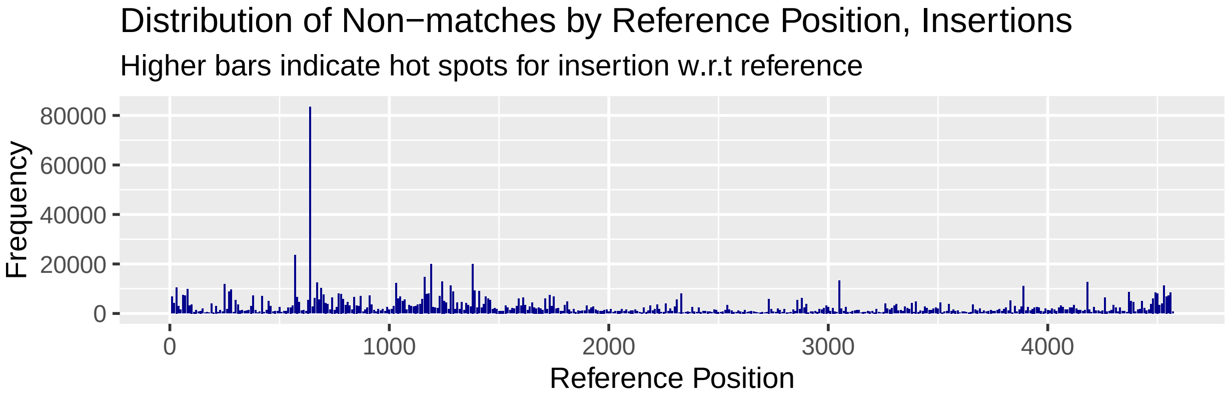
Distribution of Non−matches by Reference Position, Insertions
Higher bars indicate hot spots for insertion w.r.t reference
Frequency
80000
60000
40000
20000
0
0
1000
2000
3000
4000
Reference Position

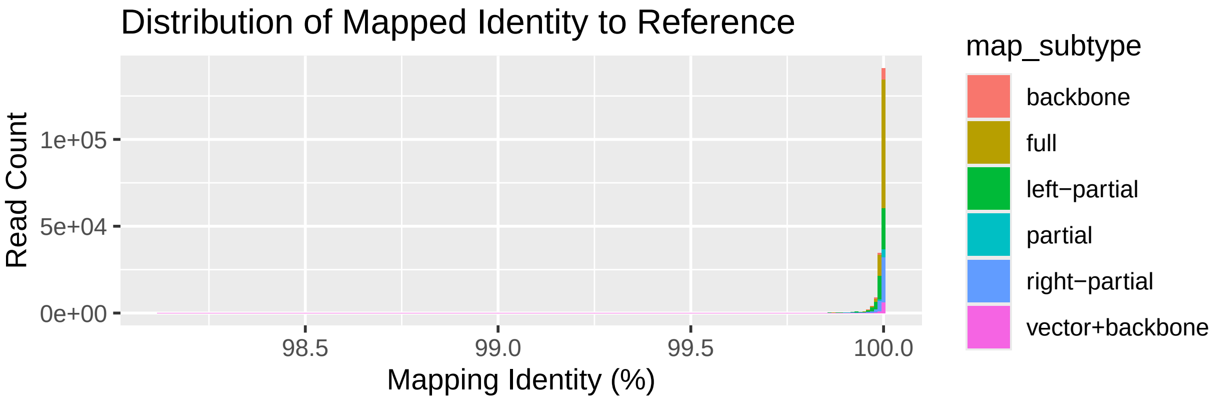
Distribution of Mapped Identity to Reference
Read Count
1e+05
5e+04
0e+00
98.5
99.0
99.5
100.0
Mapping Identity (%)
map_subtype
backbone
full
left−partial
partial
right−partial
vector+backbone

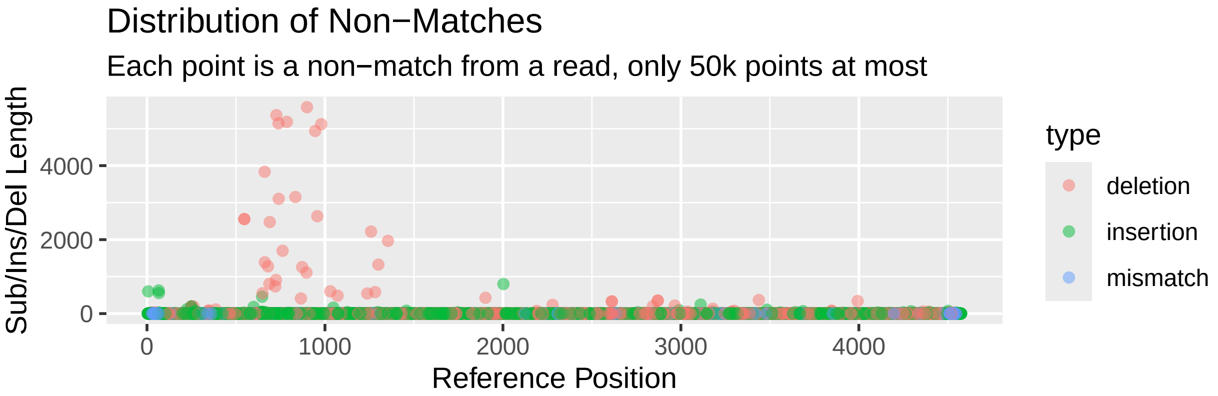
Distribution of Non−Matches
Each point is a non−match from a read, only 50k points at most
Sub/Ins/Del Length
4000
2000
0
0
1000
2000
3000
4000
Reference Position
type
deletion
insertion
mismatch

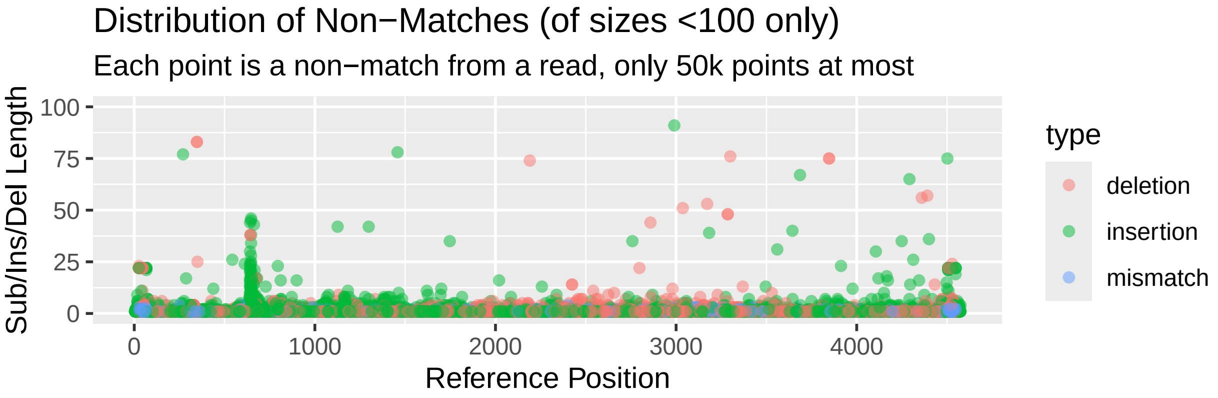
Distribution of Non−Matches (of sizes <100 only)
Each point is a non−match from a read, only 50k points at most
Sub/Ins/Del Length
100
75
50
25
0
0
1000
2000
3000
4000
Reference Position
type
deletion
insertion
mismatch

*Length Distribution of Different Non−matches*

| Err Type | Err Length | Count | Frequency (%) |
|---|---|---|---|
| deletion | 1−10 | 869132 | 23.68 |
| deletion | 11−100 | 4376 | 0.12 |
| deletion | 100−500 | 2274 | 0.06 |
| deletion | >500 | 1287 | 0.04 |
| insertion | 1−10 | 1386948 | 37.79 |
| insertion | 11−100 | 6954 | 0.19 |
| insertion | 100−500 | 371 | 0.01 |
| insertion | >500 | 154 | 0.00 |
| mismatch | 1−10 | 1399116 | 38.12 |
| mismatch | 11−100 | 10 | 0.00 |
